# Raptor downregulation rescues neuronal phenotypes in mouse models of Tuberous Sclerosis Complex

**DOI:** 10.1101/2021.07.21.453275

**Authors:** Vasiliki Karalis, Franklin Caval-Holme, Helen S. Bateup

## Abstract

Tuberous Sclerosis Complex (TSC) is a neurodevelopmental disorder caused by mutations in the *TSC1* or *TSC2* genes, which encode proteins that negatively regulate mTOR complex 1 (mTORC1) signaling. Current treatment strategies focus on mTOR inhibition with rapamycin and its derivatives. While effective at improving some aspects of TSC, chronic rapamycin inhibits both mTORC1 and mTORC2 and is associated with systemic side-effects. It is currently unknown which mTOR complex is most relevant for TSC-related brain phenotypes. Here we used genetic strategies to selectively reduce neuronal mTORC1 or mTORC2 activity in mouse models of TSC. We find that reduction of the mTORC1 component Raptor, but not the mTORC2 component Rictor, rebalanced mTOR signaling in Tsc1 knock-out neurons. Raptor reduction was sufficient to improve several TSC-related phenotypes including neuronal hypertrophy, macrocephaly, impaired myelination, network hyperactivity, and premature mortality. Raptor downregulation represents a promising potential therapeutic intervention for the neurological manifestations of TSC.

## Introduction

Tuberous Sclerosis Complex (TSC) is a neurodevelopmental disorder resulting in benign tumors in multiple organs and focal cortical malformations called tubers^1^. Some of the most significant problems associated with TSC are the neurological and psychiatric aspects^2^. Approximately 85% of TSC patients develop epilepsy, which often begins in infancy and becomes intractable in two-thirds of cases^3, 4^. TSC is also associated with varying degrees of intellectual disability, cognitive impairments, and behavioral conditions including autism spectrum disorder and attention deficit hyperactivity disorder^5^.

Current treatment strategies for TSC include inhibitors of mTOR signaling called rapalogs, which are analogs of the naturally occurring macrolide rapamycin^6, 7^. Rapamycin has been successful in treating neuropsychiatric phenotypes in rodent models of TSC, especially when treatment is started early in development^8, 9^. In the clinic, rapalogs are moderately effective at treating seizures and subependymal giant cell astrocytomas (SEGAs), which are benign brain tumors that affect about 5-15% of TSC patients^10–14^. However, symptoms can return after treatment cessation^10^ and chronic rapalog use is associated with significant side-effects including immunosuppression and systemic metabolic changes such as insulin resistance^15^. In addition, several recent clinical trials have reported that rapalogs did not significantly improve the cognitive and psychiatric aspects of TSC^16, 17^. As a result, additional therapeutic interventions are needed.

TSC is caused by loss-of-function mutations in either the *TSC1* or *TSC2* genes^18, 19^. At the biochemical level, the protein products of *TSC1* and *TSC2* form a complex that inhibits Rheb, a small GTPase that activates mTOR complex 1 (mTORC1)^20^. The Tsc2 protein harbors the GTPase activating protein (GAP) domain, while Tsc1 is required for complex stability^21^. mTORC1 is composed of several proteins including mTOR and the obligatory component Regulatory-associated protein of mTOR (Raptor)^22–24^. mTORC1 is a kinase that phosphorylates several targets including p70S6K and the 4E binding proteins (4E-BPs), which are involved in translational control^25–27^. P70S6K in turn phosphorylates ribosomal protein S6, a canonical read-out of mTORC1 activity^28^. The activity of mTORC1 is regulated by various intra- and extracellular signals including growth factors, insulin, nutrients, and neural activity^29, 30^. When active, mTORC1 promotes anabolic cellular processes such as protein, lipid and nucleotide synthesis and suppresses catabolic processes, including autophagy^30, 31^. In the absence of regulation by the TSC1/2 complex, mTORC1 is constitutively active^20^.

Rapamycin is thought to suppress mTORC1 signaling by inducing a trimeric complex between itself, mTOR and FK506-binding protein 12 (FKBP12)^22^. Binding of rapamycin-FKBP12 on mTORC1 occludes access of some substrates to the catalytic site of the mTOR kinase^22^. While acute administration of rapamycin selectively reduces mTORC1 signaling, chronic treatment also suppresses the activity of a second mTOR complex, mTORC2, potentially by preventing *de novo* mTORC2 assembly^32^. mTORC2 shares several protein components with mTORC1 but instead of Raptor, mTORC2 contains Rapamycin-insensitive companion of mTOR (Rictor) as an essential component^31, 33, 34^. mTORC2 has been reported to control aspects of cytoskeletal organization, cell survival and metabolism^30^, although its functions in neurons are not well understood. The most well characterized phosphorylation target of mTORC2 is Ser473 of Akt^35^. While mTORC1 and mTORC2 have conventionally been thought to have distinct upstream regulators and targets, studies in various systems have revealed potential points of crosstalk between the two complexes^36, 37^.

TSC-related phenotypes are canonically thought to be due to mTORC1 hyperactivity; however, recent studies have raised the possibility that mTORC2 may also be involved. In particular, a study in mice with forebrain-specific disruption of Pten, a negative regulator of mTORC1 that is upstream of Tsc1/2, showed that downregulation of mTORC2, but not mTORC1, could prevent behavioral abnormalities, seizures, and premature mortality^38^. Moreover, it was shown that mTORC2, but not mTORC1, is required for hippocampal mGluR-dependent long-term depression (LTD)^39^, a form of synaptic plasticity that is altered in mouse models of TSC^40–43^. Therefore, a careful investigation of the relationships between Tsc1/2, mTORC1, and mTORC2 in the context of brain development and function is needed to design the most effective therapeutic strategy for TSC.

Here we used *in vitro* and *in vivo* mouse models of TSC to investigate whether manipulation of mTORC1 or mTORC2 signaling via genetic reduction of Raptor or Rictor, respectively, could prevent TSC-related brain phenotypes. We find that reduction, but not complete elimination, of Raptor rebalances both mTORC1 and mTORC2 signaling in the context of Tsc1 loss. We show that heterozygous deletion of *Rptor* in conditional *Tsc1* knock-out mice (Tsc1-cKO) improves several phenotypes including neuronal hypertrophy, neural network hyperactivity, impaired myelination, altered cortical and hippocampal architecture, and premature mortality. By contrast, heterozygous or homozygous loss of *Rictor* does not ameliorate TSC-related neural phenotypes. Finally, we demonstrate that postnatal Raptor downregulation rescues neuronal hypertrophy and myelination deficits in Tsc1-cKO mice. Our results highlight the central role of mTORC1 in driving TSC-related brain phenotypes in mouse models and establish Raptor as a potential target for treating the neurological presentations of TSC.

## Results

### Chronic rapamycin treatment suppresses mTORC1 and mTORC2 signaling in Tsc1-cKO hippocampal cultures

To investigate how loss of Tsc1 affects mTORC1 and mTORC2 signaling, we generated primary hippocampal cultures from *Tsc1^fl/fl^* mice^44^ and treated them with adeno-associated virus (AAV) expressing GFP (control) or Cre (Tsc1-cKO) at two days *in vitro* (DIV 2) (Fig. 1a and Supplementary Fig. 1a,b). To assess mTORC1 signaling status we quantified Raptor protein levels and two canonical downstream phosphorylation targets, ribosomal protein S6 (p-S6 Ser240/244) and 4E-BP1 (p-4E-BP1 Thr37). To measure mTORC2 signaling we quantified Rictor protein levels and Akt phosphorylation at Ser 473 (p-Akt Ser473). We found that on DIV 14, Tsc1-cKO cultures had complete loss of Tsc1 and a small increase in Raptor protein suggesting potentially enhanced mTORC1 assembly (Fig. 1b-d). No significant changes in Rictor protein were observed (Fig. 1e). Phosphorylation of S6 was increased in Tsc1-cKO cultures, as expected (Fig. 1f). However, we did not observe significant changes in p-4E-BP1 in this system (Fig. 1g), which may be due to the timing of Tsc1 loss, as discussed further below. In terms of mTORC2, we found that *Tsc1* deletion decreased Akt phosphorylation at Ser473 (Fig. 1h), indicating reduced mTORC2 activity. Together these data show that in hippocampal cultures, loss of *Tsc1* has opposing effects on the two mTOR complexes: it increases mTORC1 signaling, particularly the p70S6K/S6 branch, and decreases mTORC2 activity.

**Figure 1.**
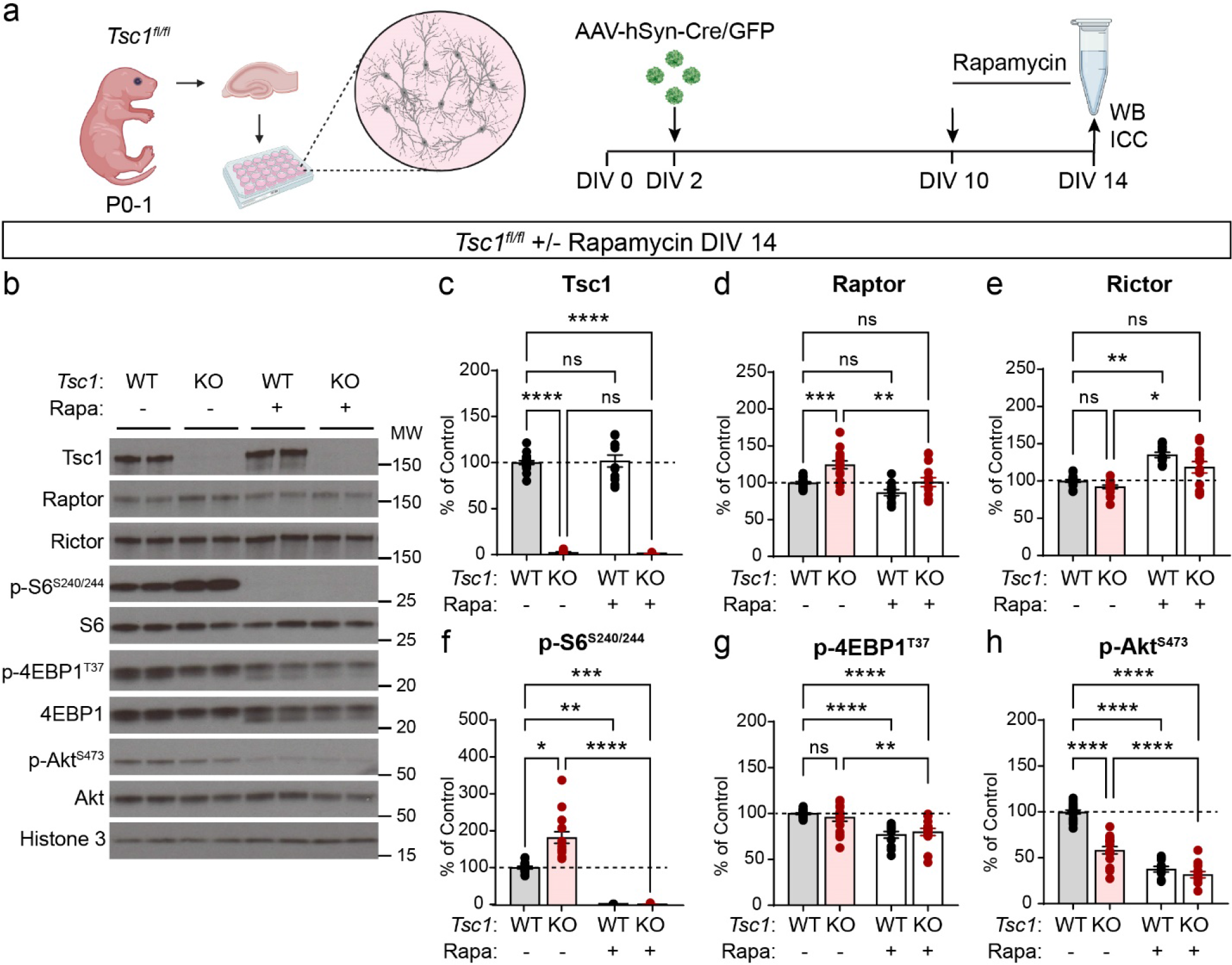
Chronic rapamycin suppresses mTORC1 and mTORC2 signaling in cultured Tsc1-cKO neurons. a) Schematic of the experiment. Primary hippocampal cultures were prepared from *Tsc1^fl/fl^* mice on postnatal day (P) 0-1. AAV-GFP or AAV-Cre-GFP was added on DIV 2 and rapamycin or vehicle was added on DIV 10. Cells were collected for analysis by western blot (WB) or immunocytochemistry (ICC) on DIV 14. Created with BioRender.com b) Representative western blots from *Tsc1^fl/fl^* cultures with (+) or without (-) four-day rapamycin (Rapa) treatment. MW indicates molecular weight. WT=*Tsc1^fl/fl^* + AAV-GFP; KO=*Tsc1^fl/fl^* + AAV-Cre-GFP. Two independent samples per genotype are shown; this experiment was replicated seven times. c-h) Bar graphs display western blot quantification (mean +/-SEM) for the indicated proteins, expressed as a percentage of Control (WT) levels. Phospho-proteins were normalized to their respective total proteins. Dots represent data from individual culture wells. WT n=18, KO n=15, WT+Rapa n=11 and KO+Rapa n=14 culture wells from 7 independent cultures; 2 mice per culture. ns=non-significant. Dashed lines at 100% indicate Control levels. c) Tsc1, Kruskal-Wallis, p<0.0001; WT vs KO, ****p<0.0001; WT vs WT+Rapa, p>0.9999; WT vs KO+Rapa, **** p<0.0001; KO vs KO+Rapa, p>0.9999; Dunn’s multiple comparisons tests. d) Raptor, One-way ANOVA, p<0.0001, F (3, 54) = 11.89; WT vs KO, ***p=0.0004; WT vs WT+Rapa, p=0.1541; WT vs KO+Rapa, p=0.9997; KO vs KO+Rapa, **p=0.0016; Sidak’s multiple comparisons tests. e) Rictor, Kruskal-Wallis, p<0.0001; WT vs KO p=0.3281; WT vs WT+Rapa, ** p=0.0023; WT vs KO+Rapa, p=0.7348; KO vs KO+Rapa, *p=0.0144, Dunn’s multiple comparisons tests. f) p-S6 Ser240/244, Kruskal-Wallis, p<0.0001; WT vs KO, *p=0.0310; WT vs WT+Rapa, ** p=0.0022; WT vs KO+Rapa, ***p=0.0008; KO vs KO+Rapa, ****p<0.0001; Dunn’s multiple comparisons tests. g) p-4E-BP1 T37, One-way ANOVA, p<0.0001, F (3, 54) = 12.40; WT vs KO, p=0.3293; WT vs WT+Rapa, ****p<0.0001; WT vs KO+Rapa, ****p<0.0001; KO vs KO+Rapa, **p=0.0021; Holm-Sidak’s multiple comparisons tests. h) p-Akt Ser473, One-way ANOVA, p<0.0001, F (3, 54) = 103.2; WT vs KO, ****p<0.0001; WT vs WT+Rapa, ****p<0.0001; WT vs KO+Rapa, ****p<0.0001; KO vs KO+Rapa, ****p<0.0001; Sidak’s multiple comparisons tests. See also Supplementary Figure 1.

We next examined the effects of rapamycin on neuronal mTOR signaling to provide a benchmark for comparing the effects of genetic manipulation of the two mTOR complexes. Four day treatment with 50 nM rapamycin had no effect on Tsc1 levels but reduced Raptor and modestly increased Rictor levels in Tsc1-cKO cultures (Fig. 1c-e). We observed the expected complete loss of p-S6 and partial reduction of p-4E-BP1 in both control and Tsc1-cKO cultures (Fig. 1f,g), indicative of suppressed mTORC1 signaling. Notably, we observed a strong reduction of p-Akt in both control and Tsc1-cKO neurons treated chronically with rapamycin that was greater than with Tsc1 loss alone (Fig. 1h). Together, these data demonstrate that rapamycin treatment does not restore balanced mTOR signaling in Tsc1-cKO hippocampal cultures but rather strongly suppresses the p70S6K branch of mTORC1, partially decreases p-4E-B1, and further reduces mTORC2-dependent Akt phosphorylation.

### Downregulation of Raptor, but not Rictor, improves mTOR signaling abnormalities in Tsc1-cKO neurons

To investigate whether genetic disruption of mTORC1 or mTORC2 could improve signaling abnormalities in Tsc1-cKO neurons, we crossed *Tsc1^fl/fl^* mice to mice with either floxed *Rptor*^45^ or *Rictor*^46, 47^ alleles for simultaneous Cre-dependent deletion of *Tsc1* and *Rptor* or *Tsc1* and *Rictor*, respectively (Fig. 2a,b and Supplementary Fig. 1c,d). We found that mTORC1 hyperactivity persisted in Tsc1-cKO cultures with deletion of *Rictor*, evidenced by significantly increased levels of p-S6 in Tsc1-cKO;Rictor-cKO cultures compared to controls (Fig. 2c,d). p-Akt-473 was strongly reduced in Tsc1-cKO;Rictor-cKO neurons (Fig. 2c,d), similar to Tsc1-cKO cells. Thus, the mTOR signaling perturbations in Tsc1-cKO hippocampal cultures were not improved by genetic reduction of Rictor.

**Figure 2.**
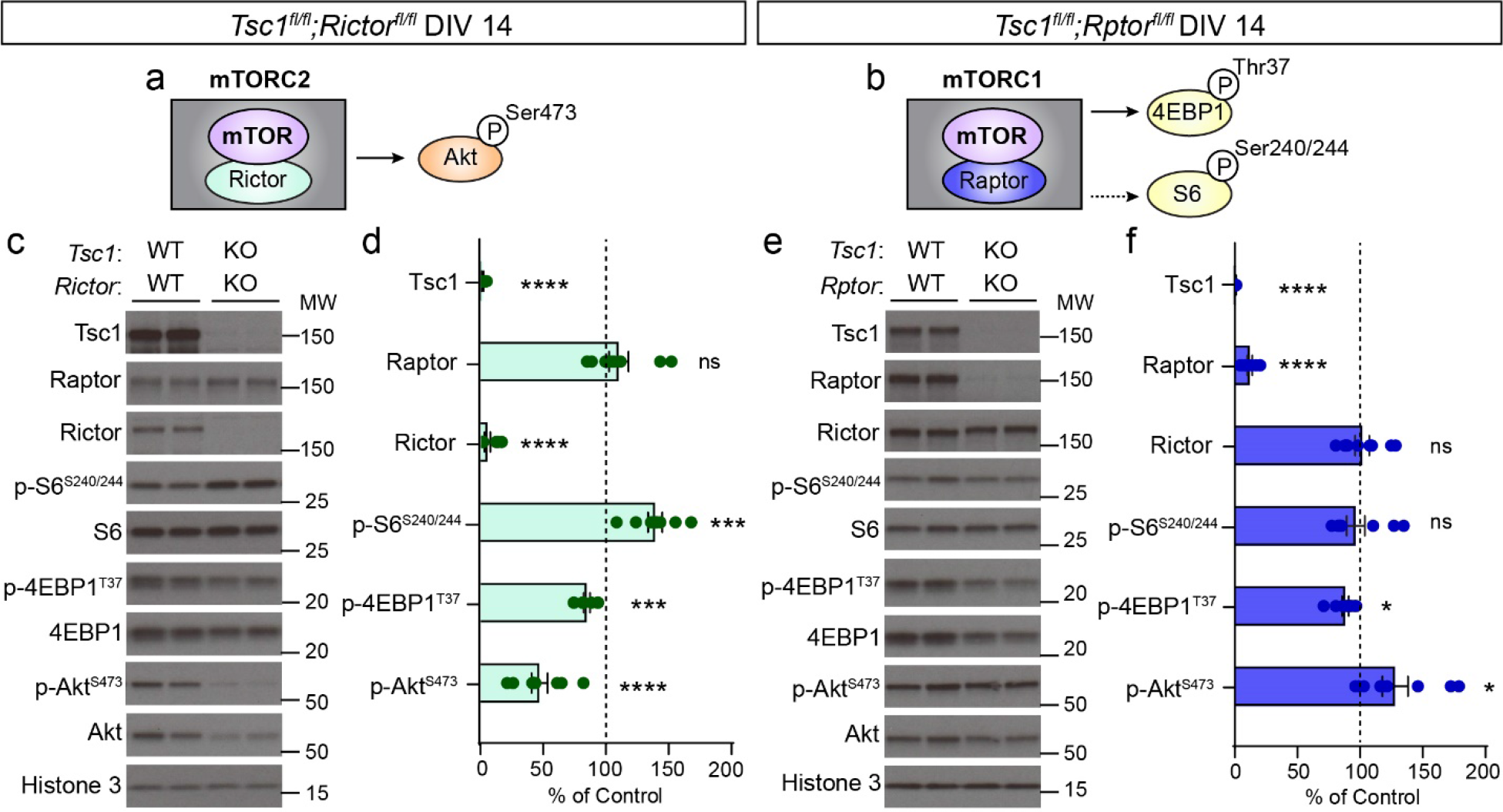
Genetic reduction of Raptor, but not Rictor, ameliorates mTOR signaling abnormalities in Tsc1-cKO neurons. a) Simplified schematic of mTORC2 showing mTOR and its obligatory binding partner Rictor. mTORC2 phosphorylates Ser473 on Akt. b) Simplified schematic of mTORC1 showing mTOR and its obligatory binding partner Raptor. mTORC1 phosphorylates 4E-BP1 (Thr37) and p70S6K, which in turn phosphorylates S6 at Ser240/244 (represented by the dashed line). c) Representative western blots from *Tsc1^fl/fl^;Rictor^fl/fl^* hippocampal cultures treated with AAV-GFP (WT;WT) or AAV-Cre-GFP (KO;KO) and harvested on DIV 14. MW indicates molecular weight. Two independent samples per genotype are shown; this experiment was replicated three times. d) Bar graphs display western blot quantification (mean +/-SEM) for the indicated proteins, expressed as a percentage of Control (WT) levels. n=9 culture wells from 3 independent cultures per genotype; 2 mice per culture. Tsc1 ****p<0.0001; Raptor p=0.2380; Rictor **** p<0.0001; p-S6 Ser240/244 *** p=0.0007; p-4EBP1 T37 *** p=0.0002; p-Akt Ser473 **** p<0.0001; Welch’s t-tests. e) Representative western blots of lysates collected from *Tsc1^fl/fl^;Rptor^fl/fl^* hippocampal cultures treated with AAV-GFP (WT;WT) or AAV-Cre-GFP (KO;KO) and harvested on DIV 14. Two independent samples per genotype are shown; this experiment was replicated three times. f) Bar graphs display western blot quantification (mean +/-SEM) for the indicated proteins, expressed as a percentage of Control (WT) levels. n=9 culture wells from 3 independent cultures per genotype; 2 mice per culture. Tsc1 ****p<0.0001; Raptor ****p<0.0001; Rictor p=0.8199; p-S6 Ser240/244 p=0.6616; p-4EBP1 T37 *p=0.0146; p-Akt Ser473 *p=0.0283; Welch’s t-tests. For panels d and f, phospho-proteins were normalized to their respective total proteins. Dots represent data from individual culture wells. Dashed lines at 100% indicate Control levels. ns=non-significant. See also Supplementary Figure. 1.

We next examined how reduction of Raptor affected mTOR signaling in Tsc1-cKO cultures. Homozygous deletion of *Rptor* effectively normalized p-S6 levels in Tsc1-cKO cultures harvested on DIV 14 (Fig. 2e,f). Deletion of *Rptor* also reversed the reduction of p-Akt caused by Tsc1 loss, suggesting a boosting of mTORC2 activity (Fig. 2e,f). The enhanced Akt phosphorylation resulting from Raptor loss may be due to relief of mTORC1-dependent negative feedback, either on mTORC2 itself or upstream regulators of both mTOR complexes^37^. These data indicate that Raptor downregulation could be an effective strategy for rebalancing mTOR signaling abnormalities due to loss of *Tsc1*.

During this experiment, we noted that low levels of Raptor protein remained in the cultures, which could have accounted for the incomplete suppression of mTORC1 signaling compared to rapamycin treatment (see Fig. 1 vs. Fig. 2e,f). To investigate this further, we added AAV-Cre to Tsc1-cKO;Rptor-cKO cultures on DIV 2 and harvested the cells at different time points. We found that while Tsc1 protein was virtually undetectable by DIV 14, residual Raptor protein remained. Raptor levels decreased over time such that by DIV 18 (16 days post-Cre) there was less than 1% of Raptor protein remaining (Supplementary Fig. 2a-g). This slow turn-over may reflect higher stability of Raptor protein bound within mTORC1. We therefore harvested Tsc1-cKO;Raptor-cKO cultures on day 18 to assess the consequences of more complete loss of Raptor on mTORC1 and mTORC2 signaling. At this time point, we found that concomitant deletion of *Tsc1* and *Rptor* strongly reduced p-S6 and increased p-Akt levels (Fig. 3a,b). Interestingly, we did not observe a consistent reduction in 4E-BP1 phosphorylation (Fig. 3b), suggesting that, as with rapamycin^48^, mTORC1 targets are differentially susceptible to Raptor downregulation.

**Figure 3.**
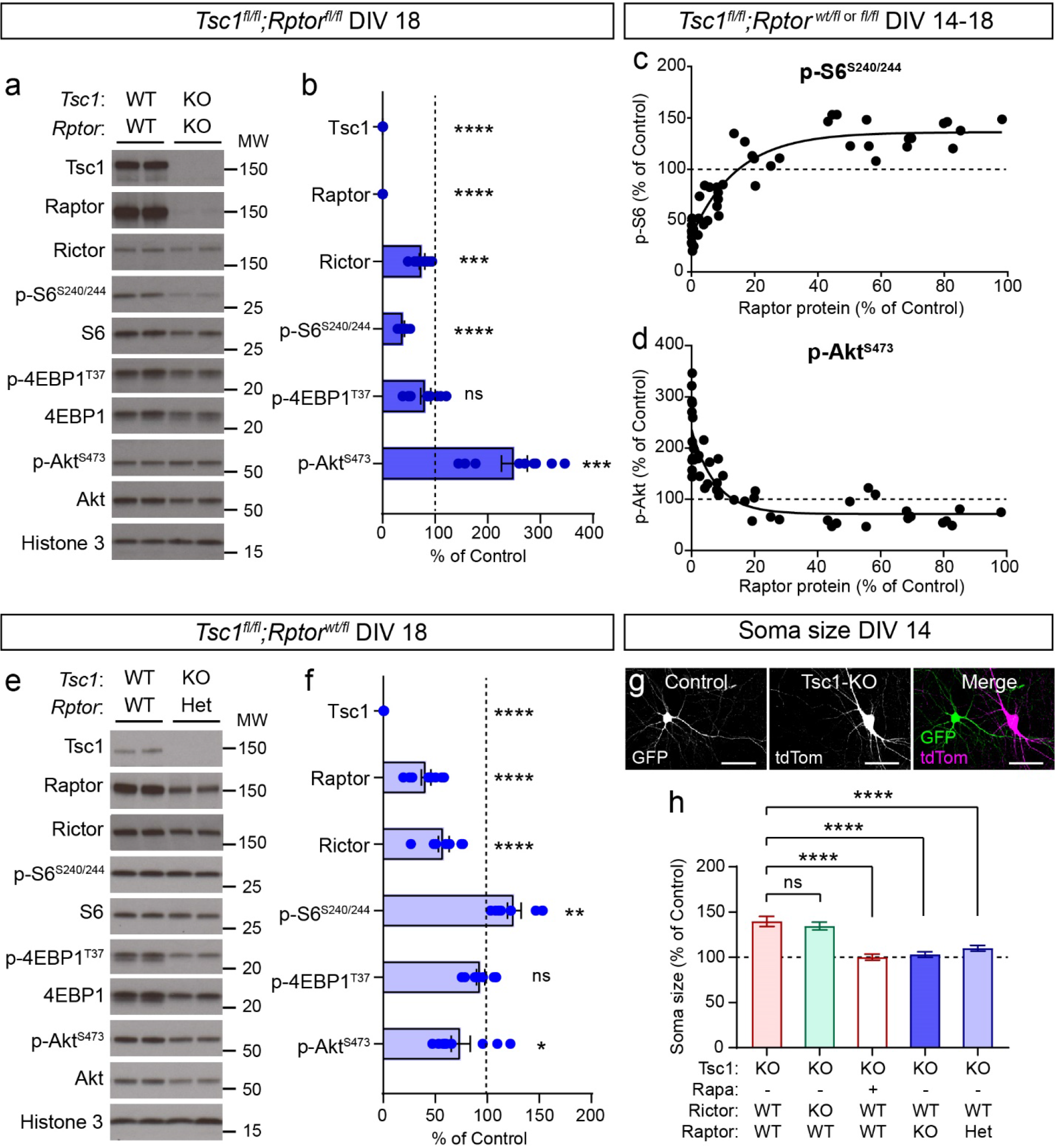
Partial reduction of Raptor normalizes mTOR signaling in Tsc1-cKO cultures. a) Representative western blots from *Tsc1^fl/fl^;Rptor ^fl/fl^* hippocampal cultures treated with AAV-GFP (WT;WT) or AAV-Cre-GFP (KO;KO) and harvested on DIV 18. MW indicates molecular weight. Two independent samples per genotype are shown; this experiment was replicated three times. b) Bar graphs display western blot quantification (mean +/-SEM) for the indicated proteins, expressed as a percentage of Control (WT) levels. n=9 culture wells from 3 independent cultures per genotype; 2 mice per culture. Tsc1 ****p<0.0001, Mann-Whitney test; Raptor ****p<0.0001, Mann-Whitney test; Rictor ***p=0.0005, Mann-Whitney test; p-S6 Ser240/244 ****p<0.0001, Welch’s t-test; p-4EBP1 T37 p=0.1018, Welch’s t-test; p-Akt Ser473 ***p=0.0003, Welch’s t-test. ns=non-significant. c,d) Correlation of Raptor protein levels to p-S6 Ser240/244 (c) or p-Akt Ser473 (d) levels within each culture, expressed as a percentage of Control. Samples were pooled across hippocampal cultures from *Tsc1^fl/fl^;Rptor^wt/fl or fl/fl^* mice treated with AAV-GFP or AAV-Cre-GFP and harvested on different days (DIV 14-18) to generate a range of Raptor protein levels (includes data from Figs. 2e-f, 3a-b & e-f, and Fig. S2a-g). Solid line depicts non-linear regression. n=48 culture wells. For panel c, r= 0.8907, ****p<0.0001, Spearman correlation. For panel d, r= -0.8662, ****p<0.0001, Spearman correlation. e) Representative western blots from *Tsc1^fl/fl^;Rptor^wt/fl^* hippocampal cultures treated with AAV-GFP (WT;WT) or AAV-Cre-GFP (KO;KO) and harvested on DIV 18. Two independent samples per genotype are shown; this experiment was replicated three times. f) Bar graphs display western blot quantification (mean +/-SEM) for the indicated proteins, expressed as a percentage of Control (WT) levels. n=9 culture wells from 3 independent cultures per genotype; 2 mice per culture. Tsc1 ****p<0.0001; Raptor ****p<0.0001; Rictor ****p<0.0001; p-S6 Ser240/244 **p=0.0052; p-4EBP1 T37 p=0.1621; p-Akt Ser473 *p=0.0235; Welch’s t-tests. For panels b,c,d and f, phospho-proteins were normalized to their respective total proteins. Dots represent data from individual culture wells. Dashed lines at 100% indicate control levels. g) Representative images showing a GFP-expressing control neuron (left panel) and a neighboring AAV-Cre + AAV-Flex-tdTomato transduced Tsc1 KO neuron (middle panel) within the same hippocampal culture. Right panel shows the merged image is on the right. Scale bars=50 μm. h) Quantification (mean +/- SEM) of the average soma area of cultured hippocampal neurons of the indicated genotypes, expressed as a percentage of Control (WT) neurons from the same culture. One-way ANOVA, p<0.0001, F (4, 705) = 19.33; Tsc1-KO vs Tsc1-KO;Rictor-KO, p=0.4030; Tsc1-KO vs Tsc1-KO+Rapa, ****p<0.0001; Tsc1-KO vs Tsc1-KO;Rptor-KO, ****p<0.0001; Tsc1-KO vs. Tsc1-KO;Rptor-Het, ****p<0.0001; Holm-Sidak’s multiple comparisons tests. See also Supplementary Figures 2 and 3.

Our findings suggest that relatively low levels of Raptor protein may be sufficient to sustain mTORC1 signaling in the absence of negative regulation by the Tsc1/2 complex. To further evaluate the relationships between Raptor, mTORC1, and mTORC2, we analyzed results from cultures harvested at different time points, which had complete loss of Tsc1 protein and variable loss of Raptor protein (as in Supplementary Fig. 2b,c). We plotted the correlation of Raptor protein levels with p-S6 and p-Akt, expressed as a percentage of control cultures from the same genotype and time point. We observed a non-linear relationship whereby p-S6 remained elevated in Tsc1-cKO cultures even when Raptor protein levels were reduced by up to 80% (Fig. 3c). Only when Raptor levels dropped below 20% of control did we observe a reduction in p-S6 levels. The inverse relationship was observed for p-Akt, which was consistently reduced with Tsc1 loss and increased sharply as Raptor levels fell below 20% (Fig. 3d). Similar relationships were observed in hippocampal cultures with downregulation of Raptor alone, which were wild-type for *Tsc1* (Supplementary Fig. 2h-j). In this case, mild mTORC1 suppression was observed with 20-80% loss of Raptor protein; however, when Raptor levels were less than 15% of control, p-S6 was strongly suppressed and p-Akt was robustly increased. Together, these analyses reveal a non-linear relationship between Raptor and mTORC1 signaling such that relatively low Raptor protein levels can maintain mTORC1 activity in neuronal cultures.

These results suggest that partial reduction of Raptor protein may be an effective strategy to normalize both mTORC1 and mTORC2 signaling in Tsc1-cKO neurons. This would avoid the strong suppression of both complexes observed with rapamycin treatment and the inhibition of mTORC1 and increase in mTORC2 activity observed with complete Raptor loss. To test this, we assessed mTOR signaling in cultures from *Tsc1^fl/fl^;Rptor^wt/fl^* mice, which had partial loss of Raptor in the Tsc1-cKO background. We harvested Tsc1-cKO;Raptor-cHet cultures on DIV 18 and found that the elevation in p-S6 was reduced compared to *Tsc1* deletion alone (Fig. 3e,f) (Tsc1-cKO;Raptor-cHet 125.9% +/-6.6 % of control vs. Tsc1-cKO 181.6% +/-15.5% of control). In addition, while p-Akt-473 was slightly reduced compared to control, the reduction was less than in Tsc1-cKO cultures (Fig. 3e,f) (Tsc1-cKO;Raptor-cHet 74.5% +/- 9.1% of control vs.

Tsc1-cKO 58.2% +/- 4.0% of control). These data show that loss of one copy of *Rptor* can significantly ameliorate, although not completely prevent, the mTORC1 and mTORC2 signaling abnormalities associated with Tsc1 loss. Supplementary Figure 2k provides a summary of the effects of rapamycin, Rictor, and Raptor manipulations on p-S6 and p-Akt levels in Tsc1-cKO cultures.

### Reduction of Raptor prevents hypertrophy of Tsc1-cKO neurons

Cellular hypertrophy is a well-established consequence of mTORC1 hyperactivity^31^. To test whether genetic reduction of Raptor could improve somatic hypertrophy, we sparsely labeled cultured neurons with florescent proteins and measured soma size. In this experiment, control cells expressed GFP and mutant cells within the same culture expressed nuclear Cre-mCherry and cytoplasmic tdTomato (Fig. 3g). We observed the expected increase in soma size of Tsc1- cKO neurons compared to controls, which was reversed by four-day treatment with rapamycin (Fig. 3h and Supplementary Fig. 3a-f). Homozygous deletion of *Rictor* did not prevent cellular hypertrophy as Tsc1-cKO;Rictor-cKO neurons exhibited significantly increased soma size compared to controls (Fig. 3h and Supplementary Fig. 3g-i). By contrast, both homozygous and heterozygous deletion of *Rptor* reduced the size of Tsc1-cKO neurons, effectively normalizing soma area to control levels (Fig. 3h and Supplementary Fig. 3j-o). Together, these results demonstrate that genetic reduction of Raptor, but not Rictor, is sufficient to prevent hypertrophy of Tsc1-cKO neurons *in vitro*.

### Genetic reduction of Raptor, but not Rictor, extends the life span of Tsc1-cKO mice

To test whether Raptor reduction could improve TSC-related phenotypes *in vivo*, we used the *Emx1*^IRESCre^ (Emx1-Cre)^49^ mouse line to induce *Tsc1* deletion around embryonic day 9.5 (E9.5), primarily in excitatory neurons of the cortex and hippocampus (Fig. 4a). Deletion of *Tsc1* in Emx1-expressing cells leads to a severe phenotype characterized by poor development, seizures, and premature mortality in the first few weeks of life^50, 51^. To test whether genetic reduction of mTORC1 or mTORC2 could prevent these phenotypes, we generated mice with heterozygous or homozygous deletion of *Rptor* or *Rictor* in the Tsc1-cKO;Emx1-Cre background (Fig. 4b). We monitored the survival and body weight of mice born from these crosses for the first 40 postnatal days. As expected, *Tsc1^fl/fl^;Emx1-Cre^+^* (Tsc1-cKO) mice exhibited premature mortality with a median survival of 18.5 days (Fig. 4c,d and Supplementary Table 1). Tsc1-cKO mice of both sexes also had reduced body weight, although the timing of death was not correlated to weight (Fig. 4e-h, Supplementary Fig. 4a, and Supplementary Table 2).

**Figure 4.**
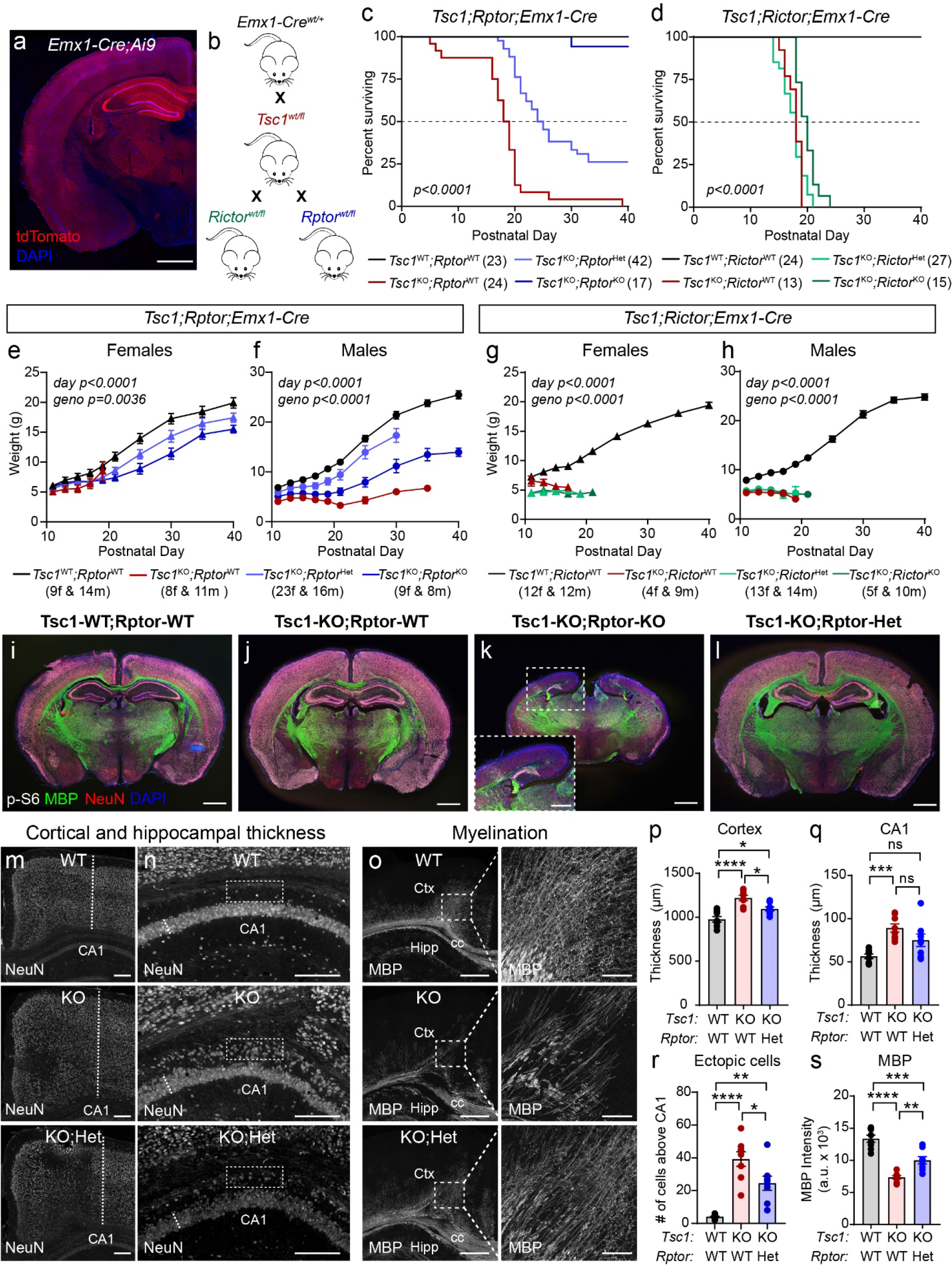
Reduction of *Rptor* prolongs the life span and improves forebrain development of Tsc1-cKO mice. a) Representative image showing one hemisphere of a coronal section from an *Emx1-Cre^+^;Ai9^+^* mouse. The Cre-dependent tdTomato reporter (red) is expressed broadly in the cortex and hippocampus. DAPI labels cell nuclei (blue). Scale bar=1 mm b) Schematic of the crosses used to generate experimental mice. *Emx1-Cre^+^* mice were crossed with *Tsc1^wtl/fl^* mice. This line was subsequently bred to either *Rictor^wt/fl^* or *Rptor^wt/fl^* mice. c,d) Survival analysis of *Tsc1;Rptor;Emx1-Cre* (c) and *Tsc1;Rictor;Emx1-Cre* (d) mice of the indicated genotypes. The number of mice for each genotype is indicated in parentheses. Dashed lines indicate 50% of the population surviving. P values from Log-rank Mantel-Cox tests are shown on the graphs. e-h) Mean +/- SEM body weight in grams measured from postnatal day 11 to 40 for mice of the indicated sex and genotype. The number of mice for each genotype and is indicated in parentheses. f=females, m=males. Mixed-effects model (REML) statistics: *Tsc1;Rptor;Emx1- Cre* females (e), day p<0.0001, F(1.447, 51.30) = 398.9; geno p=0.0036, F(3, 45) = 5.192. *Tsc1;Rptor;Emx1-Cre* males (f), day p<0.0001, F(2.287, 79.03) = 197.4; geno p<0.0001, F (3, 45) = 26.85. *Tsc1;Rictor;Emx1-Cre* females (g), day p<0.0001, F (4.257, 69.52) = 221.1; geno p<0.0001, F (3, 30) = 53.74. *Tsc1;Rictor;Emx1-Cre* males (h), day p<0.0001, F (2.389, 48.04) = 370.4; geno ****p<0.0001, F (3, 41) = 28.86. i-l) Representative whole brain coronal sections from mice of the indicated genotypes. Immunostaining for p-S6 Ser240/244 is in grey, myelin basic protein (MBP) is in green, NeuN is in red and DAPI-labeled nuclei are in blue. In panel k, inset shows zoomed-in image of the cortex and hippocampal region. Scale bars=1 mm. Panel k inset, scale bar=500 μm m) Representative images of the cortex with NeuN immunostaining. Dashed lines denote the measurement of cortical thickness. WT=Tsc1-WT;Rptor-WT, KO=Tsc1-KO;Rptor-WT, KO;Het=Tsc1-KO;Rptor-Het. Scale bars=500 μm. n) Representative images of the CA1 region of the hippocampus with NeuN immunostaining from mice of the indicated genotypes. Dashed lines denote the measurement of CA1 thickness. Boxed regions highlight the area above the CA1 that contains ectopic p-S6 expressing neurons in Tsc1-KO;Rptor-WT mice, which are reduced in Tsc1-KO;Rptor-Het mice (quantified in panel r). Scale bars=250 μm o) Representative images of MBP immunostaining in the cortex (Ctx) & dorsal hippocampus (Hipp) from mice of the indicated genotypes. cc=corpus callosum. Scale bars=500 μm. Right panels show higher magnification of the boxed regions in the cortex. Scale bars=100 μm. p) Mean +/- SEM cortical thickness for the indicated genotypes. For panels p-s, dots represent values from individual mice, n=8 mice per genotype. One-way ANOVA, p<0.0001, F (2, 21) = 18.40; WT vs KO, ****p<0.0001; WT vs KO;Het, *p=0.0226; KO vs KO;Het, *p=0.0157, Sidak’s multiple comparisons tests. q) Mean +/- SEM CA1 thickness for the indicated genotypes. One-way ANOVA, p=0.0010, F (2, 21) = 9.703; WT vs KO, ***p=0.0008; WT vs KO;Het, p=0.0628; KO vs KO;Het, p=0.1968 Sidak’s multiple comparisons tests. ns=non-significant. r) Mean +/- SEM number of p-S6 positive neurons above the CA1 (boxed regions in panel n) for the indicated genotypes. One-way ANOVA, p<0.0001, F (2, 21) = 24.77; WT vs KO, ****p<0.0001; WT vs KO;Het, **p=0.0016; KO vs KO;Het, *p=0.0237 Sidak’s multiple comparisons tests. s) Mean +/- SEM bulk MBP fluorescence intensity in the boxed cortical region shown in panel o for the indicated genotypes. One-way ANOVA, p<0.0001, F (2, 21) = 39.19; WT vs KO, ****p<0.0001; WT vs KO;Het, ***p=0.0002; KO vs KO;Het, **p=0.0025; Sidak’s multiple comparisons tests. See also Supplementary Figure 4 and Supplementary Tables 1 and 2.

Loss of Raptor extended the life span of Tsc1-cKO mice in a gene dose-dependent manner, with homozygous *Rptor* deletion resulting in near normal survival (16/17 *Tsc1^fl/fl^;Rptor^fl/fl^;Emx1- Cre^+^* animals survived until at least P40, Fig. 4c and Supplementary Table 1). Heterozygous deletion of *Rptor* led to a significant shift in the median survival of Tsc1-cKO mice from 18.5 to 24.5 postnatal days and 26% of the Tsc1-cKO;Raptor-cHet mice survived past P40 (Fig. 4c). While Tsc1-cKO;Raptor-cHet males and females had identical median survival, the length of life span extension was sex-dependent. After P26, the surviving female population stabilized with 40% of Tsc1-cKO;Raptor-cHet females surviving past P40 (Supplementary Fig. 4b and Supplementary Table 1). By contrast, the Tsc1-cKO;Raptor-cHet male population steadily declined and only 5% of mice survived past P40 (Supplementary Fig. 4b and Supplementary Table 1). Notably, while Tsc1-cKO;Raptor-cKO mice regularly survived past P40, they exhibited significantly decreased body weight (Fig. 4e,f and Supplementary Table 2). Similar body weight changes were observed with homozygous loss of *Rptor* in mice WT for *Tsc1* (Supplementary Table 2). Compared to WT mice, Tsc1-cKO;Raptor-cHet mice had reduced body weight but were significantly larger than Tsc1-cKO;Raptor-WT mice, suggesting improved physical development (Fig. 4e,f and Supplementary Table 2). The body weight differences between Tsc1;Raptor mice of different genotypes generally persisted among surviving mice at P150 (Supplementary Fig. 4c).

A recent report in *Pten^fl/fl^;CamKII*-*Cre^+^* mice (Pten-cKO) showed that concurrent deletion of *Pten* and *Rictor* significantly shifted median survival from P∼50 to ∼110 days; however, no mice survived past P130. Surprisingly, homozygous *Rptor* deletion did not significantly affect the median survival age of Pten-cKO mice^38^. We generated *Tsc1;Rictor;Emx1-Cre* mice and found that heterozygous loss of *Rictor* did not affect the survival of Tsc1-cKO mice and homozygous loss of *Rictor* only slightly shifted the median survival from P18 to P20, with no Tsc1-cKO;Rictor-cKO animals surviving past P24 (Fig. 4d and Supplementary Table 1). It was previously shown that *Rictor^fl/fl^;Emx1-Cre^+^* animals exhibited reduced body weight^52^. Consistent with this, we observed decreased body weight in mice with homozygous loss of *Rictor*, independent of *Tsc1* genotype (Fig. 4g,h and Supplementary Table 2). Together, these findings indicate that premature mortality due to loss of Tsc1 in forebrain excitatory neurons can be prevented by concomitant downregulation of mTORC1, but not mTORC2.

### Genetic downregulation of Raptor improves multiple TSC-related brain phenotypes

Given that downregulation of Raptor extended the life span of Tsc1-cKO mice in a gene-dose dependent manner, we next examined whether TSC-related brain phenotypes could be improved. We explored a battery of common phenotypes in mouse models of TSC including macrocephaly, hypertrophic neurons, impaired myelination, and reactive astrogliosis^50, 51, 53^. We harvested brain tissue from P14-15 mice and found the expected cortical hypertrophy and elevated p-S6 phosphorylation in the forebrain of Tsc1-cKO mice (Fig. 4i,j). Strikingly, when we examined the brains of Tsc1-cKO;Raptor-cKO mice, we found decreased overall brain size, severely reduced cortical thickness, and lack of a clear hippocampal structure (Fig. 4k and Supplementary Fig. 4d,e), despite near normal survival of these mice (see Fig. 4c). Underdeveloped cortical and hippocampal structures were also observed in Tsc1-WT;Raptor-cKO mice (Supplementary Fig. 4f), indicating that impaired forebrain development was due to *Rptor* deletion alone, independent of *Tsc1* genotype. In contrast, heterozygous deletion of *Rptor* did not cause changes in overall brain architecture (Supplementary Fig. 4f). These results demonstrate that while loss of Tsc1 causes cortical hypertrophy, embryonic suppression of mTORC1 signaling leads to severely impaired cortical and hippocampal development.

Given the significant neurodevelopmental abnormalities associated with complete loss of Raptor, we tested whether partial Raptor reduction could improve developmental brain phenotypes in Tsc1-cKO mice. We found that loss of one copy of *Rptor* improved overall brain architecture, reduced cortical hypertrophy, ameliorated hippocampal CA1 and dentate gyrus (DG) lamination defects, and improved cortical myelination, resulting in an intermediate phenotype between WT and Tsc1-cKO mice (Fig. 4l-s and Supplementary Fig. 5a-c). At the cellular level, heterozygous loss of *Rptor* reduced mTORC1 signaling, as measured by p-S6 levels, and partially prevented neuronal hypertrophy in the cortex, CA1 and DG regions (Fig. 5a-n and Supplementary Fig. 5d-g).

**Figure 5.**
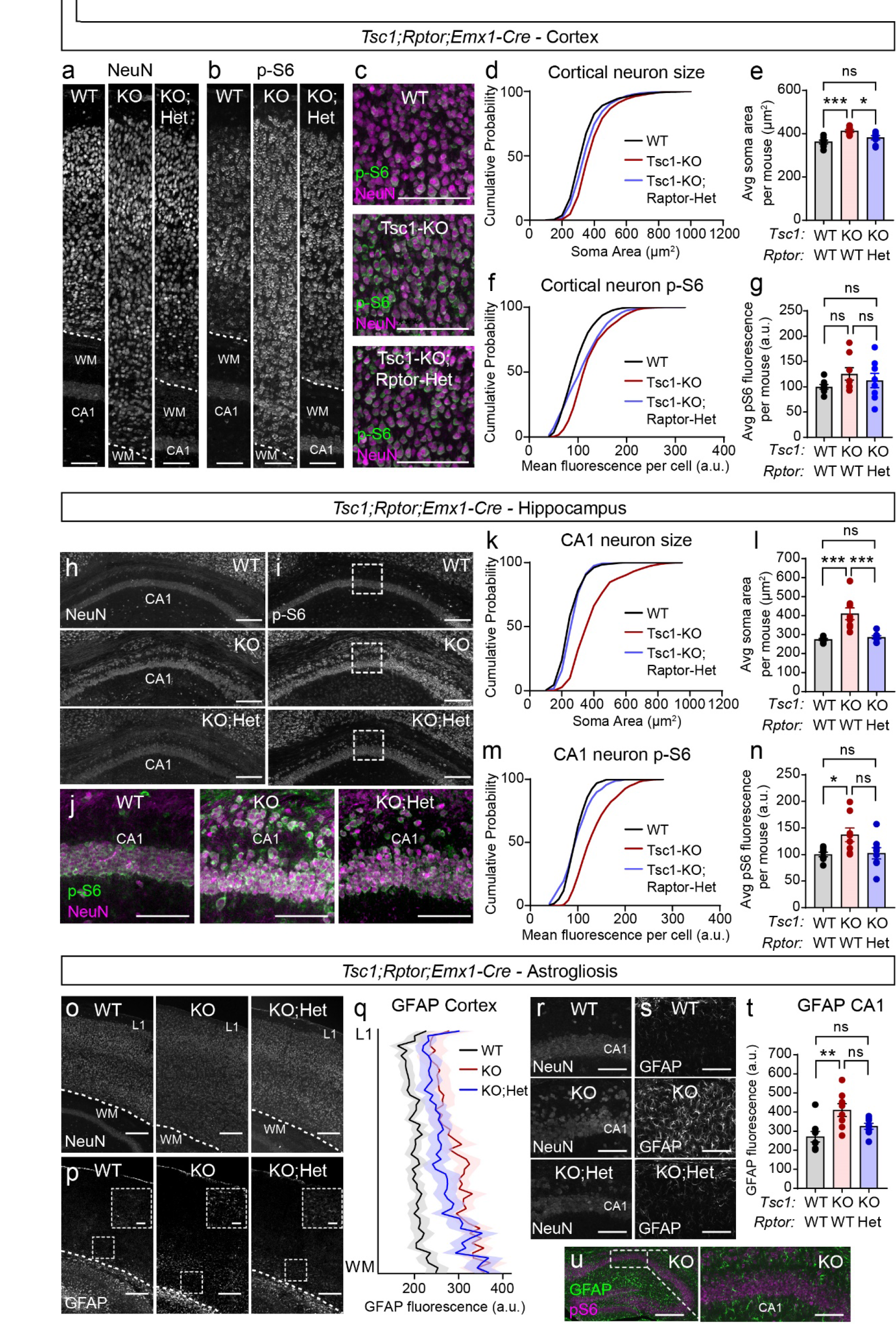
Heterozygous deletion of *Rptor* improves cellular phenotypes in Tsc1-cKO mice. a,b) Representative images of somatosensory cortex showing NeuN (left panels, a) and p-S6 Ser240/244 (right panels, b) immunostaining in Tsc1-WT;Rptor-WT (WT), Tsc1-KO;Rptor-WT (KO), and Tsc1-KO;Rptor-Het (KO;Het) mice. Dashed lines indicate the border of cortical layer 6 and the corpus callosum. Scale bars=100 μm. WM=white matter c) Representative zoomed-in images of the cortex (midway between the cortical plate and white matter) showing p-S6 Ser240/244 (green) and NeuN (magenta) immunostaining for the indicated genotypes. Scale bars=100 μm. d) Cumulative distributions of cortical neuron soma area for the indicated genotypes. n=1601 WT, 1605 Tsc1-KO, and 1602 Tsc1-KO;Rptor-Het neurons from 8 mice per genotype. Kruskal-Wallis test, p<0.0001; WT vs KO, p<0.0001; WT vs KO;Het, p<0.0001; KO vs KO;Het, p<0.0001; Dunn’s multiple comparisons tests. e) Mean +/- SEM cortical neuron soma area per mouse for the indicated genotypes. Dots represent values from individual mice. n=8 mice per genotype. One-way ANOVA, p=0.0011, F (2, 21) = 9.629; WT vs KO, ***p=0.0008, WT vs KO;Het, p=0.2760; KO vs KO;Het, *p=0.0447 Sidak’s multiple comparisons tests. ns=non-significant. f) Cumulative distributions of cortical p-S6 levels per neuron for the indicated genotypes. n is the same as for panel d. Kruskal-Wallis test, p<0.0001; WT vs KO, p<0.0001; WT vs KO;Het, p<0.0001; KO vs KO;Het, p<0.0001; Dunn’s multiple comparisons tests. g) Mean +/- SEM cortical neuron p-S6 levels per mouse for the indicated genotypes. Dots represent values from individual mice. n=8 mice per genotype. One-way ANOVA, p=0.2881, F (2, 21) = 1.321; WT vs KO, p=0.3162; WT vs KO;Het, p= 0.8216, KO vs KO;Het, p=0.7989; Sidak’s multiple comparisons tests. h,i) Representative images of hippocampal area CA1 from mice of the indicated genotypes. Left panels show NeuN (h) and right panels show p-S6 Ser240/244 (i) immunostaining. Scale bars=250 μm. j) Representative zoomed-in images of the boxed region in panel i, showing p-S6 Ser240/244 (green) and NeuN (magenta) immunostaining for the indicated genotypes. Scale bars=100 μm. k) Cumulative distributions of CA1 neuron soma area for the indicated genotypes. n=568 WT, 561 Tsc1-KO, and 561 Tsc1-KO;Rptor-Het neurons from 8 mice per genotype. Kruskal-Wallis test, p<0.0001; WT vs KO, ****p<0.0001; WT vs KO;Het, *p=0.0133; KO vs KO;Het, ****p<0.0001; Dunn’s multiple comparisons tests. l) Mean +/- SEM CA1 neuron soma area per mouse for the indicated genotypes. Dots represent values from individual mice. n=8 mice per genotype. One-way ANOVA, p<0.0001, F(2, 21) = 15.89; WT vs KO, ***p=0.0001; WT vs KO;Het, p=0.9655; KO vs KO;Het, ***p=0.0004; Sidak’s multiple comparisons tests. m) Cumulative distributions of CA1 p-S6 levels per neuron for the indicated genotypes. n is the same as for panel k. Kruskal-Wallis test, p<0.0001; WT vs KO, p<0.0001; WT vs KO;Het, p=0.7622; KO vs KO;Het, p<0.0001; Dunn’s multiple comparisons tests. n) Mean +/- SEM CA1 neuron p-S6 levels per mouse for the indicated genotypes. Dots represent values from individual mice. n=8 mice per genotype. One-way ANOVA, p=0.0263, F (2, 21) = 4.346; WT vs KO, *p=0.0464; WT vs KO;Het, p=0.9981; KO vs KO;Het, p=0.0650; Sidak’s multiple comparisons tests. o,p) Representative images of the somatosensory cortex from mice of the indicated genotypes. Top panels show NeuN (o) and bottom panels show GFAP (p) immunostaining. Scale bars=250 μm. Dashed lines indicate the border of cortical layer 6 and the beginning of white matter of the corpus callosum. L1=cortical layer 1. Zoomed-in inset scale bars=100μm. q) GFAP immunofluorescence across cortical layers for mice of the indicated genotypes. Lines represent the mean, shaded regions represent the SEM. n=8 mice per genotype. r,s) Representative images of the hippocampal CA1 region from mice of the indicated genotypes. Left panels show NeuN (r) and right panels show GFAP (s) immunostaining. Scale bars=100 μm. t) Mean +/- SEM bulk GFAP fluorescence in the CA1 region per mouse for the indicated genotypes. Dots represent values from individual mice. n=8 mice per genotype. Kruskal-Wallis test, p=0.0087; WT vs KO, **p=0.0063; WT vs KO;Het, p=0.2691; KO vs KO;Het, p=0.5038; Dunn’s multiple comparisons tests. u) Representative images showing GFAP (green) and p-S6 Ser240/244 (magenta) immunostaining in the hippocampus of a Tsc1-KO mouse Scale bar=500 μm. Right panel shows a zoomed-in image of the boxed region. Scale bar=100μm. See also Supplementary Figure 5 and Supplementary Table 3.

Glial abnormalities have previously been observed in the *Tsc1^fl/fll^;Emx1-Cre^+^* mouse model^50, 51^. Astrocytes specifically appear dysmorphic, exhibiting hypertrophic processes and increased expression of glial fibrillary acidic protein (GFAP)^50, 51^. Consistent with these findings, we observed a significant increase in GFAP fluorescence intensity across all cortical layers of Tsc1-cKO mice, that was partially attenuated in Tsc1-cKO;Raptor-cHet mice (Fig. 5o-q). We also observed a significant increase in GFAP fluorescence intensity in the hippocampal CA1 region of Tsc1-cKO mice that was restored to near WT levels in Tsc1-cKO;Rptor-cHet mice (Fig. 5r-t). Similar to prior studies, the GFAP-positive cells within the hippocampus of Tsc1-cKO mice did not exhibit high p-S6 levels^50, 51^ (Fig. 5u). Notably, the severity of phenotypes caused by Tsc1 loss and their ability to be prevented by Raptor reduction were both cell type- and sex-dependent (Supplementary Fig. 5h-k and Supplementary Table 3). Taken together, these results demonstrate that partial reduction of Raptor in the forebrain is sufficient to improve multiple developmental brain phenotypes resulting from loss of Tsc1.

### Hyperexcitability of Tsc1-cKO neurons is reduced by Raptor downregulation

*Tsc1^fl/fl^;Emx1-Cre^+^* mice exhibited behavioral seizures that were observable after the first two weeks of life. Given the challenges of performing *in vivo* EEG analysis in very young animals, and reports that cultured neurons with TSC-associated mutations show network hyperactivity^54–,56^, we used calcium imaging of hippocampal cultures from *Tsc1^fl/fl^;Rptor^wtl/fl^;Emx1-Cre^+^* mice to test whether Raptor reduction could prevent epileptiform activity. We delivered an AAV encoding the fluorescent calcium indicator jRGECO1a^57^ on DIV 2 and imaged calcium dynamics on DIV 14 (Fig. 6a-j and Supplementary Fig. 6a,b). Analysis of individual Ca^2+^ transients (Supplementary Fig. 6c,d) showed that cultures from Tsc1-cKO mice exhibited more frequent events, which were, on average, larger in amplitude and longer in duration than in control neurons (Fig. 6k-n). We calculated the area under the curve (AUC) for each Ca^2+^ transient and found that the average AUC was significantly larger in Tsc1-cKO neurons (Fig. 6o). In cultures from Tsc1-cKO;Raptor-cHet mice, Raptor downregulation significantly reduced the average frequency, amplitude, and AUC but had no effect on the duration of Ca^2+^ transients compared to Tsc1 loss alone (Fig. 6k-o). These results show that neuronal hyperactivity caused by Tsc1 loss can be ameliorated by Raptor reduction.

**Figure 6.**
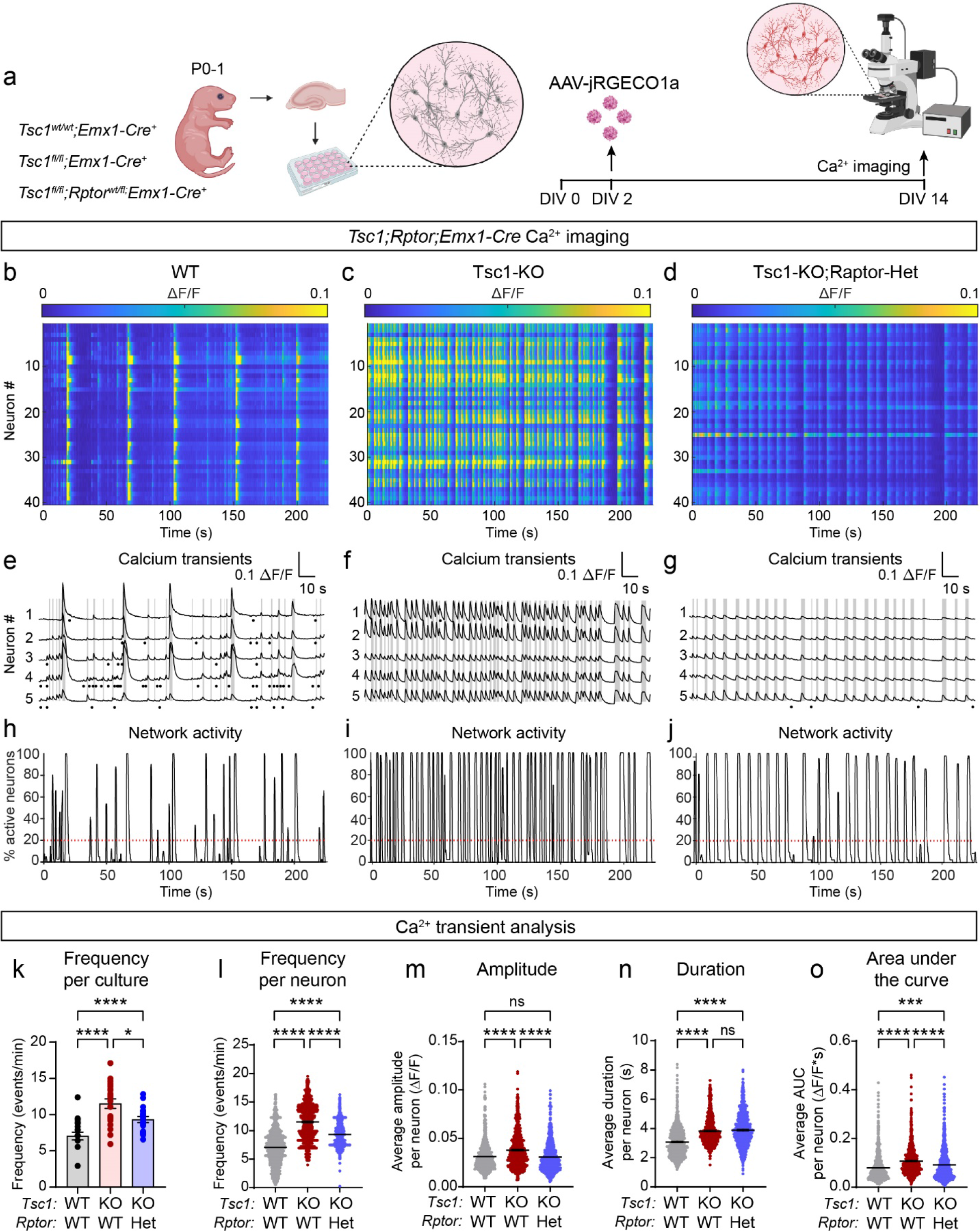
Raptor downregulation reduces hyperactivity of Tsc1-cKO neurons. a) Schematic of the experiment. Primary hippocampal cultures were prepared from P0-1 mice of different genotypes. An AAV expressing the calcium indicator jRGeco1a was added on DIV 2 and cells were imaged on DIV 14. Created with BioRender.com b-d) Representative heatmaps of ΔF/F for 40 neurons imaged in a field of view from *Tsc1^wt/wt^;Emx1-Cre^+^* (WT, b), *Tsc1^fl/fl^;Emx1-Cre^+^* (Tsc1-KO, c), and *Tsc1^fl/fl^;Rptor^wt/fl^;Emx1-Cre^+^* (Tsc1-KO;Raptor-Het, d) cultures. e-g) Ca^2+^ transients from 5 representative neurons imaged in a field of view in WT (e), Tsc1-KO (f) and Tsc1-KO;Raptor-Het (g) cultures. Grey lines indicate network events with more than 20% of neurons in the field of view active at the same time. Black dots represent spontaneous Ca^2+^ transients that were not part of network events. h-j) Graphs display the percentage of neurons in the field of view that were active at a given time for a representative WT (h), Tsc1-KO (i) and Tsc1-KO;Raptor-Het (j) culture. Red dashed lines at 20% indicate the threshold for what was considered a network event. k) Mean +/- SEM calcium transient frequency per culture. Dots represent values from individual cultures. n=20 culture wells from 6-7 independent culture preps per genotype, 1 pup per prep. One-way ANOVA, p<0.0001, F (2, 57) = 16.88; WT vs KO, ****p<0.0001; WT vs KO;Het, *p=0.0130; KO vs KO;Het, *p=0.0187; Sidak’s multiple comparisons tests. l) Scatter dot plot of the Ca^2+^ transient frequency per neuron for the indicated genotypes. Black lines indicate mean +/- SEM. n=800 neurons per genotype. Kruskal-Wallis test, p<0.0001; WT vs KO, p<0.0001; WT vs KO;Het, p<0.0001. KO vs. KO;Het, p<0.0001; Dunn’s multiple comparisons tests. m) Scatter dot plot of the average Ca^2+^ transient amplitude per neuron for the indicated genotypes. n is the same as for panel l. Kruskal-Wallis test, p<0.0001; WT vs KO, ****p<0.0001; WT vs KO;Het, p=0.9211; KO vs. KO;Het, ****p<0.0001; Dunn’s multiple comparisons tests. ns=non-significant. n) Scatter dot plot of the average Ca^2+^ transient duration per neuron for the indicated genotypes. n is the same as for panel l. Kruskal-Wallis test, p<0.0001; WT vs KO, ****p<0.0001; WT vs KO;Het, ****p<0.0001; KO vs. KO;Het, p>0.9999; Dunn’s multiple comparisons tests. o) Scatter dot plot of the average Ca^2+^ transient area under the curve (AUC) per neuron for the indicated genotypes. n is the same as for panel l. Kruskal-Wallis test, p<0.0001; WT vs KO, ****p<0.0001; WT vs KO;Het, ***p=0.0001; KO vs. KO;Het, ****p<0.0001; Dunn’s multiple comparisons tests. See also Supplementary Figures 6-8.

To investigate how loss of Tsc1 affects network activity patterns, we analyzed ‘network events’, which were defined by synchronous activity of more than 20% of neurons in the field of view (see Fig. 6h-j). We found that a significantly larger proportion of neurons participated in each network event in Tsc1-cKO cultures compared to controls and that these events occurred with greater frequency (Supplementary Fig. 6e,f). We measured the percentage of time each culture exhibited coordinated network activity during the recording period and found that this was significantly increased in Tsc1-cKO cultures compared to controls (Supplementary Fig. 6g). We analyzed the individual Ca^2+^ transients that occurred during network events and found that these were larger and lasted longer in Tsc1-cKO neurons compared to controls (Supplementary Fig. 6h-j). While partial Raptor downregulation did not significantly reduce the frequency or duration of network activity, it was sufficient to reduce the proportion of neurons participating in network events (Supplementary Fig. 6e-g). Heterozygous deletion of *Rptor* also reduced the amplitude and AUC, but not the duration, of individual Ca^2+^ transients occurring within network events (Supplementary Fig. 6h-j). Collectively, these data show that heterozygous loss of *Rptor* reduces but does not completely prevent network hyperactivity caused by loss of Tsc1.

### Heterozygous loss of *Tsc1* does not induce network hyperexcitability

Whether partial loss of Tsc1/2 complex function can drive neuronal hyperactivity remains inconclusive as reports using cultured human neurons derived from TSC patient cells, which are heterozygous for the *TSC1* or *TSC2* mutation, present conflicting findings^56, 58, 59^. To investigate this in our model, we generated *Tsc1^wt/fl^;Emx1-Cre^+^* (Tsc1-cHet) mice and found that they had normal body weight and survival (Supplementary Fig. 7a-c), with no spontaneous seizures observed. We prepared hippocampal cultures from these mice and found that in contrast to homozygous *Tsc1* deletion, heterozygous loss of *Tsc1* did not induce somatic hypertrophy, increase p-S6, or reduce p-Akt levels (Supplementary Fig. 7d-j).

We transduced Tsc1-cHet cultures with AAV-jRGECO1a and examined whether loss of one copy of *Tsc1* altered Ca^2+^ activity dynamics (Supplementary Fig. 8a-f). Tsc1-cHet neurons showed no difference in the frequency of Ca^2+^ transients per culture and had a small decrease in the frequency of events per neuron (Supplementary Fig. 8g,h). While the average event duration and AUC per neuron were not different between Tsc1-cHet and control neurons, the average amplitude of these events was larger in Tsc1-cHet neurons. (Supplementary Fig. 8i-k). Analysis of network events showed that a smaller proportion of Tsc1-cHet neurons participated in these events compared to control (Supplementary Fig. 8l). The frequency and total duration of network events were similar between control and Tsc1-cHet cultures, as was the amplitude of individual transients (Supplementary Fig. 8m-o). The duration and AUC of Ca^2+^ transients occurring within network events were slightly decreased in Tsc1-cHet neurons (Supplementary Fig. 8p,q). Together these data indicate that while loss of one copy of *Tsc1* causes some alterations in Ca^2+^ dynamics in mouse hippocampal neurons, it is not sufficient to induce neuronal or network hyperactivity.

### Postnatal Raptor reduction using AAV-shRptor

The results described above showed that ∼50% reduction of Raptor protein concurrent with *Tsc1* deletion reduced mTORC1 hyperactivity and improved multiple developmental and functional phenotypes resulting from Tsc1 loss. To determine whether Raptor reduction could be used therapeutically, we used a viral approach to reduce Raptor expression postnatally in the forebrain of Tsc1-cKO mice. Our biochemical experiments indicated that ∼80% loss of Raptor may be most effective at normalizing mTORC1 and mTORC2 signaling in neurons with *Tsc1* deletion (see Fig. 3). Therefore, we generated AAVs encoding an shRNA targeting Raptor (shRptor)^60^ or a scrambled control (shControl) under a ubiquitously expressed promoter. We transduced hippocampal cultures from *Tsc1^wt/wt or fl/fl^;Emx1-Cre^+^* mice with these viruses and found that shRptor reduced Raptor protein levels by ∼90% in WT neurons and 65-70% in Tsc1-cKO neurons (Supplementary Fig. 9a-h). The lesser reduction of Raptor in Tsc1-cKO cultures compared to WT was likely due to upregulation of Raptor protein in these cells (Supplementary Fig. 9d). In Tsc1-cKO cultures, shRptor reduced p-S6 levels, relieved p-Akt suppression, and completely rescued neuronal hypertrophy (Supplementary Fig. 9f,h-j). In this experiment, in which *Tsc1* was deleted embryonically, we observed a significant increase in 4E-BP1 phosphorylation in Tsc1-cKO neurons treated with shControl (Supplementary Fig. 9g). However, similar to rapamycin treatment^60^, 4E-BP1 phosphorylation was resistant to shRptor treatment (Supplementary Fig. 9g).

To examine the efficacy of postnatal Raptor downregulation as a therapeutic strategy, we injected AAV9-shRptor-EYFP or shControl into the cortex and hippocampus of P0 *Tsc1^fl/fl^;Emx1-Cre^+^* (Tsc1-cKO) and *Tsc1^wt/wt^;Emx1-Cre^+^* (WT) mice (Fig. 7a). We examined brain sections from P16 mice and found that shRptor consistently reduced the p-S6 levels and soma size of Tsc1-cKO neurons within the cortex and hippocampus (Fig. 7b-m and Supplementary Fig. 10). We noted that the degree of reduction compared to shControl-treated WT neurons varied by brain region, ranging from an intermediate phenotype in the DG, normalization to WT levels in CA1, and reduction below WT levels in the somatosensory cortex (Fig. 7b-m and Supplementary Fig. 10). In WT mice, shRptor strongly downregulated neuronal p-S6 levels in the cortex and CA1, but not in the DG, and modestly reduced soma size within all regions (Fig. 7b-m and Supplementary Fig. 10). Thus, postnatal downregulation of Raptor improved cellular phenotypes in mice with prenatal *Tsc1* deletion.

**Figure 7.**
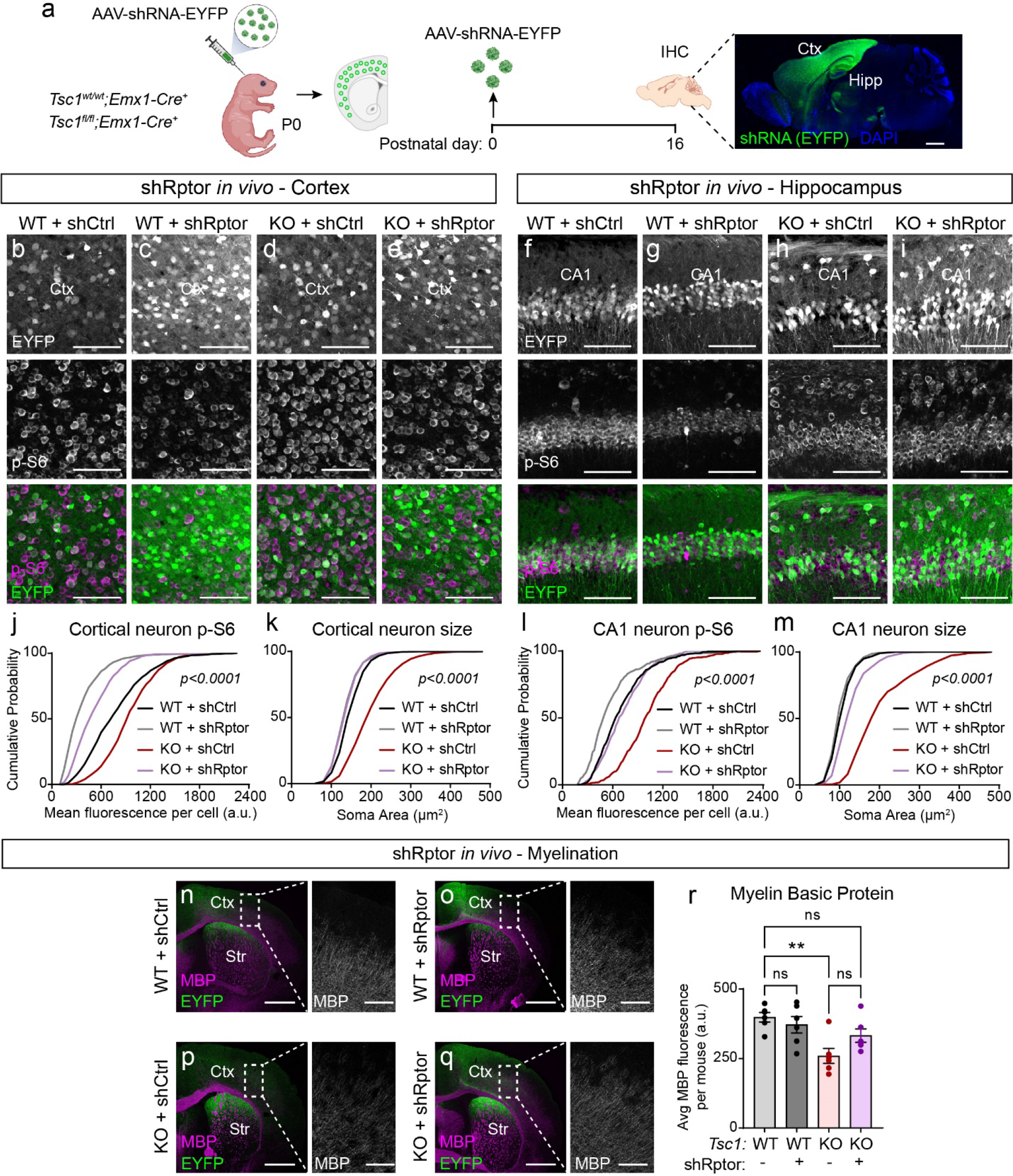
Postnatal Raptor reduction improves cellular phenotypes in Tsc1-cKO mice. a) Schematic of the experiment. P0 pups received intracranial injections with AAV-shRptor-EYFP or AAV-shControl-EYFP (shCtrl). Brains were analyzed by immunohistochemistry at P16. Example sagittal brain image of an injected mouse at P16 showing EYFP expression in green and DAPI in blue. Ctx=cortex; Hipp=hippocampus. Scale bar=1 mm. Created with BioRender.com. b-e) Representative images of the primary somatosensory cortex of *Tsc1^wt/wt^;Emx1-Cre^+^* (WT, b-c) and *Tsc1^fl/fl^;Emx1-Cre^+^* (KO, d-e) mice injected with shRptor-EYFP or shCtrl-EYFP virus showing EYFP (top panels) and p-S6 240/244 immunostaining (middle panels). Bottom panels show merged images. Scale bars=100 μm. f-i) Representative images of the CA1 region of WT (f-g) and KO (h-i) mice injected with shRptor-EYFP or shCtrl-EYFP virus showing EYFP (top panels) and p-S6 240/244 immunostaining (middle panels). Bottom panels show merged images. Scale bars=100 μm. j) Cumulative distributions of p-S6 levels in cortical EYFP+ neurons for the indicated genotypes. n=1086 WT+shCtrl, 1095 WT+shRptor, 1095 Tsc1-KO+shCtrl, and 1087 Tsc1-KO+shRptor neurons from 6 mice per group. Kruskal-Wallis test, p<0.0001; WT+shCtrl vs WT+shRptor, p<0.0001; WT+shCtrl vs Tsc1-KO+shCtrl, p<0.0001; WT+shCtrl vs Tsc1-KO+shRptor, p<0.0001; Tsc1-KO+shCtrl vs Tsc1-KO+shRptor, p<0.0001; Dunn’s multiple comparison tests. k) Cumulative distributions of EYFP+ cortical neuron soma area for the indicated genotypes. n is the same as in panel j. Kruskal-Wallis test, p<0.0001; WT+shCtrl vs WT+shRptor, p<0.0001; WT+shCtrl vs Tsc1-KO+shCtrl, p<0.0001; WT+shCtrl vs Tsc1-KO+shRptor, p<0.0001; Tsc1-KO+shCtrl vs Tsc1-KO+shRptor, p<0.0001; Dunn’s multiple comparison tests. l) Cumulative distributions of p-S6 levels in CA1 EYFP+ neurons for the indicated genotypes. n=514 WT+shCtrl, 507 WT+shRptor, 512 Tsc1-KO+shCtrl, and 497 Tsc1-KO +shRptor neurons from 6 mice per group. Kruskal-Wallis test, p<0.0001; WT+shCtrl vs WT+shRptor, p<0.0001; WT+shCtrl vs Tsc1-KO+shCtrl, p<0.0001; WT+shCtrl vs Tsc1-KO+shRptor, p=0.4315; Tsc1-KO+shCtrl vs Tsc1-KO+shRptor, p<0.0001; Dunn’s multiple comparison tests. m) Cumulative distributions of EYFP+ CA1 neuron soma area for the indicated genotypes. n is the same as for panel l. Kruskal-Wallis test, p<0.0001; WT+shCtrl vs WT+shRptor, p=0.2318; WT+shCtrl vs Tsc1-KO+shCtrl, p<0.0001; WT+shCtrl vs Tsc1-KO+shRptor, p<0.0001; Tsc1-KO+shCtrl vs Tsc1-KO+shRptor, p<0.0001; Dunn’s multiple comparison tests. n-q) Representative images of primary somatosensory cortex from mice of the indicated genotypes and treatments showing EYFP (green) and myelin basic protein (MBP, magenta) immunostaining. Scale bars=1 mm. Str=striatum. Right panels show zoomed-in images of MBP immunostaining within the boxed cortical regions; scale bars=250 μm r) Mean +/- SEM bulk MBP fluorescence intensity in the boxed cortical regions shown in panels n-q for the indicated genotypes. n=6 animals per condition. Kruskal-Wallis, p=0.0114; WT+shCtrl vs WT+shRptor, p>0.9999; WT+shCtrl vs Tsc1-KO+shCtrl, **p=0.0077; WT+shCtrl vs Tsc1-KO+shRptor, p=0.4833; Tsc1-KO+shCtrl vs Tsc1-KO+shRptor, p=0.4833; Dunn’s multiple comparison tests. ns=non-significant. See also Supplementary Figures 9 and 10.

Decreased myelination in Tsc1-cKO mouse models can arise directly from changes in oligodendrocytes^61, 62^ or can occur via a non-cell-autonomous mechanism due to altered neuronal signaling^53, 63^. In *Tsc1^fl/fl^;Syn-Cre^+^* mice, there is impaired myelination due to increased secretion of connective tissue growth factor from Tsc1-cKO neurons, which negatively regulates oligodendrocyte development^63^. In Emx1-Cre mice, Cre is expressed primarily in neurons, although some glial cells can exhibit Cre expression^49, 51^. We tested whether shRptor injection into the forebrain of Tsc1-cKO mice could rescue decreased myelination. We found that shRptor boosted bulk cortical MBP fluorescence in Tsc1-cKO mice to WT levels (Fig. 7n-r). Since *Rptor* deletion from oligodendrocytes leads to reduced myelination^61, 64^, the increased MBP expression observed here was likely a result of Raptor reduction in Tsc1-cKO neurons, which led to a non-cell-autonomous increase in cortical myelination.

## Discussion

The goal of this study was to test whether genetic downregulation of mTORC1 or mTORC2 activity could be used to improve brain phenotypes in mouse models of TSC. We found that mTORC1 suppression via Raptor reduction was able to rebalance the activity of both mTOR complexes in the context of Tsc1 loss. While complete embryonic Raptor loss caused severe defects in forebrain development, 50% reduction of Raptor was sufficient to ameliorate several TSC-related phenotypes including signaling abnormalities, cellular alterations, network hyperactivity, and changes in brain architecture. Notably, mTORC2 suppression via loss of Rictor did not confer significant benefit in TSC mouse models. To extend the potential therapeutic relevance of our findings, we showed that postnatal downregulation of Raptor rescued Tsc1-cKO cellular phenotypes to a significant extent. Together, these data suggest that mTORC1 hyperactivity is primarily responsible for TSC-related brain phenotypes and that Raptor downregulation could be a potential therapeutic approach for the neurological presentations of TSC.

### The contribution of mTORC1 vs. mTORC2 to TSC-related brain phenotypes

The manifestations of TSC have traditionally been thought to arise from mTORC1 hyperactivity^2, 65^. However, there are several lines of evidence suggesting that mTORC2 may also be involved. First, rapamycin and its derivatives, which are used to treat TSC, not only strongly inhibit the p70S6K arm of mTORC1 signaling but also suppress mTORC2 activity in neurons^32^ (see Fig. 1). Second, a recent study in mice lacking *Pten*, an upstream negative regulator of mTORC1, showed that disease-related phenotypes could be prevented by inhibition of mTORC2 but not mTORC1^38^. Third, several different mouse models of TSC exhibit a defect in mGluR-dependent LTD^40–43^. It was recently shown that mTORC2 is the most relevant complex that regulates this form of hippocampal synaptic plasticity, which was not affected by deletion of *Rptor*^39^. Taken together, this leaves open the possibility that mTORC2 may be a relevant therapeutic target for TSC.

Here we deleted *Rptor* or *Rictor* in mouse models of TSC to determine whether suppression of mTORC1 or mTORC2 could improve disease-related phenotypes. We found that while reduction of Raptor levels ameliorated multiple TSC-related phenotypes, loss of Rictor did not significantly improve mTOR signaling abnormalities, neuronal hypertrophy, physical development, or premature mortality of Tsc1-cKO mice. This indicates that while reductions in Pten or the Tsc1/2 complex both increase mTORC1 signaling, the differences in how loss of these proteins affects other parts of the PI3K/Akt/mTOR signaling network need to be considered when designing a therapeutic strategy. In the case of *Pten* deletion, Akt, mTORC1 and mTORC2 signaling are all upregulated since Pten normally suppresses PI3K-dependent activation of Akt upstream of mTORC1 and mTORC2^38, 66–69^. Disruption of the Tsc1/2 complex also leads to increased mTORC1 signaling, which is due to loss of its GAP activity towards Rheb, a direct activator of mTORC1^70^. However, in contrast to Pten, loss of Tsc1/2 leads to decreased mTORC2-dependent phosphorylation of Akt^50, 61, 71, 72^.

Here we observed an interesting relationship between mTORC1 and mTORC2 signaling in neurons, indicating significant crosstalk between the two complexes. We found that while chronic rapamycin treatment suppressed both mTORC1 and mTORC2 signaling, *Rptor* deletion led to increased Akt phosphorylation at the mTORC2 site, in both WT and Tsc1-cKO neurons, suggesting enhanced mTORC2 activity. A similar effect has been reported in oligodendrocytes with conditional deletion of *Rptor* and in cancer cell lines with Raptor siRNA^61, 73^. In addition, Chen et al. showed that concomitant deletion of *Rptor* and *Pten* from forebrain neurons led to a stronger upregulation of p-Akt than observed with loss of *Pten* alone^38^. This phenotype is opposite of what we observed with Tsc1 loss, where mTORC2 signaling was reduced. There are several possible mechanisms whereby this crosstalk between mTORC1 and mTORC2 may occur. First, in conditions of high mTORC1 signaling, there may be enhanced negative feedback to upstream regulators of Akt, which would lead to reduced Akt phosphorylation^37, 72, 74, 75^.

Second, it has been shown in other systems that p70S6K1, a direct target of mTORC1, can phosphorylate two mTORC2 components, Rictor and mSin1, and inhibit mTORC2-dependent Akt phosphorylation^76–78^. In this case, Raptor reduction and reduced mTORC1 activity would lead to decreased Rictor phosphorylation, thereby boosting mTORC2-dependent phosphorylation of Akt. Third, in the case of Raptor reduction, it is possible that limited Raptor protein availability leads to decreased formation of mTORC1, which in turn may free up more mTOR protein to form mTORC2, leading to its enhanced activity. Further work is needed to clarify the mechanistic interactions between mTORC1 and mTORC2 signaling in neurons. Regardless of the mechanism, our data show that partial reduction of Raptor alone, in the absence of direct manipulations to mTORC2, can rebalance the signaling of both mTOR complexes in Tsc1 KO neurons by constraining mTORC1 signaling and releasing inhibition of mTORC2.

### The consequences of heterozygous versus homozygous loss of *Tsc1*

The mouse model of TSC used here has embryonic deletion of *Tsc1* from most forebrain excitatory neurons. This model recapitulates several disease-relevant phenotypes including spontaneous seizures, abnormalities in brain anatomy, neuronal hypertrophy, hypomyelination and astrogliosis^50, 51^. However, the phenotypes are generally more severe than in individuals with TSC as *Tsc1^fl/fl^;Emx1-Cre^+^* mice also have reduced body weight, fail to thrive, and exhibit premature mortality in the first few weeks of life. It is important to note that individuals with TSC generally carry germline heterozygous mutations in either the *TSC1* or *TSC2* genes, although somatic mosaicism can also occur^79–81^. The prevailing model is that during embryonic brain development, somatic second-hit mutations cause loss of heterozygosity and mTORC1 hyperactivity, leading to the formation of cortical malformations, which can become seizure foci^65, 82^. In support of this model, we found that loss of one copy of *Tsc1* from excitatory forebrain neurons was not sufficient to induce seizures, consistent with previous reports^83, 84^. To further investigate this, we generated primary hippocampal cultures from Tsc1-cHet mice and found that they did not exhibit spontaneous hyperactivity or mTORC1 activation. Previous reports have demonstrated that heterozygous loss of *Tsc1* or *Tsc2* from other neuronal types can induce synaptic and behavioral changes^42, 83–86^. Therefore, further work is needed to clarify which TSC-related phenotypes may be driven by haploinsufficiency and which require complete disruption of TSC1/2 complex activity.

### Importance of the developmental timing of mTOR perturbations

Balanced mTOR signaling is crucial for proper embryonic brain development^50, 52, 87–89^. In line with this, we found that embryonic mTORC1 suppression via homozygous deletion of *Rptor* from forebrain excitatory neurons led to stunted body and brain development. In particular, Raptor-cKO mice had a severely underdeveloped cortex and failed to form a discernable hippocampal structure, which occurred independent of *Tsc1* genotype. These results are consistent with a prior study that disrupted Raptor expression broadly in neuronal progenitors (using *Rptor^fl/fl^;Nestin-Cre^+^* mice), which led to microcephaly and perinatal mortality of Raptor-cKO mice^88^. Despite significant deficits in forebrain development, here we found that *Rptor^fl/fl^;Emx1-Cre^+^* mice had remarkably normal survival, with the majority of Raptor-cKO mice surviving into adulthood (> P150). Postnatal disruption of Raptor has also been investigated using *Rptor^fl/fl^;CamkIIa-Cre^+^* mice. In this model, Raptor-cKO mice had fully developed hippocampal and cortical structures, however, these brain regions were smaller compared to WT littermates^90^. Together these studies show that the developmental timing and number or type of cells impacted by mTORC1 alterations defines the severity of the phenotypes.

### Therapeutic considerations

One of the goals of this study was to test whether genetic suppression of mTORC1 could have beneficial outcomes in mouse models of TSC and avoid side-effects associated with rapalog treatment, which strongly inhibits both mTORC1 and mTORC2 signaling. While mTORC2/Akt suppression might be beneficial for anti-tumor drugs, suppression of Akt in the brain is associated with neuronal cell death^91^ and has been implicated in synaptic alterations underlying bipolar disorder^92, 93^. Here we found that it was possible to titrate Raptor protein levels to achieve near complete normalization of mTOR signaling in Tsc1-cKO neurons, as measured by p-S6 and p-Akt. Importantly, reduction of Raptor did not strongly suppress mTORC2, as chronic rapamycin does, and in fact boosted mTORC2-dependent phosphorylation of Akt and normalized it in the context of Tsc1 loss. However, disruption of Raptor showed a similar substrate bias as rapamycin, in which the p70S6K branch of mTORC1 signaling was preferentially suppressed with less impact on 4E-BP1 phosphorylation. Despite this, reduction of Raptor was sufficient to improve neuronal hypertrophy, network excitability, brain architecture, myelination, and survival in Tsc1-cKO mice. Therefore, genetic Raptor reduction is a promising therapeutic approach. Compared to systemic administration of small molecule drugs, a gene-based therapy has the advantage of potentially being targetable to specific tissues or cell types, which would avoid on-target, off-tissue side-effects that are a limitation to the chronic use of rapalogs^15^.

The Raptor manipulations tested here were consistently effective across multiple functional and anatomical read-outs both *in vitro* and *in vivo*. However, they did not provide complete prevention or rescue of all TSC-related phenotypes. In the genetic prevention experiments, this is likely because we could maximally achieve 50% loss of Raptor with heterozygous deletion, whereas our biochemical data suggest that ∼80% loss would be most beneficial. With Raptor shRNA, we were able to achieve 60-70% Raptor downregulation in Tsc1-cKO neurons, which was sufficient to improve cellular phenotypes, however stronger suppression may show even more benefit. That said, even a partial rescue can have overall beneficial outcomes, as was recently shown in Tsc1-cKO dopamine neurons. In *Tsc1^fl/fl^;DAT-Cre^+^* mice, heterozygous, but not homozygous, deletion of *Rptor* prevented functional deficits in dopamine release and cognitive flexibility, despite not rescuing somatic hypertrophy^94^. Importantly, postnatal reduction of Raptor should avoid the severe developmental impairments caused by embryonic mTORC1 suppression. Given our findings of a non-linear relationship between Raptor protein levels and mTORC1 signaling, it would be interesting to test other strategies for fine-tuning Raptor levels, for example using antisense oligonucleotides or CRISPR/Cas-based approaches, to achieve the optimal level and developmental timing of Raptor reduction. In addition, it would be worthwhile testing Raptor reduction in a mouse model with milder or later onset phenotypes to allow more time for therapeutic intervention.

In summary, our study confirms the importance of mTORC1 hyperactivity as a key driver of TSC-related brain phenotypes in mouse models and highlights Raptor as a relevant therapeutic target that has potential advantages over current pharmacologic approaches.

## Materials and Methods

### Mice

All animal procedures were carried out in accordance with protocols approved by the University of California, Berkeley Institutional Animal Care and Use Committee (IACUC). Mice of both sexes were used, and the ages are indicated in the methods for each experiment. Mice were on a mixed genetic background. Mice were housed with same-sex littermates in groups of 5–6 animals per cage and kept on a regular 12 h light/dark with ad libitum access to standard chow and water. Mice used for shRNA experiments were housed on an inverse 12 h light/dark cycle. Mouse genotypes were confirmed by PCR using genomic DNA obtained from tail samples. See Supplementary Table 4 for a list of the transgenic mouse lines used and genotyping primers.

### Primary hippocampal cultures

Dissociated hippocampal cultures were prepared from postnatal day 0-1 (P0-1) mice using standard protocols. Briefly, hippocampi from 2-3 pups (floxed mouse lines, Figs. 1-3 and Supplementary Figs. 1-3) or from single pups (Emx1-Cre mouse lines, Fig. 7 and Supplementary Figs. 6-9) were dissected on ice. The tissue was dissociated using 34.4 μg/ml papain in dissociation media (HBSS Ca^2+^, Mg^2+^ free, 1 mM sodium pyruvate, 0.1% D-glucose, 10 mM HEPES buffer) and incubated for 3 min at 37°C. Tissue digestion was stopped by incubation in trypsin inhibitor (1 mg/ml) in dissociation media at 37°C for 4 min. After trypsin inhibition, dissociation media was carefully removed and the tissue was gently manually triturated in 5 ml plating media (MEM, 10% FBS, 0.45% D-Glucose, 1 mM sodium pyruvate, 1 mM L-glutamine). Cell density was counted using a TC10 Automated cell counter (Bio-Rad) and ∼2-2.25 × 10^5^ neurons were plated for each experiment. For western blotting and Ca^2+^ imaging experiments, neurons were plated onto 24-well plates pre-coated with Poly-D-Lysine (PDL) (Corning, Cat # 08774271). For immunocytochemistry, neurons were plated onto 12 mm glass coverslips precoated overnight at room temperature (RT) with 0.5 mg/ml PDL in 0.1 M borate buffer (pH 8.5). On the plating day, the coverslips were rinsed 4 times with sterile water and then coated with 20 μg/ml laminin (GIBCO, 23017015) in 1x PBS for ∼1.5h at 37°C. Subsequently, coverslips were rinsed 3 times with sterile water, and 400 μl of plating media were added prior to adding the dissociated neurons. For all cultures, plating media was removed after 3 h and 900 μl maintenance media (Neurobasal media (Fisher Scientific # 21103-049) with 2 mM glutamine, pen/strep, and B-27 supplement (Invitrogen # 17504-044)) were added per well. After 4 days *in vitro* (DIV 4), 1 μM Cytosine β-D-arabinofuranoside (Sigma-Aldrich # C6645) was added to prevent glial proliferation. Cultures were maintained in maintenance media for 14 - 18 days with partial media changes every 4 days.

### Adeno-associated virus (AAV) transduction of primary cultures

For hippocampal culture experiments, AAVs were added on DIV 2, except for the shRNA experiments in which AAVs were added on DIV 1. Amounts of AAVs were chosen after titration experiments for each virus to accomplish either maximum or sparse transduction efficiency while maintaining low toxicity levels. In primary cultures from *Tsc1^fl/fl^*, *Tsc1^fl/fl^;Rptor^fl/fl^*, *Tsc1^fl/fl^;Rptor^wt/fl^* and *Tsc1^fl/fl^;Rictor^fl/fl^* mice that were used for western blotting experiments, we aimed for >95% transduction efficiency using AAV1 human Synapsin 1 (*SYN1,* “hSyn”) promoter-driven Cre-GFP or GFP to generate mutant and control cultures, respectively. For immunocytochemistry experiments, we aimed for sparse transduction to resolve individual neurons using AAV1-CBA-mCherry-Cre (nuclear localized), AAV9-CAG-FLEX-tdTomato, and AAV5-hSyn-GFP. To determine the percentage of Cre-expressing neurons in primary hippocampal cultures from Emx1-Cre-positive pups we used AAV1-CAG-FLEX-GFP at a high titer (see Supplementary Fig 6a). For calcium imaging experiments, we transduced neurons with AAV1-hSyn-jRGECO1a aiming for >95% transduction efficiency. For shRNA experiments, we transduced neurons with AAV9-hU6-shRNA-EYFP on DIV 1 and achieved 85-90% transduction. See Supplementary Table 5 for the list of viruses, source, titer and number of viral genomes (vg) used.

### Protein extraction and western blot analysis

Hippocampal cultures were harvested on DIV 14 -18. Neurobasal media was aspirated from one well at a time, wells were quickly rinsed with ice cold 1x PBS with Ca^2+^/Mg^2+^ and then 75 μl of lysis buffer were added (lysis buffer: 2 mM EDTA (Sigma: E5134), 2 mM EGTA (Sigma: E3889), 1% Triton-X (Sigma: T8787), and 0.5% SDS (Sigma: 71736) in 1× PBS with Halt phosphatase inhibitor cocktail (ThermoFisher: PI78420) and Complete mini EDTA-free protease inhibitor cocktail (Roche: 4693159001)). Wells were thoroughly scraped, and lysates were collected and sonicated for 5 sec. Total protein was determined by BCA assay (ThermoFisher: PI23227) and 10 μg of protein in 1X Laemmli sample buffer (Bio-Rad:161-0747) were loaded onto 4-15% Criterion TGX gels (Bio-Rad: 5671084). Proteins were transferred overnight at low voltage to PVDF membranes (Bio-Rad: 1620177), blocked in 5% milk in 1x TBS-Tween for one hour at RT, and incubated with primary antibodies diluted in 5% milk in 1x TBS-Tween overnight at 4°C. The following day, membranes were washed 3 x 10 min in 1x TBS-Tween and incubated with HRP-conjugated secondary antibodies (1:5000) for one hour at RT, washed 6 x 10 min in 1x TBS-Tween, incubated with chemiluminescence substrate (Perkin-Elmer: NEL105001EA) and developed on GE Amersham Hyperfilm ECL (VWR: 95017-661). Membranes were stripped by two 7 min incubations in stripping buffer (6 M guanidine hydrochloride (Sigma: G3272) with 1:150 β-mercaptoethanol) with shaking followed by four 2 min washes in 1x TBS with 0.05% NP-40 to re-blot on subsequent days. Bands were quantified by densitometry using ImageJ software (NIH). Phospho-proteins were normalized to their respective total proteins. Histone-3 was used as a loading control for every experiment. Antibody vendors, catalog numbers, and dilutions are listed in Supplementary Table 6.

### Immunocytochemistry

On DIV 14, Neurobasal media was carefully aspirated from all wells and coverslips were briefly rinsed with ice cold 1× PBS with Ca^2+^/Mg^2+^ followed by fixation in freshly made 4% PFA (Electron Microscopy Sciences: 15713) in 1x PBS for 10 min. PFA was then removed and the coverslips were washed for 5 min with 1x PBS three times. Coverslips were blocked for one hour at RT in buffer containing 5% normal goat serum (NGS) (ThermoFisher: PCN5000) and 0.3% Triton-X in 1x PBS and incubated in primary antibodies in antibody dilution buffer (1% BSA and 0.3% Triton-X in 1x PBS) overnight at 4°C. The following day, coverslips were washed three times for 5 min in 1x PBS, incubated with secondary antibodies in antibody dilution buffer (1:500) for one hour at RT and washed three times for 5 min in 1x PBS. Coverslips were mounted onto slides with ProLong Gold Antifade mountant with or without DAPI (ThermoFisher: P36934 or P36935) and allowed to set for one day before imaging. For soma size quantification, neuronal cultures from the floxed mouse lines were imaged on an FV1000 Olympus FluoView confocal microscope with a 20x objective. Neuronal cultures from the Emx1-Cre lines were imaged with a Hamamatsu Orca-er digital camera with a 10× objective and Micro-Manager 1.4 software. Soma area was quantified by manually tracing neuronal cell bodies using ImageJ software. Antibody vendors, catalog numbers, and dilutions are listed in Supplementary Table 6.

### Rapamycin treatment *in vitro*

Primary hippocampal cultures were treated chronically for 4 days with rapamycin from DIV 10-14. A stock solution of 0.5 mM rapamycin (LC Laboratories: R-5000) was prepared in ethanol and stored at −20°C. Rapamycin stock was diluted in Neurobasal media 1:100 prior to use and then added to a final concentration of 50 nM. Rapamycin was first added on DIV 10 and in the final media change on DIV 12.

### Survival and body weight monitoring

Breeder cages from the *Tsc1;Rptor;Emx1-Cre* and *Tsc1;Rictor;Emx1-Cre* lines were monitored daily with minimal disturbance. The birth date of each litter was noted, and the number of pups was recorded. At P11, pups were genotyped, and their body weight was measured. Weight was measured every two days from P11-P21 and every 5 days from P21-P40. Each pup that was found dead in the cage was immediately removed and re-genotyped for confirmation. Mice were handled minimally and with care to reduce stressors that could promote seizures.

### Perfusion and immunohistochemistry

P14-P15 mice were deeply anesthetized by isoflurane and transcardially perfused with ice-cold 1x PBS, followed by 4% PFA solution (Electron Microscopy Sciences: 15713) in 1x PBS using a peristaltic pump (Instech). The brains were removed and post-fixed by immersion in 4% PFA in 1x PBS overnight at 4 °C. Brains were then suspended in 30% sucrose in 0.1 M PB solution at 4°C until processed. Brains were sectioned coronally at 30 μm on a freezing microtome (American Optical AO 860).

Free-floating brain sections were batch processed to include matched control and experimental samples. With gentle shaking, sections were washed 3x 15 min in 1x PBS, followed by 1 h incubation at RT with BlockAid blocking solution (Life Tech: B10710). Primary antibodies were diluted in 1x PBS-T (PBS + 0.25% Triton-X-100) and applied for 48 h at 4 °C. Sections were washed with cold 1x PBS 3 x 15 min and incubated for 1h at RT with secondary antibodies diluted 1:500 in 1x PBS-T. Sections were then washed in cold 1x PBS 5 x 15 min, mounted on SuperFrost slides (VWR: 48311-703), and coverslipped with Vectashield hard-set mounting media with DAPI (Vector Labs: H-1500). See Supplementary Table 6 for a list of antibodies and dilutions used for immunohistochemistry.

### Confocal microscopy and image analysis

Images of brain sections were taken on either a Zeiss LSM 710 AxioObserver with 10x or 20x objectives, or an FV3000 Olympus FluoView confocal microscope with 4x or 20x objectives. For all quantitative comparison experiments, the same microscope and acquisition settings were used for each image and samples were processed in batches to include matched control and experimental samples. All images were processed using ImageJ software.

To quantify p-S6 levels and soma area, regions of interest (ROIs) were manually drawn around neuronal bodies on max-projected Z-stack images. The location and the size of the brain region selected for analysis was consistent across genotypes for each experiment. For the cortex, a region within the somatosensory area spanning all layers was selected and ∼200 neurons per section were traced. For the CA1, ∼70 neurons per section were traced and analyzed. For the dentate gyrus, ∼50 neurons per section from a sub-region of the suprapyramidal blade were analyzed per animal. To quantify ectopic cells above CA1, the number of cells with a strong p-S6 signal within a 249 x 83 μm ROI between the CA1 pyramidal layer and white matter were counted.

For myelin basic protein (MBP) analysis, the average bulk fluorescence intensity was calculated from a ∼468 x 468 μm sized ROI from a region spanning the retrosplenial area and secondary motor area drawn on max-projected Z-stack images. To examine GFAP expression in the cortex, a line was drawn across the cortical plate spanning from layer I to the border with the white matter. Mean GFAP fluorescence along the line was plotted using plot profile function in ImageJ. To examine GFAP expression in CA1, bulk average fluorescence intensity was calculated from a sub-region of CA1 approximately 232 x 81 μm.

For animals that received intracranial injections of AAV9-shRNA-EYFP bilaterally, neurons from both hemispheres were analyzed and pooled to account for potential differences in the injection efficiency. To quantify p-S6 levels, ∼ 185 EYFP+ neurons were manually outlined within the primary somatosensory area including all cortical layers. For the hippocampus, ∼85 neurons in CA1 and ∼65 neurons in DG from a middle region (along the anterior/posterior axis) of the hippocampus were analyzed per animal. To quantify MBP, two ∼621.5 x 621.5 μm ROIs (one per hemisphere) in the primary somatosensory cortex were drawn and bulk MBP fluorescence intensity was quantified and averaged per mouse. All ROIs were drawn on max-projected Z-stack images using ImageJ.

### Calcium Imaging

Primary hippocampal cultures from P0-1 *Tsc1;Rptor;Emx1-Cre* mice of different genotypes were plated onto 24 well plates pre-coated with PDL (Corning, Cat # 08774271). On DIV 2, cultures were transduced with AAV1-jRGECO1a (Supplementary Table 5) and maintained for 12 days in Neurobasal media. On DIV 14, neurons were imaged on an AxioObserver.A1 (Zeiss) inverted microscope using a 10x Zeiss A-Plan objective (Numerical Aperture: 0.25) with wide field fluorescence illumination (X-Cite series 120Q, Lumen Dynamics). Images were taken at 8.91 Hz with a Hamamatsu Orca-er digital camera and Micro-Manager 1.4 software. A single field of view (FOV) was imaged from at least 2-3 individual wells per culture (prepped from 1 pup) and approximately 40 neurons were randomly selected and analyzed. Before proceeding to the analysis, we verified that the neurons selected were active at least once during the recording period. At least 3 mice per genotype were examined from at least 3 different litters.

### Calcium imaging analysis

Data analysis was performed using ImageJ 1.53c and custom programs written in Matlab 2020a.

#### Pre-processing

Circular ROIs corresponding to neuronal somata, were drawn manually in ImageJ on mean intensity projection images of the recorded FOV. Forty ROIs were drawn per FOV, beginning in the upper left quadrant of the image, and extending outward as necessary (to the right, bottom left, and bottom right quadrants, respectively), and were imported into Matlab for further analysis. Movies were motion corrected using the normcorre function^95^, then normalized with respect to baseline, taken to be the minimum intensity projection of the FOV, to generate a ΔF/F movie. It was necessary to use the minimum projection as Tsc1-cKO neurons exhibited high Ca^2+^ activity and thus contained few baseline frames. Ca^2+^ traces were extracted as ΔF/F by computing the mean fluorescence within each ROI at each movie frame.

#### Single event analysis

Individual Ca^2+^ transients were detected by first filtering the ΔF/F traces with a four-frame moving mean, then using Matlab’s findpeaks function to identify peaks in the ΔF/F trace. Event amplitude was defined as the difference between the event’s peak ΔF/F and the minimum ΔF/F in the preceding inter-event-interval (see Supplementary Fig. 6c). Events with an amplitude < 0.5% were excluded from further analysis. Prior to measurement of event duration and area under the curve (AUC), Matlab’s msbackadj function was used to shift the 1st percentile of the ΔF/F trace within 15 second time windows to zero. This reduced the reliance of event AUC on preceding bouts of Ca^2+^ activity, which would otherwise contribute significantly to AUC values in Tsc1-cKO neurons due to their high frequency activity. Event initiation and termination were identified by finding the 0.5% ΔF/F threshold (see Supplementary Fig. 6d) crossing preceding and following the event peak. Event termination was alternatively identified when the Ca^2+^ decay following the event peak ΔF/F was interrupted by a ΔF/F increase >1%, indicating the initiation of another event. Events without a clear initiation or termination were excluded from further analysis. AUC was defined as the area under the ΔF/F trace during the event (see Supplementary Fig. 6d) and measured using trapezoidal numerical integration implemented by the trapz function in Matlab.

### Network event analysis

Network Ca^2+^ events were defined as time intervals over which more than 20% of neuronal ROIs in the imaged area were simultaneously active (activity for a single neuron was defined as ΔF/F ≥ 0.5%) (See gray highlighted zones in Fig. 6e-g and Supplementary Fig. 8c,d). We did not use a standard deviation-based threshold, as it would have selectively reduced event detection in Tsc1-cKO cultures, due to their persistent Ca^2+^ activity. Events with a duration < 2.5 seconds were excluded from further analysis. Cell participation in events was defined as the percentage of neurons that were active at any time during the event. Response amplitude within a single neuron during a network event was defined as the difference between the maximum ΔF/F during the event and the minimum ΔF/F in the preceding inter-event-interval. Area under the curve (AUC) was defined as the area under the ΔF/F trace occurring during the event.

### shRNA constructs

The AAV-Tet3-shRNA plasmid (Addgene plasmid # 85740) was used as a backbone to generate the AAV9-hU6-shRptor-EYFP construct. The restriction enzymes BamHI and XbaI were used to excise the Tet-3 shRNA sequence. The oligonucleotide sequence: 5’-GATCCGCCTCATCGTCAAGTCCTTCAAGAAGCTTGTTGAAGGACTTGACGATGAGGCTTTTTTT -3’ that contains the *Rptor* shRNA sequence^60^ flanked by restriction sites for BamHI and XbaI was subcloned into the plasmid backbone. AAV-hU6-shControl-EYFP (Addgene plasmid # 85741) that contains the 5’-GTTCAGATGTGCGGCGAGT-3’ shRNA sequence was used as a control. For large-scale isolation and purification of the plasmids, DH5α NEB competent cells (New England Biolabs #C2987H) were transformed and Endofree Megaprep (Qiagen # 12381) was performed to generate plasmids for high titer viral packaging.

### Intracranial neonatal mouse injections

Neonatal mice (P0) were cryo-anesthetized by placing on ice for ∼2-3 min. When the animal was fully anesthetized, confirmed by lack of movement, it was placed in a head mold. Each pup received a total of 500 nl of 4x diluted AAV9 (AAV9-hU6-shRptor-EYFP or AAV9-hU6-shControl-EYFP, see Supplementary Table 5 for titer information) spread across 4 injections (2 per hemisphere): two 150 nl injections were made into the cortex and two 100 nl injections were made into the dorsal hippocampus. Cortical injections were made halfway between bregma and lambda approximately 0.6 mm from the sagittal suture and 0.5-0.6 mm ventral to the surface of the skull. Hippocampal injections were made approximately 0.5 mm anterior to lambda with the injection sites ∼0.5 mm from the sagittal suture and 1 mm ventral to the surface of the skull.

### Statistical Analysis

Statistical analyses and graphing were performed using GraphPad Prism software (versions 7-9). All datasets were first analyzed using the D’Agostino and Pearson normality test, and then parametric or non-parametric two-tailed statistical tests were employed accordingly to determine significance. Normally distributed datasets were analyzed using Welch’s t-tests when comparing two groups or a one-way ANOVA with Holm-Sidak’s multiple comparison tests when comparing three or more groups. Datasets that did not pass the normality test were analyzed using a Mann-Whitney test when comparing two groups or the Kruskal–Wallis test with Dunn’s multiple comparisons tests when comparing three or more groups. Cumulative distributions were analyzed using Kolmogorov–Smirnov tests (when comparing two groups) or the Kruskal–Wallis test with Dunn’s multiple comparisons tests (when comparing three or more groups). Survival curves were analyzed using the Log-rank (Mantel-Cox) test. Regression models were analyzed either with Pearson correlation (linear regression) or Spearman correlation (non-linear regression). Significance was set as *p<0.05, **p<0.01, ***p<0.001, and ****p<0.0001. P values were corrected for multiple comparisons.

## Acknowledgements

This work was supported by grants from the Mary Elizabeth Rennie Foundation for Epilepsy Research. V.K. was supported by an Elizabeth Roboz Einstein fellowship and pre-doctoral NIH/NIGMS training grant T32 GM7232-40. H.S.B. is a Chan Zuckerberg Biohub Investigator and was supported by an Alfred P. Sloan fellowship. Confocal imaging experiments were conducted in part at the CRL Molecular Imaging Center supported by the Gordon and Betty Moore Foundation and Helen Wills Neuroscience Institute. We thank Holly Aaron and Feather Ives for microscopy training and assistance. We thank Dr. Marla Feller for her advice on the calcium imaging experiments. We thank Dr. Viviana Gradinaru and the Caltech CLOVER center for generating AAV9 viral particles and packaging the shRNA constructs. We thank Dr. Hillel Adesnik for providing the initial Emx1-Cre mice and the AAV-jRGECO1a virus. We would like to thank the members of the Bateup lab for their feedback on this work.

## Author Contributions

V.K. designed and executed all experiments and performed the initial analyses. V.K. wrote the initial draft of the manuscript and contributed to review and editing. F.C-H. advised on the design of the calcium imaging experiments, wrote the MATLAB analysis code, performed calcium imaging analysis, and contributed to review & editing. H.S.B. advised on the design and execution of the project, performed secondary analyses, contributed to writing and editing the manuscript, and obtained funding.

