## Supplementary Information for "Raptor downregulation rescues neuronal phenotypes in mouse models of Tuberous Sclerosis Complex"

**Supplementary Figure 1**

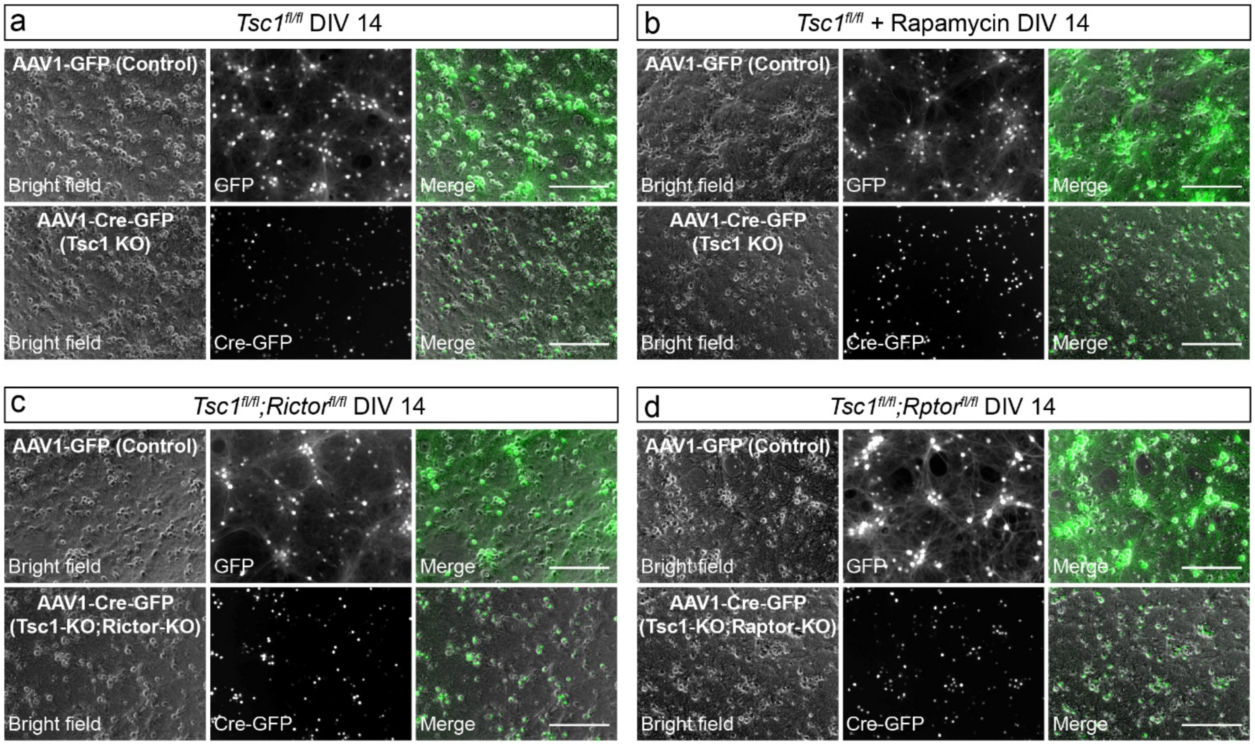

**Supplementary Figure 1. Example images of primary hippocampal cultures. Related to Figures 1-3.**

Representative bright-field and fluorescence images of primary hippocampal cultures from P0-P1 pups imaged on DIV 14. The cultures were transduced on DIV 2 with AAV-GFP (cytosolic expression) or AAV-Cre-GFP (nuclear localized).

a) *Tsc1*$^{fl/fl}$ cultures + GFP (Control, top panels) or Cre-GFP (Tsc1 KO, bottom panels)

b) *Tsc1*$^{fl/fl}$ cultures + GFP (Control, top panels) or Cre-GFP (Tsc1 KO, bottom panels) treated with rapamycin (50 nM) from DIV 10-14.

c) *Tsc1*$^{fl/fl}$*;Rictor*$^{fl/fl}$ cultures + GFP (Control, top panels) or Cre-GFP (Tsc1-KO;Rictor-KO, bottom panels).

d) *Tsc1*$^{fl/fl}$*;Rptor*$^{fl/fl}$ cultures + GFP (Control, top panels) or Cre-GFP (Tsc1-KO;Raptor-KO, bottom panels)

All scale bars=250 μm

**Supplementary Figure 2**

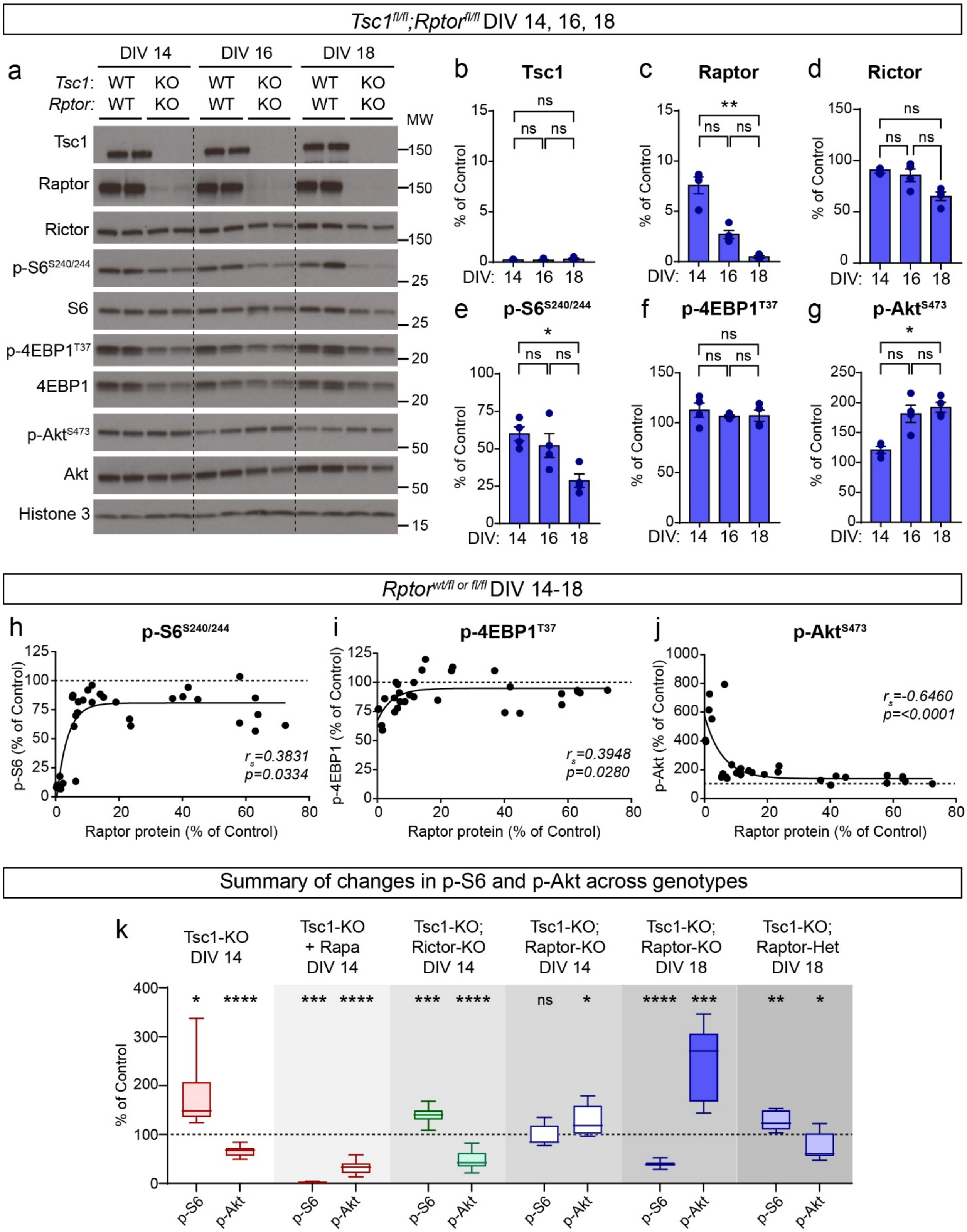

**Supplementary Figure 2. Raptor reduction affects both mTORC1 and mTORC2 signaling. Related to Figures 1-3.**

a) Representative western blots of lysates from *Tsc1$^{fl/fl}$;Rptor$^{fl/fl}$* hippocampal cultures treated with GFP (WT;WT) or Cre-GFP (KO;KO) on DIV 2 and collected at different time points (DIV 14, 16, and 18). MW indicates molecular weight. Two samples per genotype are shown; this experiment was replicated two times.

b-g) Bar graphs display western blot quantification (mean +/- SEM) for the indicated proteins. The data have been normalized to control values (WT;WT samples) for each DIV. Phospho-proteins were normalized to their respective total proteins. Dots represent data from individual culture wells. For all genotypes n=2 culture wells from 2 independent cultures; 2 mice per culture.

b) Tsc1, Kruskal-Wallis, p=0.5101; D14 vs D16, p>0.9999; D16 vs D18, p=0.7179; D14 vs D18, p>0.9999; Dunn's multiple comparisons tests. ns=non-significant.

c) Raptor, Kruskal-Wallis, p=0.0002; D14 vs D16, p=0.3500; D16 vs D18, p=0.3500; D14 vs D18, **p=0.0051; Dunn's multiple comparisons tests.

d) Rictor, Kruskal-Wallis, p=0.0263; D14 vs D16, p>0.9999; D16 vs D18, p=0.1184; D14 vs D18, p=0.0558 Dunn's multiple comparisons tests.

e) p-S6 240/244, Kruskal-Wallis, p=0.0194; D14 vs D16, p>0.9999; D16 vs D18, p=0.1873; D14 vs D18, *p=0.0324; Dunn's multiple comparisons tests.

f) p-4EBP1 T37, Kruskal-Wallis, p=0.7463; D14 vs D16, p>0.9999; D16 vs D18, p>0.9999; D14 vs D18, p>0.9999; Dunn's multiple comparisons tests.

g) p-Akt Ser473, Kruskal-Wallis, p=0.0132; D14 vs D16, p=0.0723; D16 vs D18, p>0.9999; D14 vs D18, *p=0.0427; Dunn's multiple comparisons tests.

h-j) Correlation of Raptor protein levels to p-S6 Ser240/244 (h), p-4EBP1 T37 (i), or p-Akt Ser473 (j) levels within each culture, expressed as a percentage of Control. Samples were pooled across hippocampal cultures from *Rptor$^{wt/fl\ or\ fl/fl}$* mice treated with AAV-GFP or AAV-Cre-GFP and harvested on different days (DIV 14-18) to generate a range of Raptor protein levels. Solid lines depict non-linear regression. Dashed lines represent control levels. Dots represent individual culture wells, n=31 culture wells. For panel h, r= 0.3831, p=0.0334, Spearman correlation. For panel i, r=0.3948, p =0.0280, Spearman correlation. For panel j, r= -0.6460, ****p<0.0001, Spearman correlation.

k) Box-and-whisker plots (min to max) displaying the p-S6 Ser240/244 and p-Akt Ser473 western blot results for the indicated conditions, expressed as a percentage of their respective controls. Data are replotted from Figs. 1-3. Dashed line indicates control levels.

**Supplementary Figure 3**

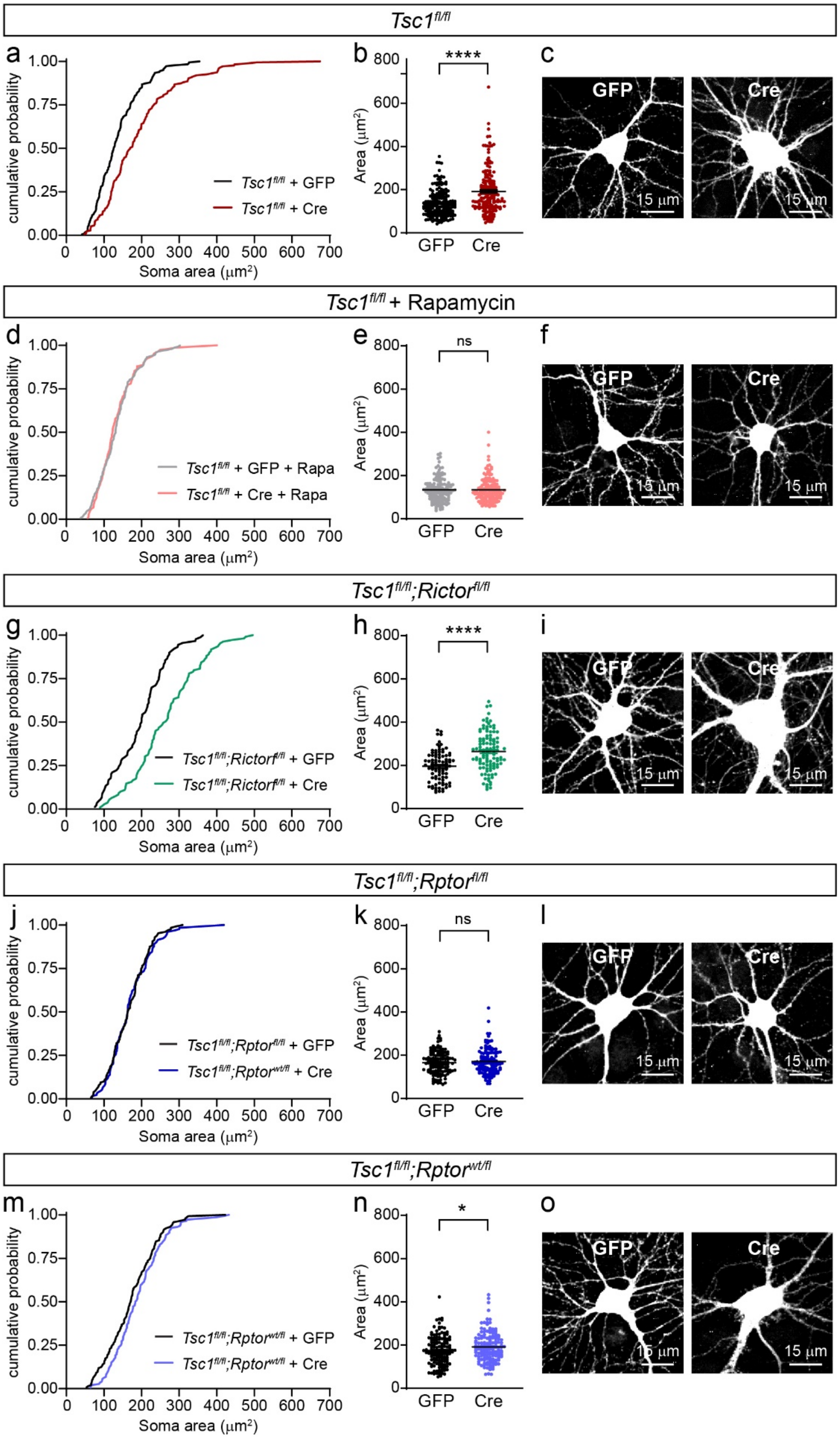

**Supplementary Figure 3. Hypertrophy of Tsc1-cKO neurons is prevented by genetic reduction of Raptor but not Rictor.**

**Related to Figure 3.**

a) Cumulative distributions of soma area for cultured *Tsc1*$^{fl/fl}$ neurons treated with AAV-GFP (control) or AAV-Cre-GFP (Tsc1-KO). n=182 GFP+ and 175 Cre+ neurons.

b) Scatter dot plot of the data in panel a. Black lines indicated mean +/- SEM. Mann-Whitney, ****p<0.0001.

c) Example fluorescence images of *Tsc1*$^{fl/fl}$ neurons expressing GFP (control) or Cre-dependent tdTomato (Tsc1-KO).

d) Cumulative distributions of soma area for cultured *Tsc1*$^{fl/fl}$ neurons treated with AAV-GFP (control) or AAV-Cre-GFP (Tsc1-KO). Cultures were treated with 50 nM rapamycin from DIV 10-14. n=147 GFP+ and 150 Cre+ neurons.

e) Scatter dot plot of the data in panel d. Black lines indicated mean +/- SEM. Mann-Whitney, p=0.7508.

f) Example fluorescence images of *Tsc1*$^{fl/fl}$ neurons treated with rapamycin expressing GFP (control) or Cre-dependent tdTomato (Tsc1-KO).

g) Cumulative distributions of soma area for cultured *Tsc1*$^{fl/fl}$*;Rictor*$^{fl/fl}$ neurons treated with AAV-GFP (control) or AAV-Cre-GFP (Tsc1-KO;Rictor-KO). n=90 GFP+ and 109 Cre+ neurons.

h) Scatter dot plot of the data in panel g. Black lines indicated mean +/- SEM. Welch’s t-test, ****p<0.0001.

i) Example fluorescence images of *Tsc1*$^{fl/fl}$*;Rictor*$^{fl/fl}$ neurons expressing GFP (control) or Cre-dependent tdTomato (Tsc1-KO;Rictor-KO).

j) Cumulative distributions of soma area for cultured *Tsc1*$^{fl/fl}$*;Rptor*$^{fl/fl}$ neurons treated with AAV-GFP (control) or AAV-Cre-GFP (Tsc1-KO;Raptor-KO). n=128 GFP+ and 130 Cre+ neurons.

k) Scatter dot plot of the data in panel j. Black lines indicated mean +/- SEM. Mann-Whitney, p=0.8062.

l) Example fluorescence images of *Tsc1*$^{fl/fl}$*;Rptor*$^{fl/fl}$ neurons expressing GFP (control) or Cre-dependent tdTomato (Tsc1-KO;Raptor-KO).

m) Cumulative distributions of soma area for cultured *Tsc1*$^{fl/fl}$*;Rptor*$^{wt/fl}$ neurons treated with AAV-GFP (control) or AAV-Cre-GFP (Tsc1-KO;Raptor-Het). n=144 GFP+ and 146 Cre+ neurons.

n) Scatter dot plot of the data in panel n. Black lines indicated mean +/- SEM. Mann-Whitney, *p=0.0452.

o) Example fluorescence images of *Tsc1*$^{fl/fl}$*;Rptor*$^{wt/fl}$ neurons expressing GFP (control) or Cre-dependent tdTomato (Tsc1-KO;Raptor-Het).

For all panels, cultures were harvested on DIV 14. Neurons were imaged from 4-5 culture wells from 3 independent culture preps, 2 pups per prep. ns=non-significant. Scale bars=15 μm.

**Supplementary Figure 4**

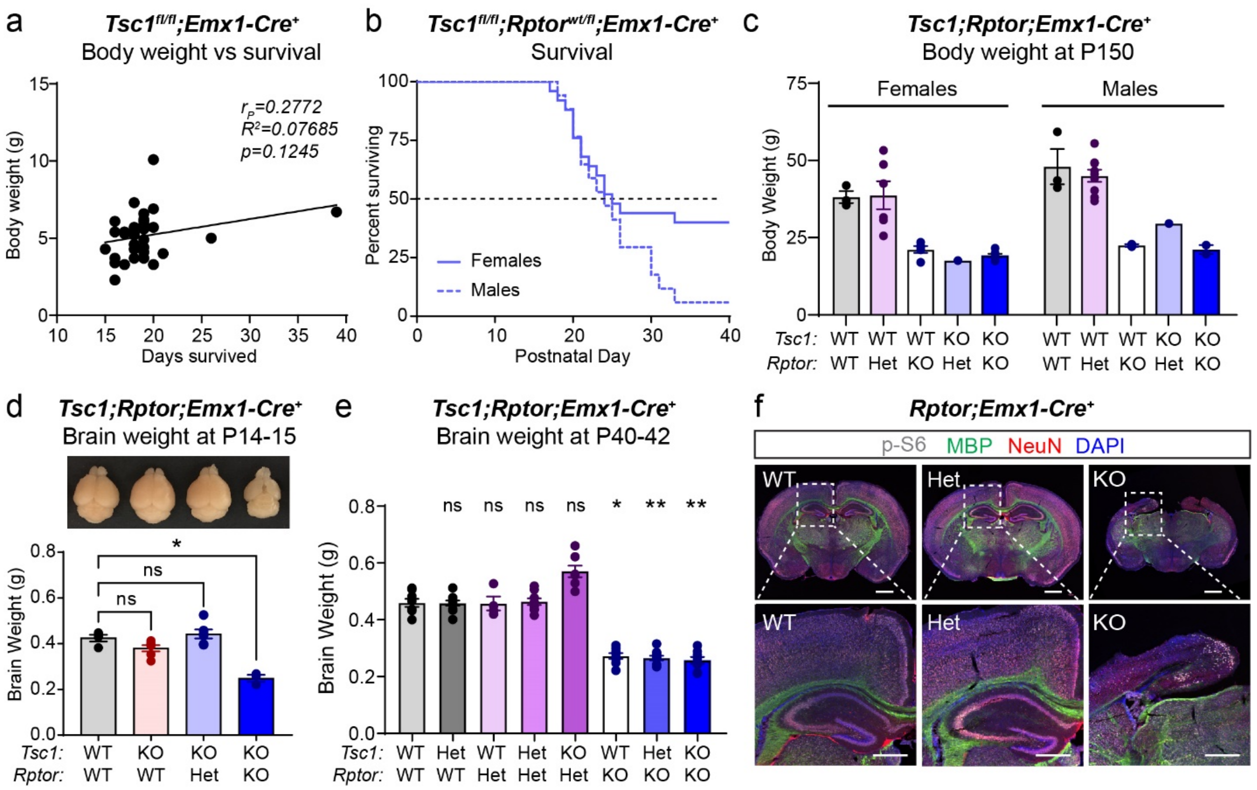

**Supplementary Figure 4. Raptor reduction impacts brain and body weight, survival, and forebrain development**

**Related to Figure 4.**

a) Scatterplot displaying the number of days *Tsc1fl/fl;Emx1-Cre+* (Tsc1-KO) mice survived versus their final recorded body weight. Dots represent individual mice, n=32. Tsc1-KO mice were pooled from the *Tsc1;Rptor;Emx1-Cre* and *Tsc1;Rictor;Emx1-Cre* lines. r= 0.2772, p=0.1245, Pearson correlation.

b) Survival analysis of *Tsc1fl/fl;Rptorwt/fl;Emx1-Cre+* mice by sex. n=17 male and 25 female mice. Log-rank (Mantel-Cox) test, p=0.1024. Dashed black line indicates 50% of the population surviving.

c) Mean +/- SEM body weight of *Tsc1;Rptor;Emx1-Cre* mice of the indicated sex and genotype on postnatal day 150. Dots represent individual mice. n=3 male and 3 female Tsc1-WT;Raptor-WT mice; 9 male and 6 female Tsc1-WT;Raptor-Het mice; 2 male and 5 female Tsc1-WT;Raptor-KO mice; 1 male and 1 female Tsc1-KO;Raptor-Het mouse; and 2 male and 7 female Tsc1-KO;Raptor-KO mice.

d) Top, representative whole brain images from mice of the genotypes indicated under the respective bar graph. Bottom, mean +/- SEM brain weight from P14-15 mice of the indicated genotypes. Dots represent individual mice. n=4 Tsc1-WT;Raptor-WT mice, 6 Tsc1-KO;Raptor-WT mice, 6 Tsc1-KO;Raptor-Het mice, and 3 Tsc1-KO;Raptor-KO mice. Kruskal-Wallis, p=0.0027; WT;WT vs KO;WT, p=0.5730; WT;WT vs KO;Het, p>0.9999; WT;WT vs KO;KO, *p=0.0371; Dunn's multiple comparison tests. ns=non-significant.
e) Mean +/- SEM brain weight of P40-42 *Tsc1;Rptor;Emx1-Cre* mice of the indicated genotypes. Dots represent individual mice. n=8 Tsc1-WT;Raptor-WT, 10 Tsc1-Het;Rptor-WT, 4 Tsc1-WT;Rptor-Het, 10 Tsc1-Het;Rptor-Het, 7 Tsc1-KO;Rptor-Het, 8 Tsc1-WT;Rptor-KO, 11 Tsc1-Het;Rptor-KO, and 8 Tsc1-KO;Rptor-KO mice. Kruskal-Wallis, p<0.0001. Dunn's multiple comparisons tests to WT;WT:  Het;WT, p>0.9999; WT;Het, p>0.9999; Het;Het, p>0.9999; KO;Het, p=0.4700; WT;KO, *p=0.0258; Het;KO, **p=0.0057; KO;KO, **p=0.0067.
f) Representative images of coronal brain sections showing p-S6 Ser240/244 (gray), myelin basic protein (MBP, green), and NeuN (red) immunostaining in P14 *Rptor*$^{wt/wt}$*;Emx1-Cre*$^{+}$ (WT), *Rptor*$^{wt/fl}$*;Emx1-Cre*$^{+}$ (Het) and *Rptor*$^{fl/fl}$*;Emx1-Cre*$^{+}$ mice (all *Tsc1*$^{wt/wt}$). DAPI staining is in blue. Scale bars=1 mm. Bottom panels show zoomed-in images of the hippocampal regions indicated by the dashed boxes. Scale bars=500 μm.

**Supplementary Figure 5**

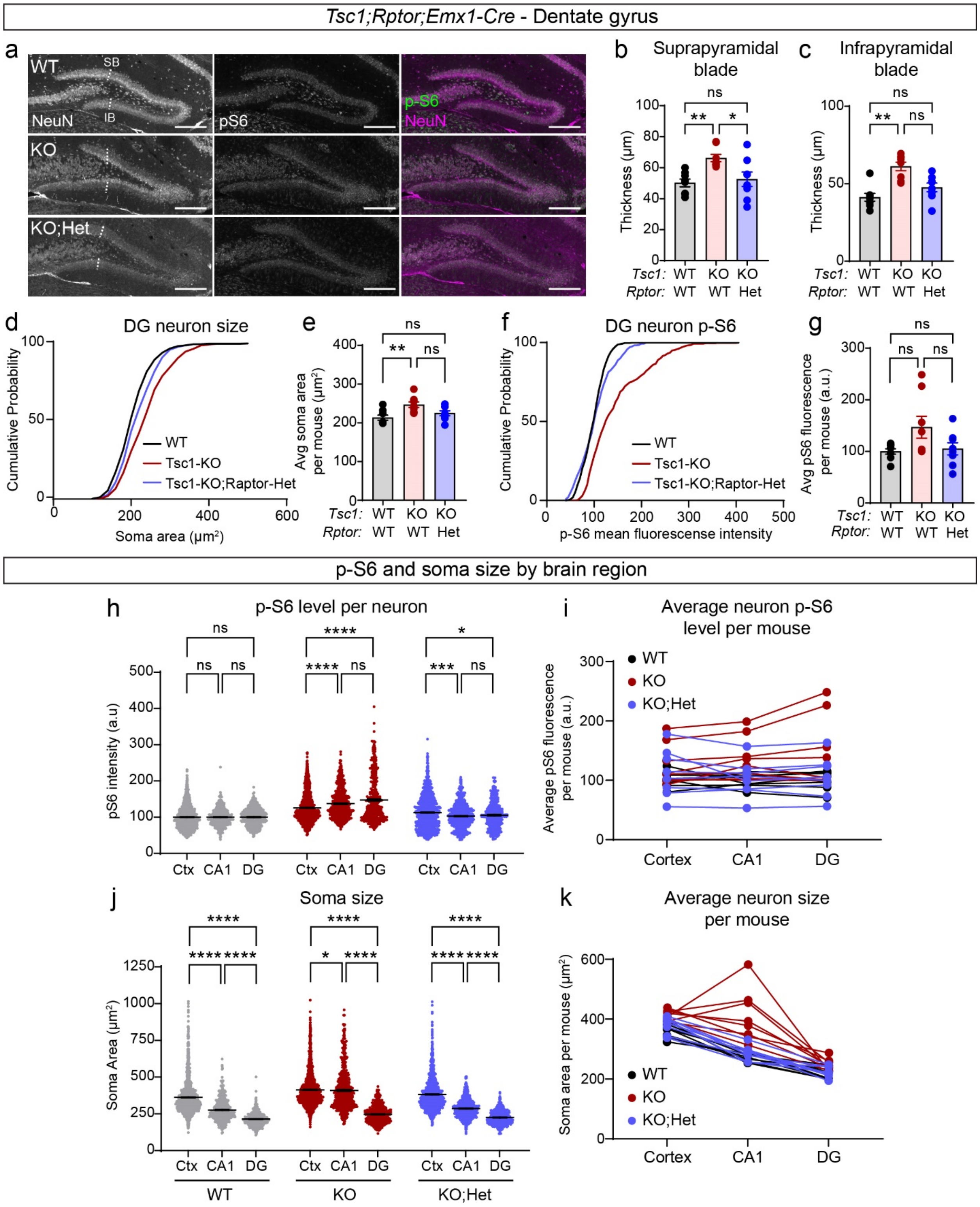

**Supplementary Figure 5. p-S6 and soma size exhibit brain region-specific differences. Related to Figure 5.**

a) Representative images from the dentate gyrus (DG) showing NeuN (left panels) and p-S6 Ser240/244 (middle panels) immunostaining in *Tsc1*$^{wt/wt}$*;Rptor*$^{wt/wt}$*;Emx1-Cre*$^{+}$ (WT), *Tsc1*$^{fl/fl}$*;Rptor*$^{wt/wt}$*;Emx1-Cre*$^{+}$ (KO), and *Tsc1*$^{fl/fl}$*;Rptor*$^{wt/fl}$*;Emx1-Cre*$^{+}$ (KO;Het) mice. Merged images on the right show NeuN in magenta and p-S6 in green. SB = Suprapyramidal blade, IB = Infrapyramidal blade. Lines denote the measurement of IB and SB thickness. Scale bars=250 µm

b) Mean +/- SEM thickness of the suprapyramidal blade for the indicated genotypes. Dots represent values from individual mice, n=8 mice per genotype. One-way ANOVA, p=0.0055, F(2, 21) = 6.745; WT vs KO, **p=0.0081; WT vs KO;Het, p=0.9473; KO vs KO;Het, *p=0.0253; Sidak's multiple comparisons tests. ns=non-significant.

c) Mean +/- SEM thickness of the infrapyramidal blade for the indicated genotypes. Dots represent values from individual mice, n=8 mice per genotype. Kruskal-Wallis, p=0.0019; WT vs KO, **p=0.0014; WT vs KO;Het, p=0.6093; KO vs KO;Het, p=0.0778; Sidak's multiple comparisons tests.

d) Cumulative distributions of dentate gyrus (DG) neuron soma area for the indicated genotypes. n=403 WT, 401 Tsc1-KO, and 400 Tsc1-KO;Rptor-Het neurons from 8 mice per genotype. Kruskal-Wallis, p<0.0001; WT vs KO, p<0.0001; WT vs KO;Het, p=0.0022; KO vs KO;Het,  p<0.0001; Dunn's multiple comparisons tests.

e) Mean +/- DG neuron soma area per mouse for the indicated genotypes. Dots represent values from individual mice, n=8 mice per genotype. One-way ANOVA, p=0.0074, F(2, 21) = 6.257; WT vs KO, **p=0.0067; WT vs KO;Het, p=0.5700; KO vs KO;Het, p=0.0951; Sidak's multiple comparisons tests.

f) Cumulative distributions of DG p-S6 levels per neuron for the indicated genotypes. n is the same as for panel d. Kruskal-Wallis, p<0.0001; WT vs KO, p<0.0001; WT vs KO;Het, p=0.4109; KO vs KO;Het, p<0.0001; Dunn's multiple comparisons tests.

g) Mean +/- SEM DG neuron p-S6 levels per mouse for the indicated genotypes. Dots represent values from individual mice, n=8 mice per genotype. One-way ANOVA, p=0.0591, F (2, 21) = 3.246; WT vs KO, p=0.0883; WT vs KO;Het, p=0.9923; KO vs KO;Het, p=0.1456; Sidak's multiple comparisons tests.

h) Scatter dot plots of p-S6 levels per neuron for the cortex (Ctx), CA1 and DG regions for WT (grey dots), KO (red dots) and KO;Het (blue dots) mice. Black lines indicate mean +/- SEM. n=1601 WT Ctx neurons, 1605 KO Ctx neurons, 1602 KO;Het Ctx neurons, 561 WT CA1 neurons, 561 KO CA1 neurons, 561 KO;Het CA1 neurons, 403 WT DG neurons, 401 KO DG neurons, and 400 KO;Het DG neurons from 8 mice per genotype. WT p-S6 levels: Kruskal-Wallis test, p=0.0669; Ctx vs CA1, p=0.1806; Ctx vs DG, p=0.2338; CA1 vs DG, p>0.9999; Dunn's multiple comparison tests. KO p-S6 levels: Kruskal-Wallis test, p<0.0001; Ctx vs CA1, ****p<0.0001; Ctx vs DG, ****p<0.0001; CA1 vs DG, p>0.9999; Dunn's multiple comparison tests. KO;Het p-S6 levels: Kruskal-Wallis test, p=0.0001; Ctx vs CA1, ***p=0.0002, Ctx vs DG, *p=0.0485, CA1 vs DG, p>0.9999, Dunn's multiple comparison tests.

i) Average p-S6 levels per mouse for neurons in the cortex, CA1 and DG regions for WT (black dots), KO (red dots) and KO;Het (blue dots) mice. Lines connect data points from the same mouse. n=8 mice per genotype.

j) Scatter dot plots of soma area for neurons in the Ctx, CA1 and DG regions for WT (grey dots), KO (red dots) and KO;Het (blue dots) mice. n is the same as in panel h. WT soma area: Kruskal-Wallis test, p<0.0001; Ctx vs CA1, ****p<0.0001; Ctx vs DG, ****p<0.0001, CA1 vs DG, ****p<0.0001; Dunn's multiple comparison tests. KO soma area: Kruskal-Wallis test, p<0.0001; Ctx vs CA1, *p=0.0299, Ctx vs DG, ****p<0.0001; CA1 vs DG, ****p<0.0001, Dunn's multiple comparison tests. KO;Het soma area: Kruskal-Wallis test, p<0.0001; Ctx vs CA1, p<0.0001; Ctx vs CA1, ****p<0.0001; Ctx vs DG, ****p<0.0001, CA1 vs DG, ****p<0.0001, Dunn's multiple comparison tests.

k) Average soma size per mouse for neurons in the cortex, CA1 and DG regions for WT (black dots), KO (red dots) and KO;Het (blue dots) mice. Lines connect data points from the same mouse. n=8 mice per genotype.

**Supplementary Figure 6**

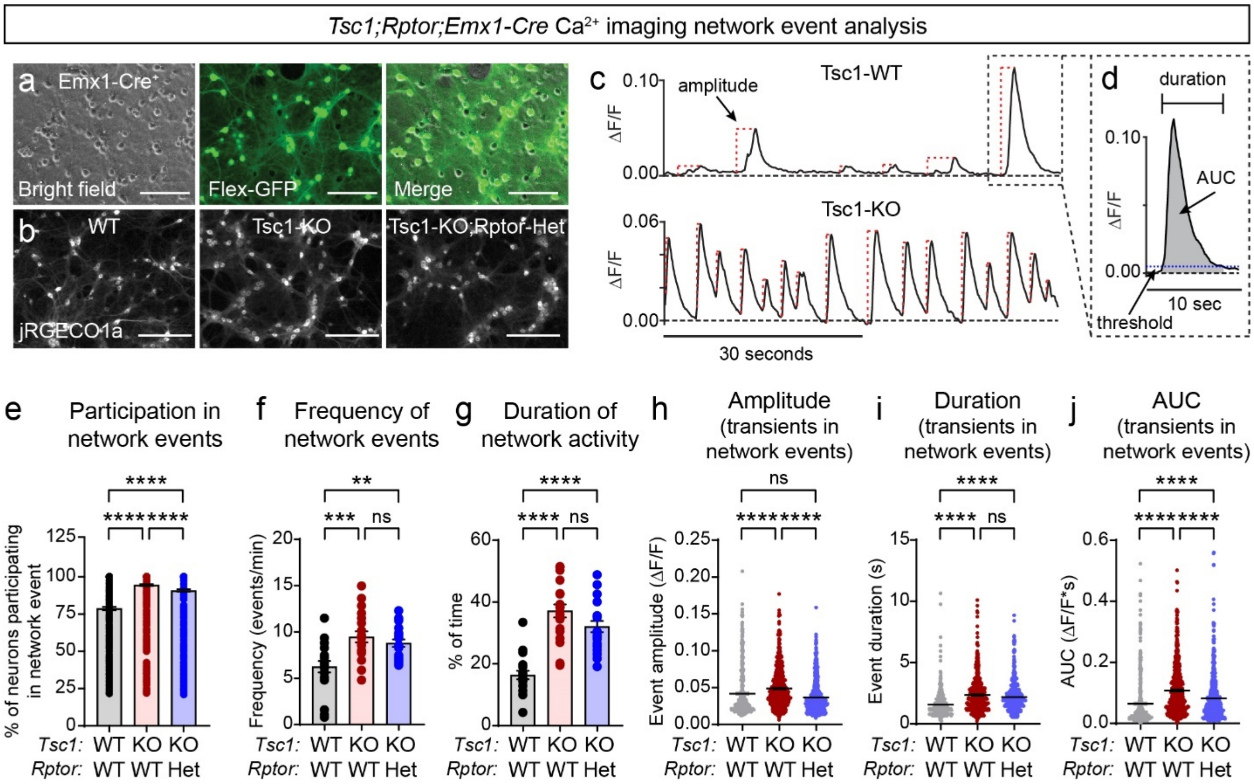

**Supplementary Figure 6. Heterozygous deletion of *Rptor* partially improves network hyperactivity of Tsc1-cKO hippocampal cultures.**

**Related to Figure 6.**

a) Representative images of a primary hippocampal culture from a *Tsc1$^{wt/wt}$;Emx1-Cre*$^{+}$ mouse transduced with a Cre-dependent GFP expressing virus (AAV-Flex-GFP). Bright field (left panel), GFP fluorescence (middle panel) and merged (right panel) images are shown. Scale bars=150 µm

b) Representative images of cultures from *Tsc1$^{wt/wt}$;Rptor$^{wt/wt}$;Emx1-Cre*$^{+}$ (WT), *Tsc1$^{fl/fl}$;Rptor$^{wt/wt}$;Emx1-Cre*$^{+}$ (KO), and *Tsc1$^{fl/fl}$;Rptor$^{wt/fl}$;Emx1-Cre*$^{+}$ (KO;Het) mice expressing jRGeco1a. Scale bars=250 µm.

c) Example of $Ca^{2+}$ imaging analysis showing individual $Ca^{2+}$ transients from a Tsc1-WT (top) and Tsc1-KO (bottom) neuron. Vertical red dashed lines indicate amplitude measurements. See Methods for additional details.

d) Example analysis of a single $Ca^{2+}$ transient showing the duration, area under the curve (AUC, grey shaded region) and threshold (crossings of the threshold define event initiation and termination) measurements.

e) Mean +/- SEM percentage of neurons in a field of view that participated in a network event for the indicated genotypes. Each dot represents a network event. n=467 WT, 708 KO, and 657 KO;Het network events from 20 individual culture wells per genotype. Culture wells were from 6-7 independent culture preps, 1 pup per prep. Kruskal-Wallis, $p<0.0001$; WT vs KO, ****$p<0.0001$; WT vs KO;Het, ****$p<0.0001$; KO vs KO;Het, ****$p<0.0001$; Dunn's multiple comparisons tests.

f) Mean +/- SEM frequency of network events per culture for the indicated genotypes. Each dot represents a single culture well. n=20 individual culture wells per genotype, from 6-7 independent culture preps, 1 pup per prep. One-way ANOVA, $p=0.0002$, $F (2, 57) = 9.773$; WT vs KO, ***$p=0.0003$; WT vs KO;Het, **$p=0.0049$; KO vs KO;Het, $p=0.7599$; Sidak's multiple comparisons tests. ns=non-significant.

g) Mean +/- SEM duration of network activity, expressed as the percent of the recording time during which network events occurred for the indicated genotypes. Each dot represents a single culture well. n=20 individual culture wells per genotype, from 6-7 independent culture preps, 1 pup per prep. Kruskal-Wallis, $p<0.0001$; WT vs KO, ****$p<0.0001$; WT vs KO;Het, ****$p<0.0001$; KO vs KO;Het, $p=0.5496$; Dunn's multiple comparisons tests.

h) Scatter dot plot of the average $Ca^{2+}$ transient amplitude per network event for the indicated genotypes. Black lines indicate the mean +/- SEM. n is the same as for panel e. Kruskal-Wallis, $p<0.0001$; WT vs KO, ****$p<0.0001$; WT vs KO;Het, $p>0.9999$; KO vs KO;Het, ****$p<0.0001$; Dunn's multiple comparison tests.

i) Scatter dot plot of the average $Ca^{2+}$ transient duration per network event for the indicated genotypes. Black lines indicate the mean +/- SEM. n is the same as for panel e. Kruskal-Wallis, $p<0.0001$; WT vs KO, ****$p<0.0001$; WT vs KO;Het, ****$p<0.0001$; KO vs KO;Het, $p>0.9999$; Dunn's multiple comparison tests.

j) Scatter dot plot the average $Ca^{2+}$ transient AUC per network event for the indicated genotypes. Black lines indicate the mean +/- SEM. n is the same as for panel e. Kruskal-Wallis, $p<0.0001$; WT vs KO, ****$p<0.0001$; WT vs KO;Het, ****$p<0.0001$; KO vs KO;Het, ****$p<0.0001$; Dunn's multiple comparison tests.

**Supplementary Figure 7**

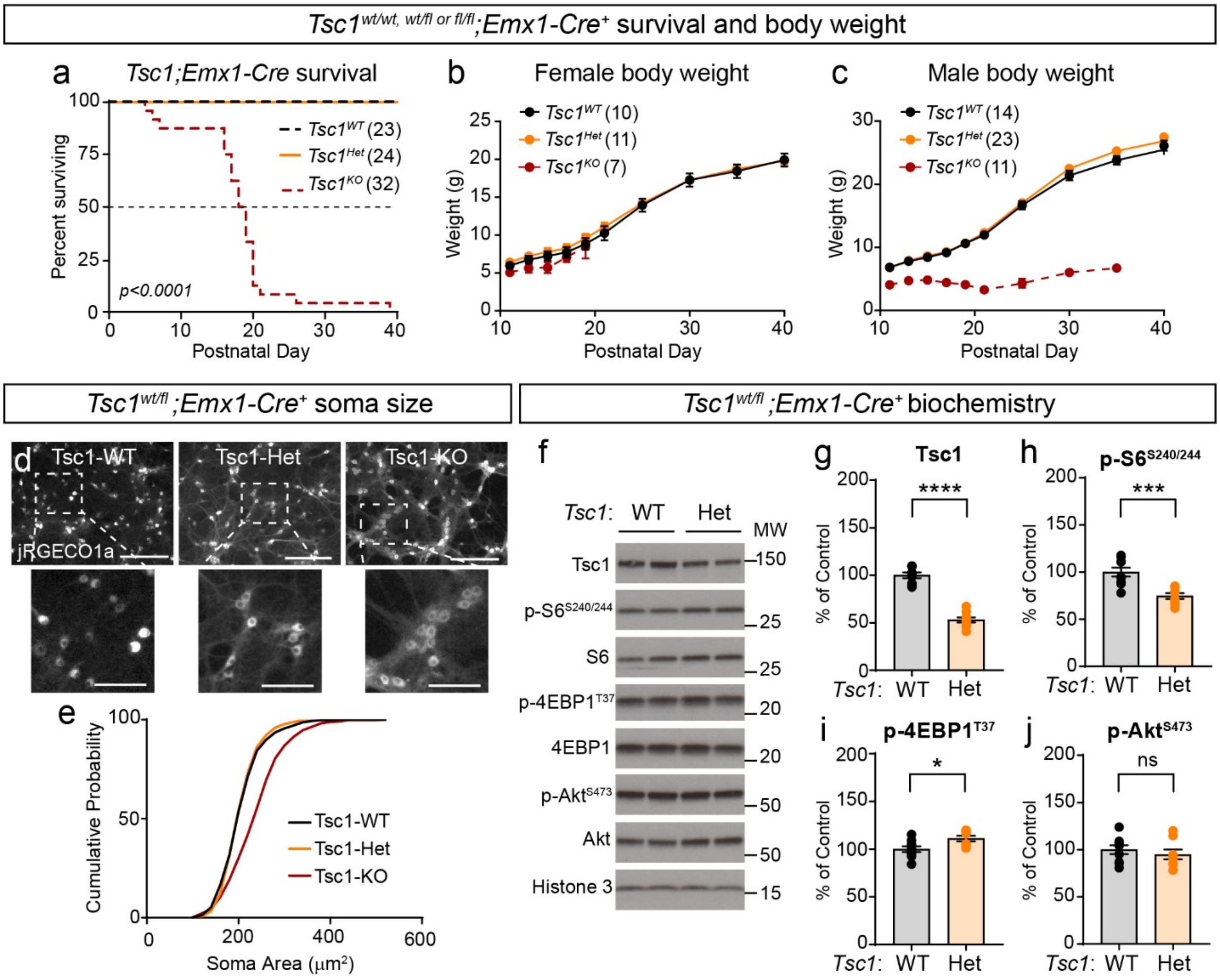

**Supplementary Figure 7. Heterozygous deletion of *Tsc1* does not affect survival, body weight, or soma size.**

**Related to Figures 1, 4 and 6.**

a) Survival analysis of *Tsc1*$^{wt/wt}$*;Rptor*$^{wt/wt}$*;Emx1*-Cre$^{+}$ (*Tsc1*$^{WT}$), *Tsc1*$^{wt/fl}$*;Rptor*$^{wt/wt}$*;Emx1*-Cre$^{+}$ (*Tsc1*$^{Het}$), and *Tsc1*$^{fl/fl}$*;Rptor*$^{wt/wt}$*;Emx1*-Cre$^{+}$ (*Tsc1*$^{KO}$) mice. Data for *Tsc1*$^{WT}$ and *Tsc1*$^{KO}$ mice are re-plotted from Fig. 4c for reference. The number of mice for each genotype is indicated in parentheses. Dashed line indicates 50% of the population surviving. P value from Log-rank Mantel-Cox tests is shown.

b,c) Mean +/- SEM body weight in grams measured from postnatal day 11 to 40 for female (b) and male (c) mice of the indicated genotypes. Data for *Tsc1*$^{WT}$ and *Tsc1*$^{KO}$ mice are re-plotted from Fig. 4e,f for reference. The number of mice for each genotype is indicated in parentheses. Mixed-effects model (REML) statistics: Females (b), day $p<0.0001$, $F(1.199, 25.84) = 1056$;

geno p=0.0569, F (2, 25) = 3.221. Males (c), day p<0.0001, F (1.792, 71.09) = 1113; geno p<0.0001, F (2, 44) = 29.07.

d) Representative images of *Tsc1$^{wt/wt}$;Emx1*-Cre$^{+}$ (WT), *Tsc1$^{wt/fl}$;Emx1*-Cre$^{+}$ (Tsc1-Het) and *Tsc1$^{fl/fl}$;Emx1*-Cre$^{+}$ (Tsc1-KO) primary hippocampal cultures expressing jRGeco1a on DIV 14. Scale bars for top panels=250 µm. Scale bars for in zoomed-in images (bottom panels)=100 µm.

e) Cumulative distributions of soma area for WT, Tsc1-Het and Tsc1-KO cultured hippocampal neurons. n=454 WT, 451 Tsc1-Het and 453 Tsc1-KO neurons from 5 independent culture preps, 1 pup per prep. Kruskal-Wallis test, p<0.0001; WT vs Het, p>0.9999; WT vs KO, ****p<0.0001; Het vs KO, ****p<0.0001; Dunn's multiple comparison tests.

f) Representative western blots of lysates collected from Tsc1-WT and Tsc1-Het primary hippocampal cultures. MW indicates molecular weight. Two independent samples per genotype are shown; this experiment was replicated three times.

g-j) Bar graphs display western blot quantification (mean +/- SEM) for Tsc1 (g) Welch's test, ****p<0.0001; p-S6 Ser240/244 (h) Welch's test, ***p= 0.0005; p-4EBP1 T37 (i) Mann-Whitney test, *p= 0.0152; and p-Akt Ser473 (j) Welch's test, p= 0.4734. Phospho-proteins were normalized to their respective total proteins. Dots represent data from individual culture wells. n=9 WT and 8 Tsc1-Het culture wells from 3 independent culture preps, 1 pup per prep.

### Supplementary Figure 8

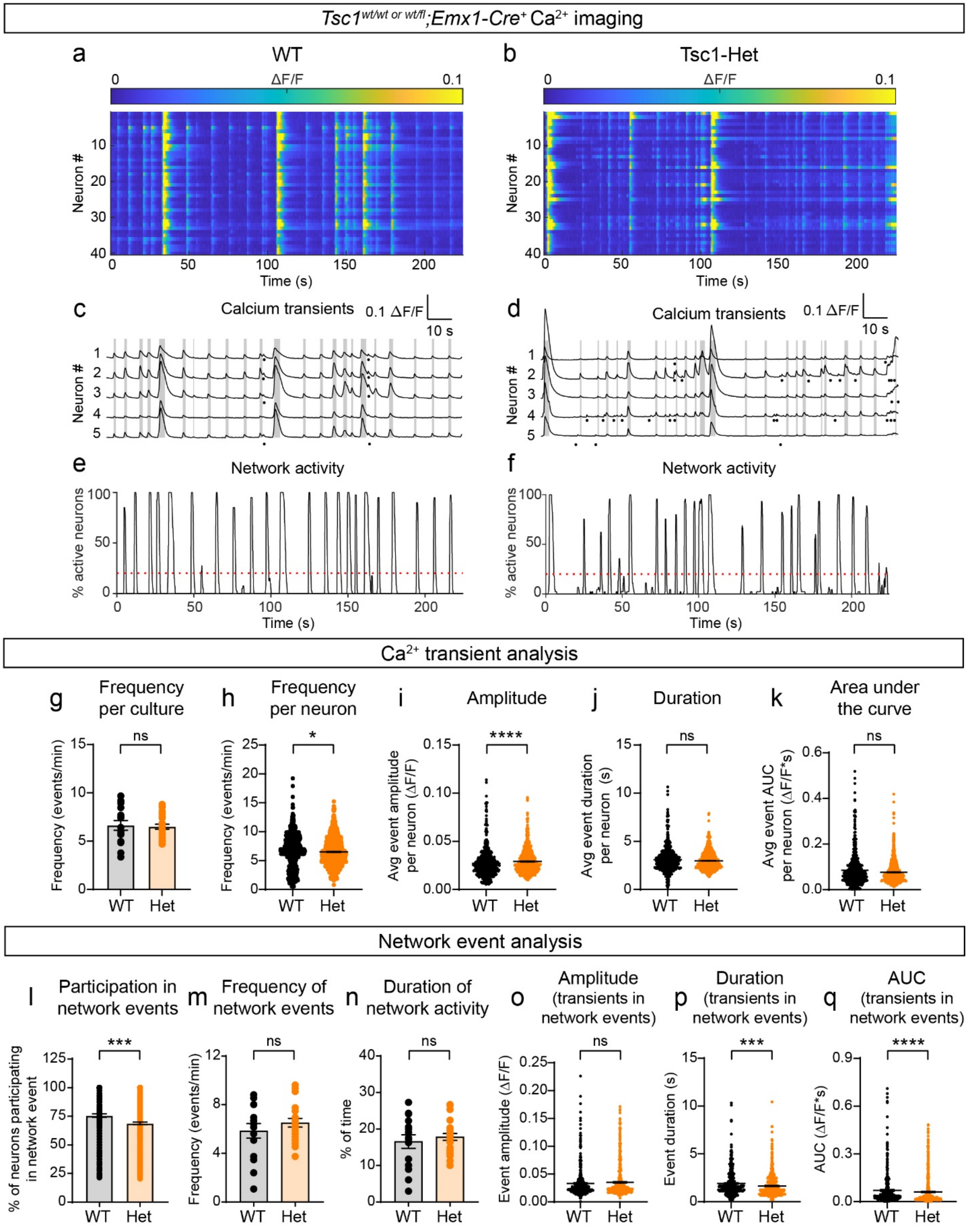

**Supplementary Figure 8. Loss of one copy of *Tsc1* does not induce neuronal or network hyperexcitability.**
**Related to Figure 6.**
a,b) Representative heatmaps of ∆F/F for 40 neurons imaged in a field of view from *Tsc1$^{wt/wt}$;Emx1*-Cre$^{+}$ (WT) and *Tsc1$^{wt/fl}$;Emx1*-Cre$^{+}$ (Tsc1-Het) cultures.
c,d) $Ca^{2+}$ transients from 5 representative neurons imaged in a field of view from WT (c) and Tsc1-Het (d) cultures. Grey lines indicate network events with more than 20% of neurons in the field of view active at the same time. Black dots represent spontaneous $Ca^{2+}$ transients that were not part of network events.
e,f) Graphs show the percentage of neurons in the field of view that were active at a given time for WT (d) and Tsc1-Het (e) cultures. One representative culture is shown per genotype. Red dashed lines at 20% indicate the threshold for a network event.
g) Mean +/- SEM $Ca^{2+}$ transient frequency per culture. Dots represent values from individual cultures. For WT: n=15 individual culture wells from 5 independent culture preps, 1 pup per prep for WT. For Tsc1-Het: n=24 individual culture wells from 9 independent culture preps, 1 pup per prep. Welch's test, p=0.8128; ns=non-significant.
h) Scatter dot plot of the $Ca^{2+}$ transient frequency per neuron for the indicated genotypes. Black lines indicate mean +/- SEM. For WT: n=600 neurons from 15 individual culture wells from 5 independent culture preps, 1 pup per prep. For Tsc1-Het: n=960 neurons from 24 individual culture wells from 9 independent culture preps, 1 pup per prep. Mann-Whitney test, *p = 0.0402.
i) Scatter dot plot of the average $Ca^{2+}$ transient amplitude per neuron for the indicated genotypes. n is the same as for panel h. Mann-Whitney test, ****p<0.0001.
j) Scatter dot plot of the average $Ca^{2+}$ transient duration per neuron for the indicated genotypes. n is the same as for panel h. Mann-Whitney test, p=0.0932.
k) Scatter dot plot of the average $Ca^{2+}$ transient area under the curve (AUC) per neuron for the indicated genotypes. n is the same as for panel h. Mann-Whitney test, p=0.4472.
l) Mean +/- SEM percentage of neurons in a field of view that participated in a network event for the indicated genotypes. Each dot represents a network event. For WT: n=362 network events from 15 individual culture wells from 5 independent culture preps, 1 pup per prep. For Tsc1-Het: n=585 network events from 24 individual culture wells from 9 independent culture preps, 1 pup per prep. Mann-Whitney test, ***p=0.0005.
m) Mean +/- SEM frequency of network events per culture for the indicated genotypes. Each dot represents a single culture well. For WT: n=15 individual culture wells from 5 independent

culture preps, 1 pup per prep. For Tsc1-Het: n=24 individual culture wells from 9 independent culture preps, 1 pup per prep. Welch's test, p=0.3462.

n) Mean +/- SEM duration of network activity, expressed as the percent of the recording time during which network events occurred for the indicated genotypes. Each dot represents a single culture well. n is the same as for panel m. Welch's test, p=0.5464

o) Scatter dot plot of the average $Ca^{2+}$ transient amplitude per network event for the indicated genotypes. Black lines indicate the mean +/- SEM. n is the same as for panel l. Mann-Whitney, p=0.8654.

p) Scatter dot plot of the average $Ca^{2+}$ transient duration per network event for the indicated genotypes. Black lines indicate the mean +/- SEM. n is the same as for panel l. Mann-Whitney, ***p=0.0003.

q) Scatter dot plot of the average $Ca^{2+}$ transient AUC per network event for the indicated genotypes. Black lines indicate the mean +/- SEM. n is the same as for panel l. Mann-Whitney, ****p<0.0001.

**Supplementary Figure 9**

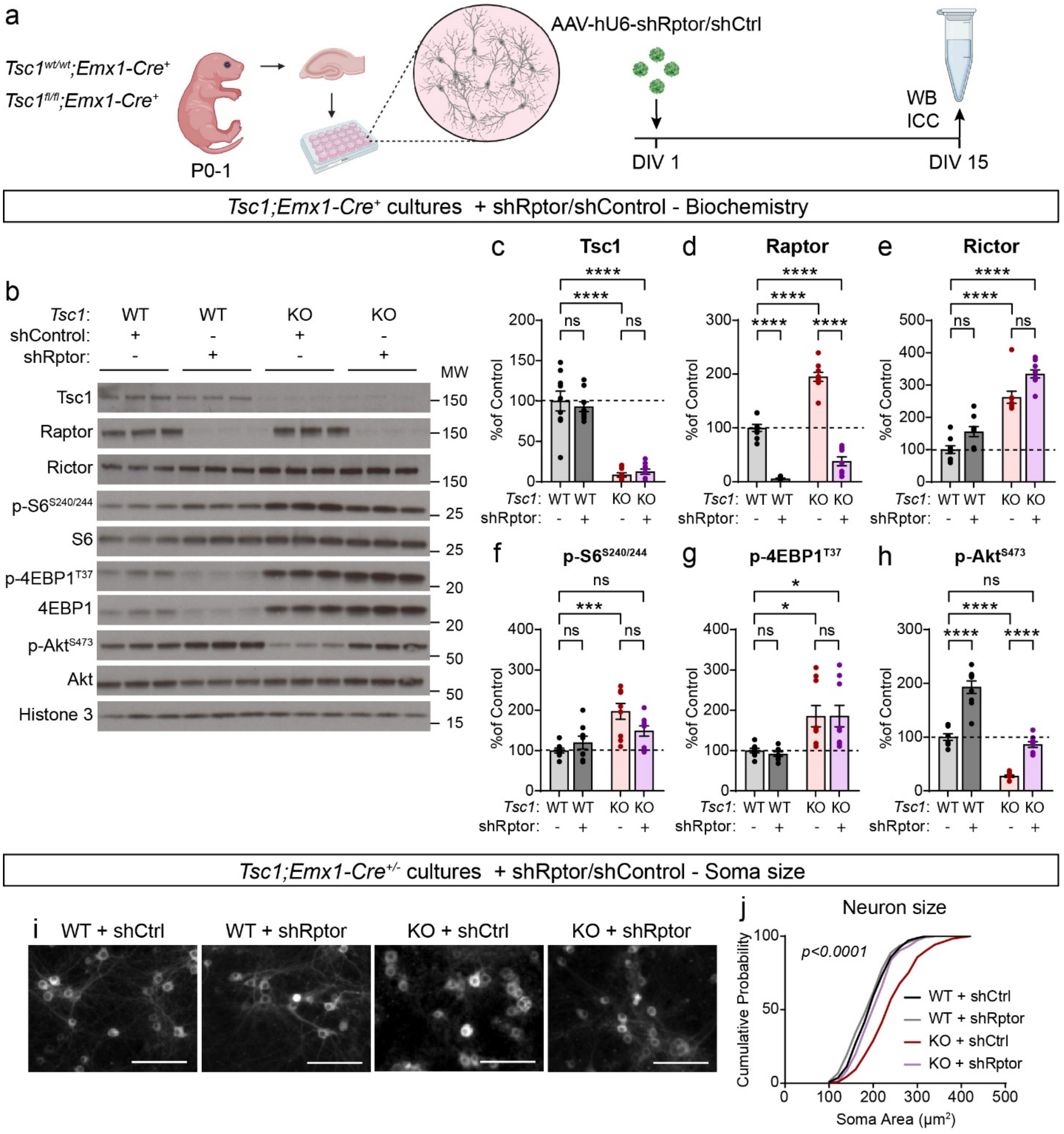

**Supplementary Figure 9. shRptor rebalances mTOR signaling and rescues neuronal hypertrophy in cultured Tsc1-KO neurons.**

**Related to Figure 7.**

a) Schematic of the experiment. Primary hippocampal cultures were prepared from P0-1 *Tsc1*$^{wt/wt}$*;Emx1-Cre*$^{+}$ (WT) and *Tsc1*$^{fl/fl}$*;Emx1-Cre*$^{+}$ (Tsc1-KO) mice and transduced with AAVshControl-EYFP (shControl) or AAV-shRptor-EYFP (shRptor) on DIV 1. Cell were collected for analysis by western blot (WB) or immunocytochemistry (ICC) on DIV 15. Created with BioRender.com

b) Representative western blots of lysates collected from WT and Tsc1-KO primary hippocampal cultures treated with shControl or shRptor. MW indicates molecular weight. Three samples per genotype and treatment are shown; this experiment was replicated three times.

c-h) Bar graphs display western blot quantification (mean +/- SEM) for the indicated proteins, expressed as a percentage of control (WT + shControl) levels. Phospho-proteins were normalized to their respective total proteins. Dots represent data from individual culture wells. n=8-9 culture wells per condition from 3 independent culture preps, 1 pup per prep.

c) Tsc1, One-way ANOVA, $p<0.0001$, $F(3, 32) = 48.58$; WT+shControl vs WT+shRptor, p= 0.9360; WT+shControl vs KO+shControl, ****$p<0.0001$; WT+shControl vs KO+shRptor, ****$p<0.0001$; KO+shControl vs KO+shRptor, p=0.9918; Sidak's multiple comparison tests. ns=non-significant.

d) Raptor, One-way ANOVA, $p<0.0001$, $F(3, 32) = 153.6$; WT+shControl vs WT+shRptor, ****$p<0.0001$; WT+shControl vs KO+shControl, ****$p<0.0001$; WT+shControl vs KO+shRptor, ****$p<0.0001$; KO+shControl vs KO+shRptor, ****$p<0.0001$; Sidak's multiple comparison tests.

e) Rictor, Kruskal-Wallis, $p<0.0001$; WT+shControl vs WT+shRptor, p=0.8411; WT+shControl vs KO+shControl, **p=0.0023; WT+shControl vs KO+shRptor, ****$p<0.0001$; KO+shControl vs KO+shRptor, p=0.5592; Dunn's multiple comparison tests.

f) p-S6 Ser240/244, One-way ANOVA, p=0.0004, $F(3, 30) = 8.234$; WT+shControl vs WT+shRptor, p=0.8388; WT+shControl vs KO+shControl, ***p=0.0003; WT+shControl vs KO+shRptor, p=0.1045; KO+shControl vs KO+shRptor, p=0.0894; Sidak's multiple comparison tests.

g) p-4EBP1 T37, One-way ANOVA, p=0.0015, $F(3, 30) = 6.582$; WT+shControl vs WT+shRptor, p=0.9973; WT+shControl vs KO+shControl, *p=0.0224; WT+shControl vs KO+shRptor,* p=0.0220; KO+shControl vs KO+shRptor, $p>0.9999$; Sidak's multiple comparison tests.

h) p-Akt Ser473, One-way ANOVA, $p<0.0001$, $F(3, 32) = 92.30$; WT+shControl vs WT+shRptor, ****$p<0.0001$; WT+shControl vs KO+shControl, ****$p<0.0001$; WT+shControl vs KO+shRptor, p=0.5561; KO+shControl vs KO+shRptor, ****$p<0.0001$; Sidak's multiple comparison tests.

i) Representative images of DIV 15 WT and Tsc1-KO primary hippocampal cultures transduced with shControl or shRptor expressing jRGeco1a. Scale bars=100 μm.

j) Cumulative distributions of soma area for WT and Tsc1-KO cultured hippocampal neurons treated with shControl or shRptor. n=246 WT+shControl, 249 WT+shRptor, 245 Tsc1-KO+shControl and 246 Tsc1-KO+shRptor neurons from 8 culture wells from 4 independent culture preps, 1 pup per prep. Kruskal-Wallis, $p<0.0001$; WT+shControl vs WT+shRptor, $p=0.4789$; WT+shControl vs KO+shControl, $p<0.0001$; WT+shControl vs KO+shRptor, $p=0.2352$; KO+shControl vs KO+shRptor, $p<0.0001$; Dunn's multiple comparison tests.

**Supplementary Figure 10**

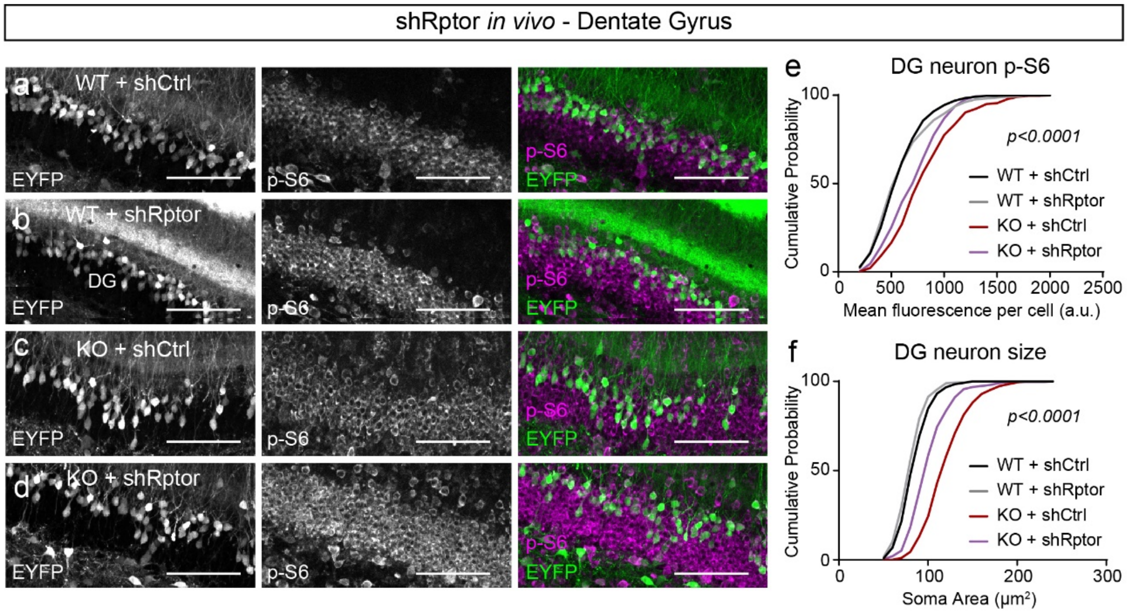

**Supplementary Figure 10. shRptor improves cellular phenotypes in the dentate gyrus of Tsc1-cKO mice.**

**Related to Figure 7.**

a-d) Representative images of the dentate gyrus (DG) of *Tsc1$^{wt/wt}$;Emx1-Cre$^{+}$* (WT, a-b) and *Tsc1$^{fl/fl}$;Emx1-Cre$^{+}$* (KO, c-d) mice injected with AAV-shRptor-EYFP or AAV-shCtrl-EYFP virus showing EYFP fluorescence (left panels) and p-S6 240/244 immunostaining (middle panels). Right panels show merged images. Scale bars=100 μm.

e) Cumulative distributions of p-S6 levels in DG EYFP+ neurons for the indicated genotypes. n=396 WT+shCtrl, 396 WT+shRptor, 382 Tsc1-KO+shCtrl and 350 Tsc1-KO+shRptor neurons from 6 mice per group. Kruskal-Wallis, $p<0.0001$; WT+shCtrl vs WT+shRptor, $p>0.9999$; WT+shCtrl vs Tsc1-KO+shCtrl, $p<0.0001$; WT+shCtrl vs Tsc1-KO+shRptor, $p<0.0001$; Tsc1-KO+shCtrl vs Tsc1-KO+shRptor, $p=0.0026$; Dunn's multiple comparison tests.

f) Cumulative distributions of EYFP+ DG neuron soma area for the indicated genotypes. n is the same as in panel e. Kruskal-Wallis, $p<0.0001$; WT+shCtrl vs WT+shRptor, $p=0.0241$; WT+shCtrl vs Tsc1-KO+shCtrl, $p<0.0001$; WT+shCtrl vs Tsc1-KO+shRptor, $p<0.0001$; Tsc1-KO+shCtrl vs Tsc1-KO+shRptor, $p<0.0001$; Dunn's multiple comparison tests.

**Supplementary Table 1. Survival analysis of *Tsc1;Rptor;Emx1-Cre* and *Tsc1;Rictor;Emx1-Cre* mice. Related to Figure 4.**

| Mouse line | Genotype | Sex | Day of first mortality (P0-P40)[1] | Median survival in days (P0-P40)[1] | Age of oldest surviving animal (days)[2] |
|---|---|---|---|---|---|
| *Tsc1;Rptor;Emx1-Cre* | *Tsc1*$^{wt/wt}$*;Rptor*$^{wt/wt}$*;Emx1-Cre*$^{+}$ | M | n/a | n/a | >150 |
| *Tsc1;Rptor;Emx1-Cre* | *Tsc1*$^{wt/wt}$*;Rptor*$^{wt/wt}$*;Emx1-Cre*$^{wt/+}$ | F | n/a | n/a | >150 |
| *Tsc1;Rptor;Emx1-Cre* | *Tsc1*$^{wt/wt}$*;Rptor*$^{wt/fl}$*;Emx1-Cre*$^{+}$ | M | n/a | n/a | >150 |
| *Tsc1;Rptor;Emx1-Cre* | *Tsc1*$^{wt/wt}$*; Rptor*$^{wt/fl}$*;Emx1-Cre*$^{+}$ | F | n/a | n/a | >150 |
| *Tsc1;Rptor;Emx1-Cre* | *Tsc1*$^{wt/wt}$*;Rptor*$^{fl/fl}$*;Emx1-Cre*$^{wt/+}$ | M | n/a | n/a | >150 |
| *Tsc1;Rptor;Emx1-Cre* | *Tsc1*$^{wt/wt}$*;Rptor*$^{fl/fl}$*;Emx1-Cre*$^{+}$ | F | n/a | n/a | >150 |
| *Tsc1;Rptor;Emx1-Cre* | *Tsc1*$^{wt/fl}$*;Rptor*$^{wt/wt}$*;Emx1-Cre*$^{+}$ | M | n/a | n/a | >150 |
| *Tsc1;Rptor;Emx1-Cre* | *Tsc1*$^{wt/fl}$*;Rptor*$^{wt/wt}$*;Emx1-Cre*$^{+}$ | F | n/a | n/a | >150 |
| *Tsc1;Rptor;Emx1-Cre* | *Tsc1*$^{wt/fl}$*;Rptor*$^{wt/fl}$*;Emx1-Cre*$^{+}$ | M | n/a | n/a | >150 |
| *Tsc1;Rptor;Emx1-Cre* | *Tsc1*$^{wt/fl}$*;Rptor*$^{wt/fl}$*; Emx1-Cre*$^{+}$ | F | n/a | n/a | >150 |
| *Tsc1;Rptor;Emx1-Cre* | *Tsc1*$^{wt/fl}$*;Rptor*$^{fl/fl}$*;Emx1-Cre*$^{+}$ | M | n/a | n/a | >150 |
| *Tsc1;Rptor;Emx1-Cre* | *Tsc1*$^{wt/fl}$*;Rptor*$^{fl/fl}$*;Emx1-Cre*$^{+}$ | F | n/a | n/a | >150 |
| *Tsc1;Rptor;Emx1-Cre* | *Tsc1*$^{fl/fl}$*;Rptor*$^{wt/wt}$*;Emx1-Cre*$^{+}$ | M[3] | 16 | 19 | 39 |
| *Tsc1;Rptor;Emx1-Cre* | *Tsc1*$^{fl/fl}$*;Rptor*$^{wt/wt}$*;Emx1-Cre*$^{+}$ | F[3] | 16 | 18.5 | 20 |
| *Tsc1;Rptor;Emx1-Cre* | *Tsc1*$^{fl/fl}$*;Rptor*$^{wt/fl}$*;Emx1-Cre*$^{+}$ | M | 18 | 24 | >150 |
| *Tsc1;Rptor;Emx1-Cre* | *Tsc1*$^{fl/fl}$*;Rptor*$^{wt/fl}$*;Emx1-Cre*$^{+}$ | F | 17 | 25 | >150 |
| *Tsc1;Rptor;Emx1-Cre* | *Tsc1*$^{fl/fl}$*;Rptor*$^{fl/fl}$*;Emx1-Cre*$^{+}$ | M | n/a | n/a | >150 |
| *Tsc1;Rptor;Emx1-Cre* | *Tsc1*$^{fl/fl}$*;Rptor*$^{fl/fl}$*;Emx1-Cre*$^{+}$ | F | 30 | n/a[4] | >150 |

| | | | | | |
|---|---|---|---|---|---|
| *Tsc1;Rictor;Emx1-Cre* | $Tsc1^{wt/wt};Rictor^{wt/wt};Emx1\text{-}Cre^{+}$ | M | n/a | n/a | - |
| *Tsc1;Rictor;Emx1-Cre* | $Tsc1^{wt/wt};Rictor^{wt/wt};Emx1\text{-}Cre^{+}$ | F | n/a | n/a | - |
| *Tsc1;Rictor;Emx1-Cre* | $Tsc1^{wt/wt};Rictor^{wt/fl};Emx1\text{-}Cre^{+}$ | M | n/a | n/a | - |
| *Tsc1;Rictor;Emx1-Cre* | $Tsc1^{wt/wt};Rictor^{wt/fl};Emx1\text{-}Cre^{+}$ | F | n/a | n/a | - |
| *Tsc1;Rictor;Emx1-Cre* | $Tsc1^{wt/wt};Rictor^{fl/fl};Emx1\text{-}Cre^{+}$ | M | n/a | n/a | - |
| *Tsc1;Rictor;Emx1-Cre* | $Tsc1^{wt/wt};Rictor^{fl/fl};Emx1\text{-}Cre^{+}$ | F | n/a | n/a | - |
| *Tsc1;Rictor;Emx1-Cre* | $Tsc1^{wt/fl};Rictor^{wt/wt};Emx1\text{-}Cre^{+}$ | M | n/a | n/a | - |
| *Tsc1;Rictor;Emx1-Cre* | $Tsc1^{wt/fl};Rictor^{wt/wt};Emx1\text{-}Cre^{+}$ | F | n/a | n/a | - |
| *Tsc1;Rictor;Emx1-Cre* | $Tsc1^{wt/fl};Rictor^{wt/fl};Emx1\text{-}Cre^{+}$ | M | n/a | n/a | - |
| *Tsc1;Rictor;Emx1-Cre* | $Tsc1^{wt/fl};Rictor^{wt/fl};Emx1\text{-}Cre^{+}$ | F | n/a | n/a | - |
| *Tsc1;Rictor;Emx1-Cre* | $Tsc1^{wt/fl};Rictor^{fl/fl};Emx1\text{-}Cre^{+}$ | M | n/a | n/a | - |
| *Tsc1;Rictor;Emx1-Cre* | $Tsc1^{wt/fl};Rictor^{fl/fl};Emx1\text{-}Cre^{+}$ | F | n/a | n/a | - |
| *Tsc1;Rictor;Emx1-Cre* | $Tsc1^{fl/fl};Rictor^{wt/wt};Emx1\text{-}Cre^{+}$ | M | 15 | 19 | 19 |
| *Tsc1;Rictor;Emx1-Cre* | $Tsc1^{fl/fl};Rictor^{wt/wt};Emx1\text{-}Cre^{+}$ | F | 17 | 18 | 18 |
| *Tsc1;Rictor;Emx1-Cre* | $Tsc1^{fl/fl};Rictor^{wt/fl};Emx1\text{-}Cre^{+}$ | M | 14 | 18 | 21 |
| *Tsc1;Rictor;Emx1-Cre* | $Tsc1^{fl/fl};Rictor^{wt/fl};Emx1\text{-}Cre^{+}$ | F | 14 | 17 | 21 |
| *Tsc1;Rictor;Emx1-Cre* | $Tsc1^{fl/fl};Rictor^{fl/fl};Emx1\text{-}Cre^{+}$ | M | 18 | 19.5 | 22 |
| *Tsc1;Rictor;Emx1-Cre* | $Tsc1^{fl/fl};Rictor^{fl/fl};Emx1\text{-}Cre^{+}$ | F | 18 | 20 | 24 |

Footnotes:

[1] n/a = No animals died within the first 40 postnatal days

[2] - = Animals were not monitored past the first 40 postnatal days

[3] Three $Tsc1^{fl/fl};Rptor^{wt/wt};Emx1\text{-}Cre^{+}$ mice were found dead before P10 and sex could not be determined

[4] One $Tsc1^{fl/fl};Rptor^{fl/fl};Emx1\text{-}Cre^{+}$ female died at P30

**Supplementary Table 2. Body weights of *Tsc1;Rptor;Emx1-Cre* and *Tsc1;Rictor;Emx1-Cre* mice. Related to Figure 4.**

| Mouse line | Genotype | Sex | Mean +/- SEM body weight at P15 (n) | Mean +/- SEM body weight at P150 (n)[1,2] |
|---|---|---|---|---|
| *Tsc1;Rptor;Emx1-Cre* | *Tsc1*$^{wt/wt}$;*Rptor*$^{wt/wt}$;*Emx1-Cre*$^{+}$ | M | 8.42 +/- 0.31 (14) | 48.00 +/- 5.69 (3) |
| *Tsc1;Rptor;Emx1-Cre* | *Tsc1*$^{wt/wt}$;*Rptor*$^{wt/wt}$;*Emx1-Cre*$^{+}$ | F | 7.56 +/- 0.57 (9) | 38.17 +/- 1.92 (3) |
| *Tsc1;Rptor;Emx1-Cre* | *Tsc1*$^{wt/wt}$;*Rptor*$^{wt/fl}$;*Emx1-Cre*$^{+}$ | M | 8.38 +/- 0.32 (14) | 45.06 +/- 1.99 (9) |
| *Tsc1;Rptor;Emx1-Cre* | *Tsc1*$^{wt/wt}$;*Rptor*$^{wt/fl}$;*Emx1-Cre*$^{+}$ | F | 7.90 +/- 0.40 (13) | 38.73 +/- 4.52 (6) |
| *Tsc1;Rptor;Emx1-Cre* | *Tsc1*$^{wt/wt}$;*Rptor*$^{fl/fl}$;*Emx1-Cre*$^{+}$ | M | 6.26 +/- 0.36 (7) | 22.50 +/- 0.40 (2) |
| *Tsc1;Rptor;Emx1-Cre* | *Tsc1*$^{wt/wt}$;*Rptor*$^{fl/fl}$;*Emx1-Cre*$^{+}$ | F | 6.28 +/- 0.48 (10) | 21.18 +/- 1.12 (5) |
| *Tsc1;Rptor;Emx1-Cre* | *Tsc1*$^{wt/fl}$;*Rptor*$^{wt/wt}$;*Emx1-Cre*$^{+}$ | M | 8.64 +/- 0.30 (23) | 46.60 +/- 1.50 (6) |
| *Tsc1;Rptor;Emx1-Cre* | *Tsc1*$^{wt/fl}$;*Rptor*$^{wt/wt}$;*Emx1-Cre*$^{+}$ | F | 7.78 +/- 0.51 (11) | 44.20 +/- 1.10 (2) |
| *Tsc1;Rptor;Emx1-Cre* | *Tsc1*$^{wt/fl}$;*Rptor*$^{wt/fl}$;*Emx1-Cre*$^{+}$ | M | 8.03 +/- 0.29 (38) | 43.05 +/- 1.52 (16) |
| *Tsc1;Rptor;Emx1-Cre* | *Tsc1*$^{wt/fl}$;*Rptor*$^{wt/fl}$;*Emx1-Cre*$^{+}$ | F | 8.44 +/- 0.23 (39) | 36.47 +/- 3.01 (11) |
| *Tsc1;Rptor;Emx1-Cre* | *Tsc1*$^{wt/fl}$;*Rptor*$^{fl/fl}$;*Emx1-Cre*$^{+}$ | M | 6.72 +/- 0.42 (17) | 24.50 +/- 1.17 (3) |
| *Tsc1;Rptor;Emx1-Cre* | *Tsc1*$^{wt/fl}$;*Rptor*$^{fl/fl}$;*Emx1-Cre*$^{+}$ | F | 5.64 +/- 0.88 (9) | 20.05 +/- 1.45 (2) |
| *Tsc1;Rptor;Emx1-Cre* | *Tsc1*$^{fl/fl}$;*Rptor*$^{wt/wt}$;*Emx1-Cre*$^{+}$ | M | 4.81 +/- 0.34 (11) | n/a |
| *Tsc1;Rptor;Emx1-Cre* | *Tsc1*$^{fl/fl}$;*Rptor*$^{wt/wt}$;*Emx1-Cre*$^{+}$ | F | 5.55 +/- 0.63 (8) | n/a |
| *Tsc1;Rptor;Emx1-Cre* | *Tsc1*$^{fl/fl}$;*Rptor*$^{wt/fl}$;*Emx1-Cre*$^{+}$ | M | 7.00 +/- 0.50 (16) | 29.60 (1) |
| *Tsc1;Rptor;Emx1-Cre* | *Tsc1*$^{fl/fl}$;*Rptor*$^{wt/fl}$;*Emx1-Cre*$^{+}$ | F | 6.65 +/- 0.29 (23) | 17.60 (1) |
| *Tsc1;Rptor;Emx1-Cre* | *Tsc1*$^{fl/fl}$;*Rptor*$^{fl/fl}$;*Emx1-Cre*$^{+}$ | M | 5.68 +/- 0.55 (8) | 21.15 +/- 1.45 (2) |
| *Tsc1;Rptor;Emx1-Cre* | *Tsc1*$^{fl/fl}$;*Rptor*$^{fl/fl}$;*Emx1-Cre*$^{+}$ | F | 6.80 +/- 0.32 (9) | 19.29 +/- 0.51 (7) |
| *Tsc1;Rictor;Emx1-Cre* | *Tsc1*$^{wt/wt}$;*Rictor*$^{wt/wt}$;*Emx1-Cre*$^{+}$ | M | 9.42 +/- 0.24 (12) | - |

| | | | | |
|---|---|---|---|---|
| *Tsc1;Rictor;Emx1-Cre* | *Tsc1*$^{wt/wt}$;*Rictor*$^{wt/wt}$;*Emx1-Cre*$^{+}$ | F | 8.78 +/- 0.28 (12) | - |
| *Tsc1;Rictor;Emx1-Cre* | *Tsc1*$^{wt/wt}$;*Rictor*$^{wt/fl}$;*Emx1-Cre*$^{+}$ | M | 9.63 +/- 0.22 (6) | - |
| *Tsc1;Rictor;Emx1-Cre* | *Tsc1*$^{wt/wt}$;*Rictor*$^{wt/fl}$;*Emx1-Cre*$^{+}$ | F | 9.45 +/- 0.81 (6) | - |
| *Tsc1;Rictor;Emx1-Cre* | *Tsc1*$^{wt/wt}$;*Rictor*$^{fl/fl}$;*Emx1-Cre*$^{+}$ | M | 7.55 +/- 0.45 (6) | - |
| *Tsc1;Rictor;Emx1-Cre* | *Tsc1*$^{wt/wt}$;*Rictor*$^{fl/fl}$;*Emx1-Cre*$^{+}$ | F | 9.07 +/- 0.69 (3) | - |
| *Tsc1;Rictor;Emx1-Cre* | *Tsc1*$^{wt/fl}$;*Rictor*$^{wt/wt}$;*Emx1-Cre*$^{+}$ | M | 9.38 +/- 0.42 (19) | - |
| *Tsc1;Rictor;Emx1-Cre* | *Tsc1*$^{wt/fl}$;*Rictor*$^{wt/wt}$;*Emx1-Cre*$^{+}$ | F | 8.76 +/- 0.47 (14) | - |
| *Tsc1;Rictor;Emx1-Cre* | *Tsc1*$^{wt/fl}$;*Rictor*$^{wt/fl}$;*Emx1-Cre*$^{+}$ | M | 9.54 +/- 0.27 (27) | - |
| *Tsc1;Rictor;Emx1-Cre* | *Tsc1*$^{wt/fl}$;*Rictor*$^{wt/fl}$;*Emx1-Cre*$^{+}$ | F | 9.48 +/- 0.35 (30) | - |
| *Tsc1;Rictor;Emx1-Cre* | *Tsc1*$^{wt/fl}$;*Rictor*$^{fl/fl}$;*Emx1-Cre*$^{+}$ | M | 8.56 +/- 0.33 (24) | - |
| *Tsc1;Rictor;Emx1-Cre* | *Tsc1*$^{wt/fl}$;*Rictor*$^{fl/fl}$;*Emx1-Cre*$^{+}$ | F | 8.07 +/-0.32 (16) | - |
| *Tsc1;Rictor;Emx1-Cre* | *Tsc1*$^{fl/fl}$;*Rictor*$^{wt/wt}$;*Emx1-Cre*$^{+}$ | M | 5.31 +/- 0.54 (9) | n/a |
| *Tsc1;Rictor;Emx1-Cre* | *Tsc1*$^{fl/fl}$;*Rictor*$^{wt/wt}$;*Emx1-Cre*$^{+}$ | F | 5.60 +/- 0.39 (4) | n/a |
| *Tsc1;Rictor;Emx1-Cre* | *Tsc1*$^{fl/fl}$;*Rictor*$^{wt/fl}$;*Emx1-Cre*$^{+}$ | M | 5.82 +/- 0.59 (14) | n/a |
| *Tsc1;Rictor;Emx1-Cre* | *Tsc1*$^{fl/fl}$;*Rictor*$^{wt/fl}$;*Emx1-Cre*$^{+}$ | F | 4.81 +/- 0.38 (13) | n/a |
| *Tsc1;Rictor;Emx1-Cre* | *Tsc1*$^{fl/fl}$;*Rictor*$^{fl/fl}$;*Emx1-Cre*$^{+}$ | M | 5.91 +/- 0.34 (10) | n/a |
| *Tsc1;Rictor;Emx1-Cre* | *Tsc1*$^{fl/fl}$;*Rictor*$^{fl/fl}$;*Emx1-Cre*$^{+}$ | F | 4.76 +/- 0.34 (5) | n/a |

Footnotes:

[1] n/a = No animals survived beyond postnatal day 40

[2] - = Animals were not monitored past the first 40 postnatal days

**Supplementary Table 3. Summary of *in vivo* phenotypes in *Tsc1;Rptor;Emx1-Cre* mice by genotype and sex. Related to Figures 4 and 5.**

| Genotype | *Tsc1$^{wt/wt}$;Rptor$^{wt/wt}$; Emx1-Cre$^{+}$* | | *Tsc1$^{fl/fl}$;Rptor$^{wt/wt}$; Emx1-Cre$^{+}$* | | *Tsc1$^{fl/fl}$;Rptor$^{wt/fl}$; Emx1-Cre$^{+}$* | |
|---|---|---|---|---|---|---|
| **Sex** | **Females (n=4)** | **Males (n=4)** | **Females (n=4)** | **Males (n=4)** | **Females (n=4)** | **Males (n=4)** |
| | **Mean +/- SEM** | | **Mean +/- SEM** | | **Mean +/- SEM** | |
| Cortical thickness (μm) | 963.50 +/- 37.62 | 990.10 +/- 54.72 | 1176.00 +/- 48.51 | 1262.00 +/- 24.62 | 1085.00 +/- 14.12 | 1105.00 +/- 46.99 |
| CA1 thickness (μm) | 56.85 +/- 3.16 | 56.03 +/- 4.83 | 83.58 +/- 7.54 | 95.01 +/- 5.33 | 80.26 +/- 13.22 | 69.82 +/- 7.62 |
| DG suprapyramidal blade thickness (μm) | 48.20 +/- 2.97 | 52.09 +/- 4.23 | 66.25 +/- 3.58 | 66.21 +/- 3.57 | 52.68 +/- 6.39 | 52.29 +/- 7.81 |
| DG infrapyramydal blade thickness (μm) | 38.52 +/- 2.19 | 44.01 +/- 4.47 | 55.92 +/- 3.88 | 66.53 +/- 1.22 | 51.90 +/- 2.33 | 43.33 +/- 4.45 |
| GFAP intensity across cortical layers (a.u.) | 213.30 +/- 3.07 | 190.80 +/- 2.72 | 288.60 +/- 6.09 | 292.10 +/- 5.37 | 300.70 +/- 5.81 | 236.30 +/- 5.77 |
| GFAP intensity in CA1(a.u.) | 278.90 +/- 53.94 | 263.10 +/- 23.43 | 363.40 +/- 32.79 | 457.00 +/- 51.27 | 339.40 +/- 14.30 | 312.20 +/- 28.31 |
| MBP intensity (a.u.) | 13340.00 +/- 670.00 | 13365.00 +/- 988.50 | 7561.00 +/- 418.40 | 7143.00 +/- 319.60 | 10059.00 +/- 933.40 | 9946.00 +/- 790.20 |
| # of ectopic neurons above CA1 | 4.25 +/- 0.25 | 3.75 +/- 0.85 | 33.00 +/- 6.49 | 45.50 +/- 4.83 | 23.00 +/- 9.00 | 26.00 +/- 1.47 |
| Cortical neurons soma area (μm) | 362.30 +/- 4.18 | 361.50 +/- 3.90 | 403.80 +/- 4.14 | 420.90 +/- 3.93 | 383.00 +/- 3.93 | 380.50 +/- 3.72 |
| Cortical neurons p-S6 intensity (a.u.) | 107.30 +/- 1.19 | 92.71 +/- 0.99 | 131.40 +/- 1.53 | 119.80 +/- 1.27 | 126.80+/- 1.57 | 97.96 +/- 1.37 |
| CA1 neurons soma area (μm) | 272.70 +/- 4.16 | 277.70 +/- 4.55 | 409.50 +/- 8.24 | 409.30 +/- 8.32 | 285.60 +/- 3.99 | 286.30 +/- 3.45 |
| CA1 neurons p-S6 intensity (a.u.) | 98.38 +/- 1.47 | 101.60 +/- 1.30 | 149.90 +/- 2.27 | 124.20 +/- 2.32 | 118.50 +/- 1.86 | 85.92 +/- 1.53 |
| DG neurons soma area (μm) | 209.00 +/- 3.27 | 217.60 +/- 3.34 | 233.30 +/- 3.56 | 259.80 +/- 4.57 | 232.50 +/- 3.24 | 216.80 +/- 3.39 |
| DG neurons p-S6 intensity (a.u.) | 96.49 +/- 1.72 | 103.50 +/- 1.61 | 162.20 +/- 4.29 | 132.00 +/- 4.17 | 126.40 +/- 2.27 | 83.65 +/- 1.67 |

**Supplementary Table 4. Mouse strains and genotyping primers.**

| Mouse Line | Genotyping Primers | Source | Reference |
|---|---|---|---|
| *Emx1-Cre* | WT F: AAG GTG TGG TTC CAG AAT CG | JAX strain # 005628 | 49 |
| | WT R: CTC TCC ACC AGA AGG CTG AG | | |
| | Mut F: GCG GTC TGG CAG TAA AAA CTA TC | | |
| | Mut R: GTG AAA CAG CAT TGC TGT CAC TT | | |
| $Tsc1^{fl/fl}$ | F: GTC ACG ACC GTA GGA GAA GC | JAX strain # 005680 | 44 |
| | R: GAA TCA ACC CCA CAG AGC AT | | |
| $Rptor^{fl/fl}$ | F: AGCCTTTAGTACCCACTTGGC | JAX strain # 013188 | 45 |
| | R: GGCATCTCACAAAGGGTACAG | | |
| $Rictor^{fl/fl}$ | F: ACTGATATGTTCATGGTTGTG | JAX strain # 020649 | 46,47 |
| | R: GACACTGGATTCAGTGGCTTG | | |
| Ai9 | WT F: AAG GGA GCT GCA GTG GAG TA | JAX strain # 007909 | 96 |
| | WT R: CCG AAA ATC TGT GGG AAG TC | | |
| | Mut F: CTG TTC CTG TAC GGC ATG G | | |
| | Mut R: GGC ATT AAA GCA GCG TAT CC | | |

**Supplementary Table 5. Viruses and titers.**

| Virus | Serotype | Promoter | Source | Titer of viral stock (vg/ml) | Dilution amount (*in vitro*) | Dilution amount (*in vivo*) |
|---|---|---|---|---|---|---|
| AAV1.hSyn.HI.eGFP-Cre.WPRE.SV40 | 1 | hSyn | Penn Vector Core | $1.78 \times 10^{13}$ | 1:20 0.5 µl/well | n/a |
| AAV1.hSyn.eGFP.WPRE.bGH | 1 | hSyn | Penn Vector Core | $3.86 \times 10^{13}$ | 1:20 0.5 µl/well | n/a |
| AAV5.hSyn.eGFP | 5 | hSyn | UNC Vector Core | $4 \times 10^{12}$ | 1:100 0.5 | n/a |
| AAV1.CBA.mCherry-nCre.WPRE.bGH | 1 | CBA | Penn Vector Core | $1.04 \times 10^{13}$ | 1:100 0.5 µl/well | n/a |
| AAV9.CAG.Flex.tdTomato.WPRE.bGH (AllenInstitute864) | 9 | CAG | Penn Vector Core | unknown | 1:20 0.5 µl/well | n/a |
| AAV1.Syn.NES-jRGECO1a.WPRE.SV40 | 1 | hSyn | Gift from Adesnik lab | $2.08 \times 10^{13}$ | 1:20 0.5 µl/well | n/a |
| AAV9-U6-shRptor-EYFP | 9 | hU6 | Caltech CLOVER Center | $1.93 \times 10^{14}$ | 1:20 0.5 µl/well | 1:4 500 nl/mouse |
| AAV9-U6-shContorl-EYFP | 9 | hU6 | Caltech CLOVER Center | $2.17 \times 10^{14}$ | 1:20 0.5 µl/well | 1:4 500 nl/mouse |

**Supplementary Table 6. Antibodies and dilutions.**

| | Antibody | Host species | Company and catalog # | WB dilution | IHC dilution | ICC dilution |
|---|---|---|---|---|---|---|
| Primary | Tsc1 | Rabbit | Cell Signaling 6935 | 1:800 | - | - |
| | Raptor | Rabbit | Cell Signaling 2280 | 1:800 | - | - |
| | Rictor | Rabbit | Cell Signaling 2114 | 1:600 | - | - |
| | rpS6 | Rabbit | Cell Signaling 2317 | 1:1000 | - | - |
| | phospho-rpS6 Ser244/246 | Rabbit | Cell Signaling 5364 | 1:2000 | 1:800 | - |
| | Akt | Rabbit | Cell Signaling 4691 | 1:1500 | - | - |
| | phospho-Akt Ser 473 | Rabbit | Cell Signaling 4060 | 1:1000 | - | - |
| | 4E-BP1 | rabbit | Cell Signaling 9644 | 1:1000 | - | - |
| | phospho-4E-BP1 Thr37/46 | Rabbit | Cell Signaling 2855 | 1:1000 | - | - |
| | Histone 3 | Mouse | Cell Signaling 3638 | 1:2000 | - | - |
| | GFP | Chicken | AbCam ab13970 | - | 1:1000 | 1: 5000 |
| | MBP | Rat | Abcam ab7349 | - | 1: 350 | - |
| | GFAP | Rabbit | Fisher 180063 | - | 1: 400 | - |
| | GFAP | Mouse | Cell Signaling 3670 | - | 1: 400 | - |
| | NeuN | Mouse | Millipore MAB377 | - | 1: 800 | - |
| | **Antibody** | **Species (reactivity)** | **Company and catalog #** | **WB dilution** | **IHC dilution** | **ICC dilution** |
| Secondary | Goat anti-Rabbit-HFP | Rabbit | Bio-Rad 170-5046 | 1:5000 | - | - |
| | Goat anti-Mouse-HRP | Mouse | Bio-Rad 170-5047 | 1:5000 | - | - |
| | Goat anti-Rat Alexa Fluor 488 | Rat | Thermo Fisher A-11006 | - | 1:500 | - |
| | Goat anti-Chicken Alexa Fluor 488 | Chicken | Thermo Fisher A-11039 | - | 1:500 | 1:500 |
| | Goat anti-Mouse Alexa Fluor 546 | Mouse | Thermo Fisher A-11003 | - | 1:500 | - |

| | | | | | | |
|---|---|---|---|---|---|---|
| | Goat anti-Rabbit Alexa Fluor 633 | Rabbit | Thermo Fisher A-21070 | - | 1:500 | - |
